# Two parallel lineage-committed progenitors contribute to the developing brain

**DOI:** 10.1101/2025.07.02.662771

**Authors:** Carolyn E. Dundes, Rayyan T. Jokhai, Hadia Ahsan, Rachel S. Kang, Rachel E.A. Salomon-Shulman, Arjun Rajan, Yoon Seok Kim, Liam J. Stanton, Christine Xu, Stephanie Do, Brennan D. McDonald, José Miguel Andrade López, Hugo A. Urrutia, Hannah Greenfeld, Alicia Wong, Yimiao Qu, Andrew S. Petkovic, Yi Miao, K. Christopher Garcia, Michelle Monje, Daniel E. Wagner, Marianne E. Bronner, Christopher J. Lowe, Kyle M. Loh

## Abstract

The hindbrain is a life-sustaining brain region. In one model, a common neural progenitor generates all brain regions. Here our studies of mouse embryos and human pluripotent stem cells (hPSCs) support a different model: two parallel brain progenitors emerge simultaneously during gastrulation, anterior neural ectoderm (forebrain/midbrain progenitor) and posterior neural ectoderm (hindbrain progenitor). Not only are they lineage-committed to respectively form forebrain/midbrain vs. hindbrain *in vitro*, but they also have diverging chromatin landscapes foreshadowing future forebrain/midbrain vs. hindbrain identities. Leveraging these differences, we differentiated hPSCs into hindbrain rhombomere 5/6-specific motor neurons, hitherto difficult to generate *in vitro*. We postulate the brain is a composite organ emanating from two lineage-restricted progenitors; these dual progenitors may be evolutionarily conserved across 550 million years from hemichordates to mammals.

## Introduction

Stem cell research strives to create diverse types of human brain cell. Given plentiful successes in converting human pluripotent stem cells (hPSCs) into forebrain and midbrain cells^1-3^, here we focus on generating cells of the human hindbrain, a life-sustaining brain region^4-6^. Whereas the forebrain executes higher-level cognitive processes, the hindbrain constitutes most of the brainstem and coordinates life-sustaining processes such as breathing, eating, sleep, and wakefulness^4-6^. Consequently, hindbrain injury and diffuse intrinsic pontine glioma (a childhood hindbrain cancer) are both deadly, as they impair consciousness, sensation, reflexes, and breathing^7,8^. Additionally, hindbrain motor neurons project through the cranial nerves to innervate face and neck muscles, and degeneration of hindbrain motor neurons likely compromises eating and swallowing in diseases such as spinal muscular atrophy (SMA) and amyotrophic lateral sclerosis (ALS), leading to choking, aspiration, pneumonia, and in some cases, death^9-11^. However, differentiation of hPSCs into specific brain cells that correspond to different brain regions—such as the forebrain, midbrain, and hindbrain—is predicated on a developmental roadmap of when and how brain progenitors diversify from one another^4-6,12-15^.

During gastrulation, pluripotent cells form the ectoderm germ layer in the 7-day-old (∼E7.0) mouse embryo^16^. Subsequently, ectoderm forms neural, border, and surface ectoderm by E7.5, which will respectively create the brain, neural crest, and epidermis^16,17^. Arising during gastrulation, neural ectoderm represents the earliest brain-restricted progenitor^13,16^ and it subsequently differentiates into the forebrain, midbrain, and hindbrain by ∼E8.5^18,19^.

An open question is whether, even during gastrulation, neural ectoderm cells are diverse and are already committed to generate different brain regions. In 1952, Nieuwkoop^20,21^ proposed that neural ectoderm harbors the broad potential to generate the forebrain, midbrain, and hindbrain^22-25^. Indeed, the forebrain, midbrain, and hindbrain are anatomically contiguous, consistent with a scenario wherein they originate from a common brain precursor. Certain methods to differentiate hPSCs into brain cell-types implicitly assume the existence of a common neural progenitor that can generate the forebrain, midbrain, and hindbrain^22,23^. By contrast, classical fate maps of zebrafish^26^, frog^27^, chicken^28^, and mouse^29,30^ embryos have instead suggested that during gastrulation, neural ectoderm cells are already *fated* to give rise to distinct parts of the brain. However, these fate maps could not exclude the common neural progenitor model, because they generally focused on the fates of neural ectoderm cells if left in their native signaling environment in the gastrulating embryo^26-30^. One possible interpretation of these fate maps is that neural ectoderm cells have the potential to generate all brain regions, but owing to their position, later in development they are fated to encounter anterior- or posterior-inducing signals that only reveal part of their full lineage potential. An open question is whether distinct neural ectoderm cells are already *committed*: i.e., if challenged by ectopic extracellular signals, can they generate brain regions that they normally would not form? Assessing lineage commitment rests on the ability to challenge neural ectoderm cells with various signals, which we attempt here.

Inspired by pioneering embryological studies^12,13,31-33^, the current principal strategy to differentiate hPSCs into neural cells entails inhibition of BMP, TGFβ, and WNT *in vitro*^34-38^. This breakthrough has enabled many successes in generating human forebrain and midbrain cells from hPSCs^34-37,39-57^. Other work has focused on differentiating hPSCs into hindbrain^43,46,58-69^, but we do not fully understand the earliest hindbrain progenitors and the extracellular signals that induce them. To this end, we revisited the first ectodermal precursors in the mouse embryo. As a complementary approach, we also modeled ectoderm differentiation from hPSCs, with the proviso that *in vitro* models may not recreate the full complexity of *in vivo* development.

### Neural and non-neural ectoderm formation

During gastrulation, pluripotent cells first differentiate into either primitive streak or the definitive ectoderm germ layer^16^ (**Fig. 1a**). *Sox2*+ *Nanog*- *T*- definitive ectoderm^16,70-72^ arises by E6.75 in the mouse embryo, as shown by multiplexed *in situ* staining (**Fig. 1b**) and single-cell RNA-sequencing (scRNAseq, **Fig. S1a**). While BMP, FGF, TGFβ and WNT activation induces primitive streak^73,74^, conversely simultaneous BMP, FGF/ERK, TGFβ and WNT inhibition^34-37,39,75,76^ instead generated 96.5±3.4% pure SOX2+ NANOG-definitive ectoderm within 24 hours of hPSC differentiation (**Fig. 1c**, **Fig. S1b-h**). In addition to inhibiting BMP and TGFβ, the suppression of FGF/ERK and WNT signaling further downregulated *NANOG* and drove exit from pluripotency towards ectoderm (**Fig. S1b,d,h**). hPSC-derived definitive ectoderm appeared to be ectoderm-committed, and failed to respond to endoderm^77^- or paraxial mesoderm^78^-inducing signals (**Fig. S1i,j**), thereby providing a foundation to subsequently generate ectodermal derivatives.

**Figure 1:**
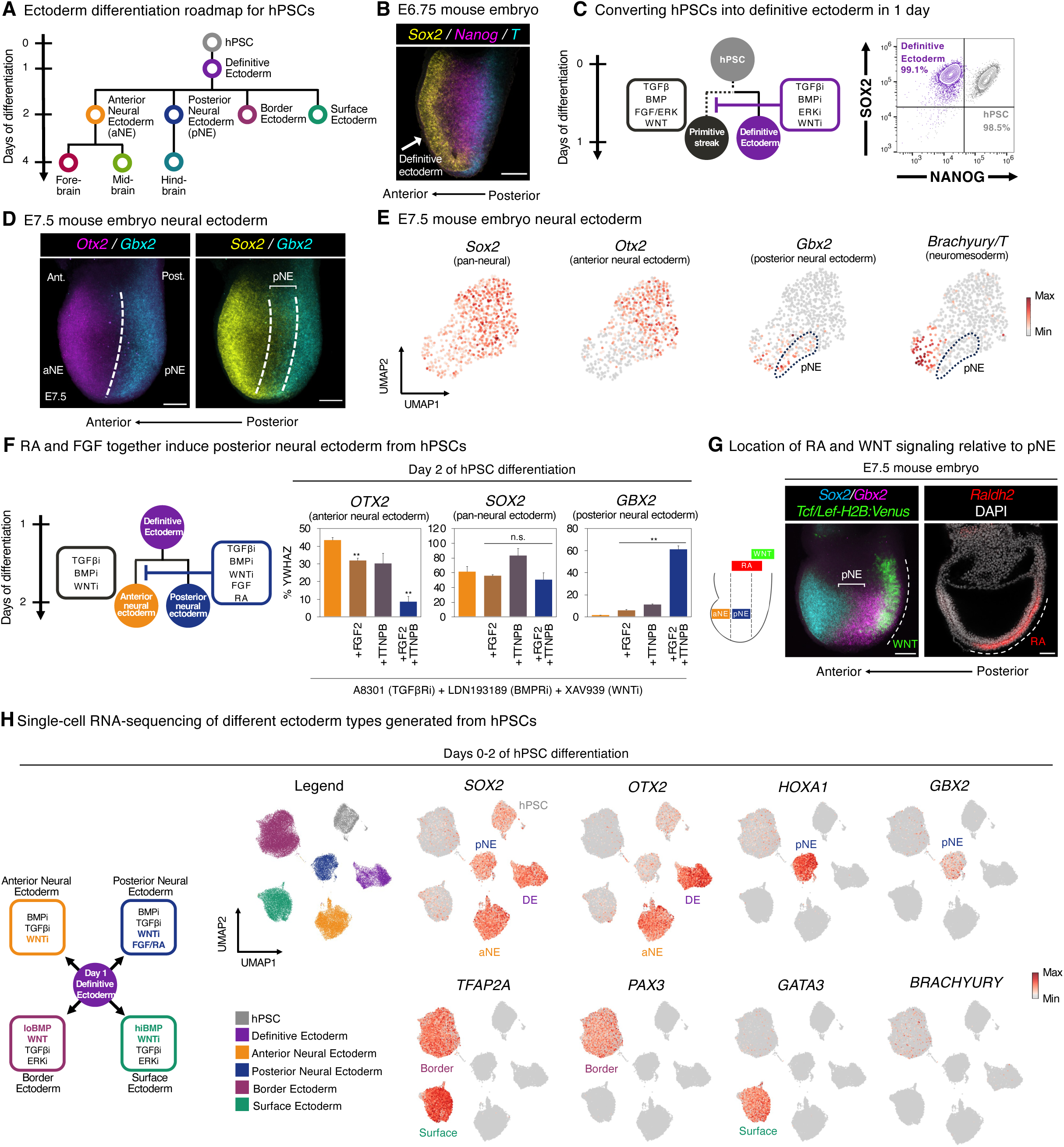
Anterior and posterior neural ectoderm *in vivo* and *in vitro*. A) hPSC differentiation strategy. B) *In situ* staining of E6.75 mouse embryo. Scale: 50 μm. Ant.: anterior. Post.: posterior. C) Flow cytometry of H1 hPSCs prior to, and after, definitive ectoderm differentiation for 24 hours. i: inhibitor. D) *In situ* staining of E7.5 mouse embryo. Scale: 50 μm. aNE: anterior neural ectoderm. pNE: posterior neural ectoderm. E) Analysis of published E7.5 mouse embryo single-cell RNA-sequencing data ^169^. Neural ectoderm cells are shown. F) Quantitative PCR (qPCR) of H1 hPSCs differentiated into definitive ectoderm for 24 hours, and then treated with neural-inducing signals (A-83-01 [1 μM], LDN193189 [250 nM], and XAV939 [1 μM]), in the presence or absence of FGF2 (20 ng/mL) and/or retinoid agonist TTNPB (50 nM), for 24 hours. Gene expression is shown relative to reference gene *YWHAZ*; 100% indicates that a gene is expressed at the same level as *YWHAZ*. G) *In situ* staining of the E7.5 mouse embryo. First, *in situ* staining for *Sox2* and *Gbx2* in a *Tcf/Lef:H2B:Venus* reporter mouse embryo ^173^ shows that *Gbx2*+ *Sox2*+ posterior neural ectoderm is largely devoid of WNT signaling; rather WNT signaling is active in the posterior embryo, where presumptive spinal cord progenitors are located ^25,92-102^ (*left*). A maximum intensity projection of lateral optical sections of a *Tcf/Lef:H2B:Venus* mouse embryo is shown. Second, *in situ* staining shows that *Raldh2/Aldh1a2* (the rate-limiting enzyme in retinoic acid [RA] synthesis) is expressed adjacent to posterior neural ectoderm (*right*), consistent with the notion that posterior neural ectoderm is exposed to RA *in vivo*. Scale bar = 50 μm. H) Single-cell RNA-sequencing of H7 hPSCs differentiated into day-1 definitive ectoderm, day-2 aNE, day-2 pNE, day-2 border ectoderm, or day-2 surface ectoderm. Colors denote differentiation conditions. qPCR data depicts the mean of two biological replicates, with s.e.m. shown. P values were calculated using a t-test. For flow cytometry and *in vitro* staining, a representative image from two biological replicates is shown. *In vitro* scRNAseq data is from a single experiment.

How do definitive ectoderm cells subsequently adopt neural vs. non-neural ectoderm identities^16^? Within E7.5 mouse embryos, *Sox2*+ neural ectoderm contains two mutually-exclusive *Otx2*+ vs. *Gbx2*+ populations^19,79-81^, as shown by multiplexed *in situ* staining (**Fig. 1d**, **Fig. S1k**) and scRNAseq (**Fig. 1e**) to confirm overlap of neural ectoderm marker *Sox2* with these two other genes. We respectively designate these cell-types as *Sox2*+ *Otx2*+ “anterior neural ectoderm” (aNE) and *Sox2*+ *Gbx2*+ “posterior neural ectoderm” (pNE). The locations of anterior and posterior neural ectoderm in the E7.5 mouse embryo generally comport with the positions of cells fated to respectively generate future forebrain/midbrain vs. hindbrain in classical fate maps^29^. These E7.5 cells represented early neural ectoderm, as they did not express markers of later-stage neural progenitors (*Sox1* and *Pax6*)^72,82^; moreover, posterior neural ectoderm expressed additional posterior markers *Hoxa1* and *Hoxb1*^81,83^ (**Fig. S1l**). Here we define anterior and posterior neural ectoderm as neural ectoderm that arises during gastrulation; these are distinct from subsequent forebrain, midbrain, and hindbrain neural progenitors that emerge after gastrulation.

What signals induce anterior vs. posterior neural ectoderm *in vitro*? Treating day 1 hPSC-derived definitive ectoderm with prevailing neural induction signals (combined BMP, TGFβ, and WNT inhibition^34-37,39,75^) for 24 hours generated OTX2+ GBX2-anterior neural ectoderm *in vitro* (**Fig. 1f**, **Fig. S1m,n**, **Fig. S2a**). This thus raised the question of what extracellular signals specify GBX2+ posterior neural ectoderm. We found that FGF and RA activation, together with BMP, TGFβ, and WNT blockade, for 24 hours specified GBX2+ OTX2^low^ posterior neural ectoderm (**Fig. 1f**). FGF and RA were combinatorially required to induce *GBX2*+ posterior neural ectoderm and to repress *OTX2*: either signal alone was insufficient (**Fig. 1f**, **Fig. S2b-d**). This thus extends the important roles of FGF and RA in inducing *Gbx2* and repressing *Otx2* in various model organisms^79,84-88^ to human neural ectoderm, and is consistent with how FGF and RA pathway genes (e.g., *Fgf4*, *Fgf8*, and *Raldh2*) are expressed nearby the E7.5 *Gbx2*+ posterior neural ectoderm *in vivo*^89-91^ (**Fig. 1g**). The requirement for WNT inhibition to induce posterior neural ectoderm induction *in vitro* (**Fig. S2e-i**, **Fig. S3a**) is consistent with the apparent lack of WNT activity in posterior neural ectoderm *in vivo* (**Fig. 1g**). Instead, WNT signaling was activated in the posterior-most region of the E7.5 mouse embryo (**Fig. 1g**), which is where *Brachyury* or *Cdx2*—markers of spinal cord or neuromesodermal progenitors—are expressed^92-102^. FGF and RA induced a different type of posterior character than WNT activation, the latter of which has been previously used to differentiate hPSCs into posterior neural cells^43,46,58-60,62,63,66,67,103^ (**Fig. S2e,g**; **Fig. S3a**). In particular, posterior neural ectoderm—induced by FGF and RA activation, alongside WNT inhibition—did not express *BRACHYURY* or *CDX2 in vivo* or *in vitro* (**Fig. 1e,h**; **Fig. S3a**); these are markers of spinal cord or neuromesodermal progenitors that are known to be induced by WNT^92-102^. Taken together, anterior vs. posterior neural ectoderm arise within 2 days of hPSC differentiation, and we identify the diverging extracellular signals that specify them.

In parallel, we could also differentiate hPSC-derived definitive ectoderm into two types of non-neural ectoderm within 24 hours: PAX3+ TFAP2A^low^ border ectoderm (by WNT activation, alongside low BMP^35,104-107^) and GATA3+ TFAP2A^high^ surface ectoderm (by WNT inhibition, alongside high BMP^35^); both border and surface ectoderm induction also required TGFβ and ERK blockade (**Fig. 1h, Fig. S3b-g**). This is consistent with how BMP gradients pattern ectoderm into border vs. surface domains *in vivo*^108,109^; we further find that WNT instructs border fate and blocks surface markers (**Fig. S3b,c,e**). In summary, we generated neural (anterior and posterior neural) and non-neural (border and surface) ectoderm within 2 days of hPSC differentiation, across 4 independent hPSC lines (**Fig. 1h**, **Fig. S4a-d**). These two types of neural ectoderm already express anterior or posterior markers, implying early acquisition of region-specific neural identities, which we test below.

### Neural ectoderm lineage tracing *in vivo*

To test the hypothesis that early neural ectoderm cells are already fated to generate either forebrain/midbrain vs. hindbrain *in vivo*^26-30^, we used two complementary genetic lineage-tracing approaches.

First, to test whether during gastrulation, *Gbx2*+ posterior neural ectoderm is already fated to form hindbrain, we crossed a *Gbx2-CreER* driver^110^ to a Cre-dependent *tdTomato* reporter^111^. Past *Gbx2-CreER* lineage-tracing^30^ suggested that *Gbx2*+ cells are posteriorly fated, but exploited tamoxifen, which perdures several days *in vivo*^112,113^ and could thus label *Gbx2*+ hindbrain cells that arise after gastrulation^80,81^. To acutely label E7.5 *Gbx2*+ posterior neural ectoderm, at E7.5 we injected the short-lived tamoxifen metabolite 4-hydroxytamoxifen (4OHT) to achieve a 12-hour labeling window^114,115^ (∼E7.5-E8.0). Posterior neural ectoderm-derived tdTomato^+^ progeny were restricted to the E8.5 and E9.5 hindbrain, and were not present in the *Otx2*^+^ forebrain or midbrain (**Fig. 2a**). At E18.5, posterior neural ectoderm-derived progeny were still restricted to the hindbrain, and were essentially absent from the forebrain and midbrain; we determined the mid/hindbrain boundary by the limit of hindbrain serotonergic neurons (**Fig. 2a**). Similar results were obtained after 4OHT administration at E7.0 (which would be expected to label from ∼E7.0-E7.5) to mark *Gbx2*+ posterior neural ectoderm incipiently emerging at E7.5; in these conditions, posterior neural ectoderm likewise exclusively contributed to the hindbrain, but not the forebrain/midbrain (**Fig. S4e,f**).

**Figure 2:**
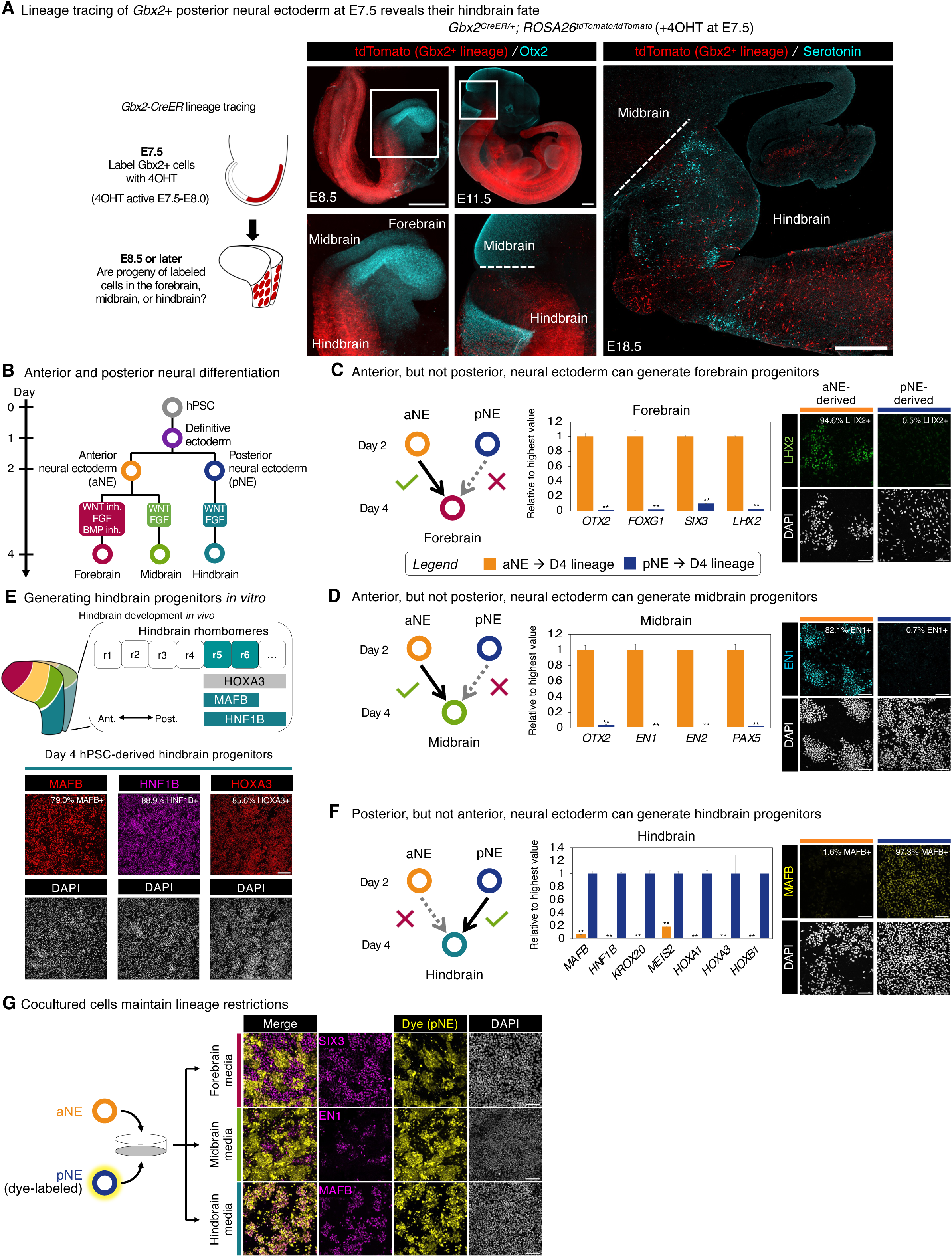
*In vivo* lineage tracing, and *in vitro* lineage commitment assays, of anterior and posterior neural ectoderm. A) E7.5 *Gbx2^CreER^*; *ROSA26^tdTomato^* mouse embryos were labeled with 4-hydroxytamoxifen (4OHT) to induce tdTomato expression in pNE. Subsequently, E8.5-E18.5 embryos were stained to visualize *Gbx2*^+^ pNE-derived progeny. The forebrain and midbrain were delimited by *Otx2 in situ* staining (for E8.5) or Otx2 immunostaining (for E11.5). At E18.5, the hindbrain was delimited by immunostaining for serotoninergic neurons present in the hindbrain, but not the midbrain. Scale: 100 μm (for E8.5 and E9.5) and 500 μm (for E11.5 and E18.5). B) hPSC differentiation strategy. C) qPCR (*left*) and immunostaining (*right*) of H1 hPSC-derived day-2 aNE or pNE treated with forebrain-inducing signals for 48 hours. Scale: 120 μm. Gene expression is shown relative to the sample with the highest expression. D) qPCR (*left*) and immunostaining (*right*) of H1 hPSC-derived day-2 aNE or pNE treated with midbrain-inducing signals for 48 hours. Scale: 120 μm. Gene expression is shown relative to the sample with the highest expression. E) Hindbrain gene expression *in vivo* ^4-6^ (*top*). Immunostaining of H1 hPSCs differentiated into definitive ectoderm for 24 hours, followed by posterior neural ectoderm for 24 hours, and hindbrain progenitors for 48 hours (*bottom*). Scale: 250 μm. F) qPCR (*left*) and immunostaining (*right*) of H1 hPSC-derived day-2 aNE or pNE treated with hindbrain-inducing signals for 48 hours. Scale: 120 μm. Gene expression is shown relative to the sample with the highest expression. G) H1 hPSCs were differentiated into either aNE or pNE within 2 days; pNE was fluorescently labeled, and then aNE (uncolored) and pNE (dye-labeled) were mixed. Cocultures were treated with either forebrain-, midbrain-, or hindbrain-inducing signals for 2 additional days, prior to immunostaining. Scale: 100 μm. qPCR data depicts the mean of two biological replicates, with s.e.m. shown. P values were calculated using a t-test. For *in vitro* staining, a representative image from two biological replicates is shown.

Second, we sparsely labeled *Sox2*+ neural ectoderm cells within the E7.5 mouse embryo (**Fig. S4g,h**). To this end, we crossed a *Sox2-CreER* driver^116^ to a Cre-dependent multicolor Confetti fluorescent reporter^117^, followed by delivery of a low 4OHT dose to sparsely label E7.5 neural ectoderm cells with one of three detectable colors (red, yellow, or cyan). An advantage of this approach is that it assesses the fate of any *Sox2*+ neural ectoderm progenitor, without regard to whether it already express anterior (*Otx2*) or posterior (*Gbx2*) markers. Analysis of 494 neural ectoderm-derived cell clusters spanning 16 independent embryos revealed each cell cluster was almost exclusively present in either the forebrain/midbrain (62.96%) or hindbrain (32.59%) (**Fig. S4g,h**). Therefore, by the end of gastrulation, most *Sox2*+ neural ectoderm cells are already fated to become either fore/midbrain or hindbrain.

Lineage tracing reveals the natural fate of progenitors in their native environment, but not their developmental potential ^118^ (i.e., the full range of fates they could adopt if challenged by ectopic signals). It is difficult to assess cellular potential in mammalian embryos *in utero*. Consequently, we turned to an *in vitro* model, whereby hPSC-derived anterior and posterior neural ectoderm were challenged with extracellular signals to test whether they were already committed to form specific brain regions, or whether they retained the potential to adopt alternative fates *in vitro*.

### Studies of lineage commitment *in vitro*

We developed a system to differentiate hPSCs into forebrain, midbrain, and hindbrain progenitors (**Fig. 2b**). Day 2 anterior neural ectoderm could further bifurcate into *FOXG1*+ *SIX3*+ forebrain progenitors (by blocking posteriorizing signals WNT and BMP, while activating FGF; **Fig. 2c, Fig. S5a-d**) or *EN1*+ midbrain progenitors (by activating WNT and FGF; **Fig. 2d**, **Fig. S5e-i**) within 2 days of further differentiation. This is congruent with the posterior WNT gradient across the developing forebrain and midbrain *in vivo* (**Fig. S5j**) and how WNT establishes midbrain identity *in vivo*^119-121^ and *in vitro*^34-36,39-41,43-47,56^.

Along the other developmental path, day 2 posterior neural ectoderm could further differentiate into hindbrain progenitors upon combined FGF and WNT activation for 2 days (**Fig. 2e,f**; **Fig. S6a-f**). The developing hindbrain is divided into six segments known as rhombomeres^4-6^, and hPSC-derived hindbrain progenitors expressed MAFB, HNF1B, and HOXA3, which together identify rhombomeres 5 and 6 *in vivo*^6,122,123^ (**Fig. 2e**). Indeed, FGF is likewise required for rhombomere 5 and 6 specification *in vivo*^124,125^. Together, these results reveal that FGF and WNT specify human hindbrain identity, building on observations in model organisms^103,126-128^. To our knowledge, this is the first demonstration that human rhombomere 5 and 6 hindbrain progenitors can be generated *in vitro*, thereby complementing past work differentiating hPSCs into other types of hindbrain progenitor^43,46,58-69^.

Day 2 hPSC-derived anterior vs. posterior neural ectoderm were respectively committed to generating forebrain/midbrain or hindbrain progenitors *in vitro*. Strikingly, when posterior neural ectoderm was treated with forebrain- or midbrain-inducing signals, it largely failed to acquire either of these identities by day 4 (**Fig. 2c,d**, **Fig. S6g**). Reciprocally, when anterior neural ectoderm was challenged with hindbrain-inducing conditions, it failed to differentiate into hindbrain by day 4 (**Fig. 2f, Fig. S6g**, **Fig. S7a**). Thus, in the *in vitro* conditions tested, anterior and posterior neural ectoderm are both lineage committed. However, it remains formally possible that there may be other experimental interventions that could enable them to interconvert.

Another consideration is that the community effect^129^ could affect the interpretation of these experiments, and we thus tested if hPSC-derived anterior and posterior neural ectoderm remained lineage-committed even when mixed together. hPSC-derived unlabeled anterior neural ectoderm was mixed with fluorescently-labeled posterior neural ectoderm, and these cocultures were then challenged with forebrain-, midbrain-, or hindbrain-inducing signals (**Fig. 2g**, **Fig. S7b**). Within these cocultures, anterior neural ectoderm generated SIX3+ forebrain and EN1^+^ midbrain progenitors, but not MAFB+ hindbrain progenitors, and *vice versa* for posterior neural ectoderm, although a few posterior neural ectoderm cells expressed SIX3 in these cocultures (**Fig. 2g**, **Fig. S7b**). Therefore, even when cocultured with one another, anterior and posterior neural ectoderm generally obey their respective lineage commitments.

### Divergent chromatin landscapes

Anterior and posterior neural ectoderm harbored different accessible chromatin landscapes (**Fig. 3a**, **Fig. S7c**, **Table S1**), foreshadowing their potentials to develop into differing downstream brain lineages. At the genome-wide level, anterior neural ectoderm-enriched accessible chromatin was enriched for OTX2 motifs (**Fig. 3b**, **Table S2**). By contrast, posterior neural ectoderm-enriched accessible chromatin was instead enriched for the motifs of future hindbrain transcription factors (HOX, HNF1B, and MAFB), and the motifs of FGF (ETV1) and RA (RARB and RXRG) pathway transcriptional effectors (**Fig. 3b**, **Table S2**). This is consistent with how FGF and RA induce posterior neural ectoderm and have key roles in later hindbrain development^4-6^.

**Figure 3:**
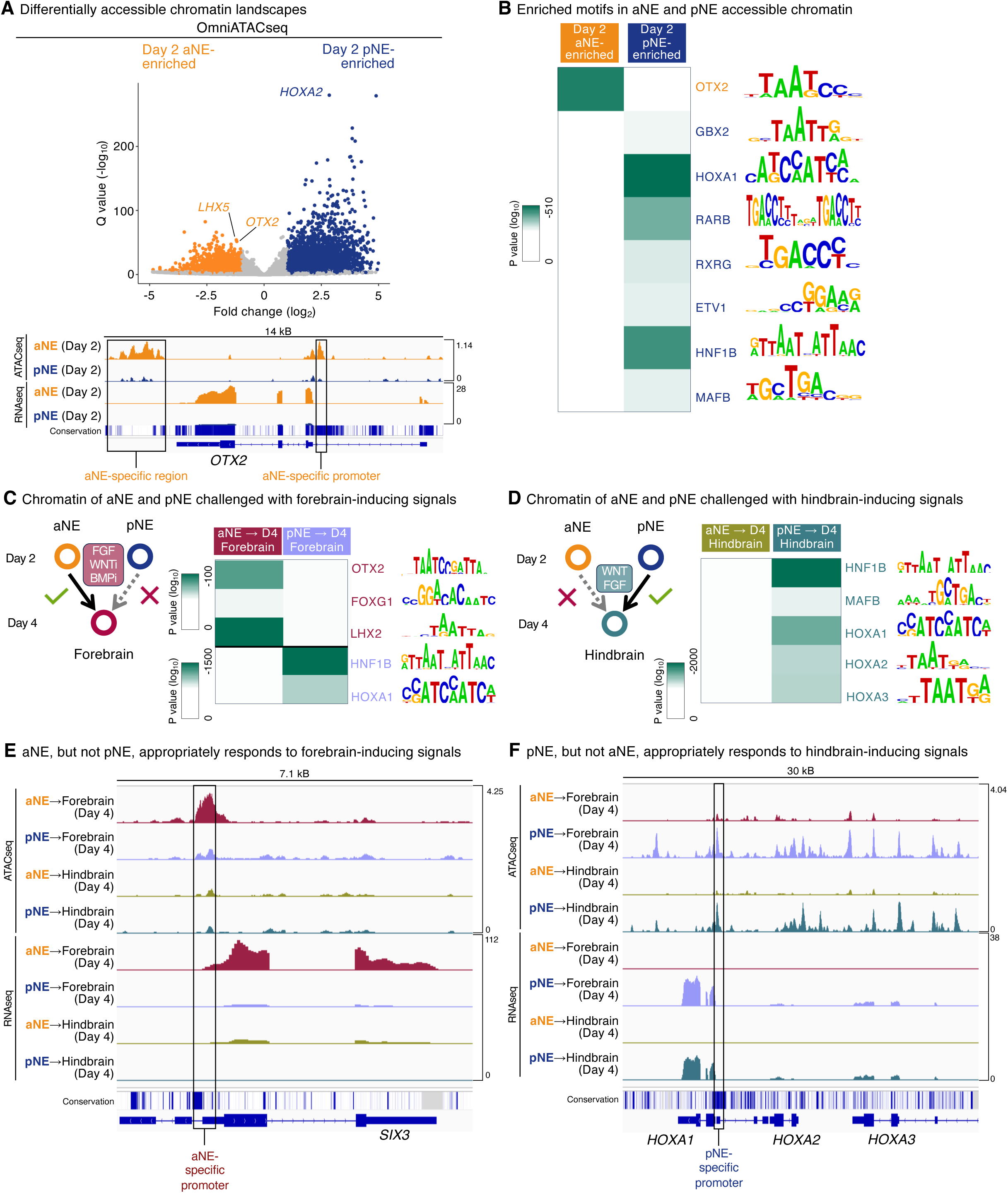
Different accessible chromatin landscapes distinguish hPSC-derived anterior vs. posterior neural ectoderm. A) Differentially accessible chromatin regions in hPSC-derived day-2 aNE or pNE, as detected by OmniATACseq. Top: each dot indicates a single genomic locus, which was assigned to its nearest gene. Bottom: OmniATACseq and bulk-population RNAseq of hPSC-derived aNE or pNE. Conservation: Phastcons evolutionary conservation of genome sequence across 46 vertebrate species. B) Transcription factor motifs that were respectively enriched in either day-2 aNE- or pNE- specific accessible chromatin regions. C) hPSC-derived aNE and pNE were treated with forebrain-inducing signals for 2 days, followed by OmniATACseq to identify chromatin regions that were preferentially accessible in either population. Transcription factor motifs enriched in the accessible chromatin of either aNE or pNE derivatives are shown. D) OmniATACseq and bulk-population RNAseq of hPSC-derived aNE and pNE that were treated with either forebrain- or hindbrain-inducing signals for 2 days. E) hPSC-derived aNE and pNE were treated with hindbrain-inducing signals for 2 days, followed by OmniATACseq to identify chromatin regions that were preferentially accessible in either population. Transcription factor motifs enriched in the accessible chromatin of either aNE or pNE derivatives are shown. F) OmniATACseq and bulk-population RNAseq of hPSC-derived aNE and pNE that were treated with either forebrain- or hindbrain-inducing signals for 2 days. OmniATACseq and bulk-population RNAseq were each performed on two biological replicates. Q values for differential accessibility were calculated using the Wald test, followed by Benjamini-Hochberg adjustment. P values for motif enrichment were calculated using the Fisher exact test.

How do anterior and posterior neural ectoderm respond at the chromatin level when challenged with developmentally-inappropriate cues? Anterior neural ectoderm treated with forebrain-inducing signals appropriately increased chromatin accessibility at forebrain genes such as *SIX3*, and at the genome-wide level, their accessible chromatin landscape was enriched for key forebrain transcription factor motifs (OTX2, FOXG1, and LHX2; **Fig. 3c,e**, **Fig. S7d**, **Table S2**). Conversely, posterior neural ectoderm challenged with the same forebrain-inducing signals (WNT inhibition and FGF activation) instead adopted a hindbrain-like chromatin landscape. They failed to increase accessibility at forebrain genes (**Fig. 3e**), and instead increased accessibility at a swath of hindbrain regulatory elements enriched for hindbrain transcription factor motifs (HNF1B and HOX; **Fig. 3c**, **Fig. S7d,e**, **Table S2**). In sum, posterior neural ectoderm resists forebrain-inducing signals, and its chromatin landscape seems committed to subsequently adopt a hindbrain regulatory program.

Anterior neural ectoderm, even when confronted with hindbrain-inducing signals (WNT and FGF activation), instead adopted a forebrain-like regulatory program and failed to increase accessibility at hindbrain genes (e.g., the *HOXA* locus) or hindbrain transcription factor motifs (**Fig. 3d,f**, **Fig. S7e,f**, **Table S2**). On the other hand, posterior neural ectoderm treated with hindbrain-inducing signals activated a hindbrain regulatory program (**Fig. 3d,f**, **Fig. S7e,f**, **Table S2**). Taken together, anterior vs. posterior neural ectoderm have divergent transcriptional and chromatin landscapes that prefigure their respective potentials to subsequently develop into forebrain and hindbrain lineages (**Fig. S7g,h**). Though it typically takes weeks or months to generate neurons *in vitro*, we propose that cellular competence to generate forebrain/midbrain or hindbrain has already become restricted within the first 2 days of hPSC differentiation.

### Dorsal vs. ventral forebrain and hindbrain

After its formation, the forebrain undergoes dorsal-ventral patterning into the dorsal forebrain (neocortex, which primarily generates cortical glutamatergic neurons^3,15^) and ventral forebrain (which forms cortical GABAergic interneurons and certain hypothalamic neurons^130,131^) (**Fig. 4a**). Likewise, the hindbrain also undergoes dorsal-ventral patterning, forming dorsal and ventral hindbrain progenitors that respectively produce hindbrain sensory and motor neurons^132,133^ (**Fig. 4a**).

**Figure 4:**
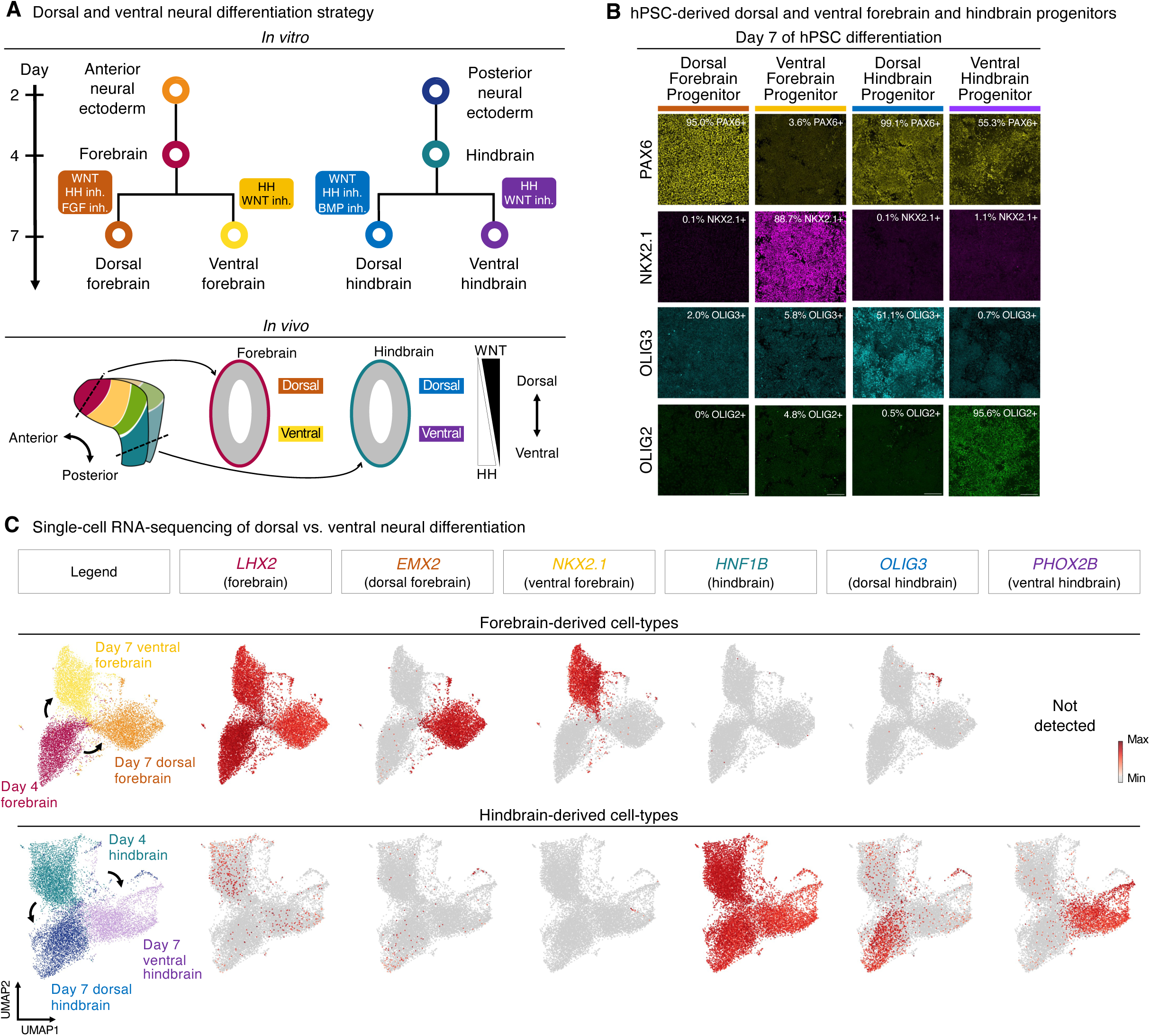
Generation of dorsal forebrain, ventral forebrain, dorsal hindbrain, and ventral hindbrain progenitors from hPSCs *in vitro*. A) Summary of hPSC differentiation *in vitro* (*top*) and brain development *in vivo* (*bottom*). B) Immunostaining of H1 hPSC-derived day-4 forebrain or hindbrain progenitors that were treated with dorsalizing or ventralizing signals for 72 hours. Scale: 120 μm. C) scRNAseq of H7 hPSC-derived day-4 forebrain or hindbrain progenitors, and day-7 dorsal forebrain, ventral forebrain, dorsal hindbrain, or ventral hindbrain progenitors. Colors denote differentiation conditions. For *in vitro* staining, a representative image from two biological replicates is shown. *In vitro* scRNAseq data is from a single experiment.

Starting from day 4 hPSC-derived forebrain progenitors, we found that WNT activation— combined with HH and FGF inhibition—specified PAX6+ EMX2+ dorsal forebrain progenitors (**Fig. 4b,c**, **Fig. S8a-c**). Conversely, HH activation^34^ together with WNT inhibition yielded NKX2.1+ ventral forebrain progenitors (**Fig. 4b,c**, **Fig. S8d**). Our *in vitro* results are consistent with how countervailing dorsal WNT vs. ventral HH gradients pattern the forebrain *in vivo*^134-137^. Notably, the requirement for WNT to specify dorsal forebrain agrees with *in vivo* studies^134-137^ and contrasts with prolonged WNT inhibitor treatment to specify hPSC-derived dorsal forebrain progenitors^34-37,39,40^.

Starting from day 4 hPSC-derived hindbrain progenitors, WNT activation together with HH and BMP inhibition generated OLIG3+ PAX3+ dorsal hindbrain progenitors^132,138^ (**Fig. 4b,c**, **Fig. S8e-h**). By contrast, application of the converse signals—WNT inhibition and HH activation— generated OLIG2+ NKX6.2+ NKX2.2+ ventral hindbrain progenitors^133,138^ (**Fig. 4b,c**, **Fig. S8g-i**). Generation of a given progenitor (e.g., ventral hindbrain) relied on activation of a lineage-instructive signal (e.g., HH), together with simultaneously blocking the signal that induced the opposite lineage (e.g., WNT).

Despite the similarity of the signals added to induce dorsal (WNT) vs. ventral (HH) patterning in forebrain and hindbrain cultures, dorsal and ventral forebrain progenitors minimally expressed hindbrain markers, and *vice versa* (**Fig. 4b,c**). Therefore, forebrain (anterior neural ectoderm-derived) and hindbrain (posterior neural ectoderm-derived) progenitors continued to maintain their respective identities during dorsal-ventral patterning. This likely reflects the deeply ingrained differences in anterior vs. posterior neural ectoderm laid down earlier during differentiation.

### Making human hindbrain motor neurons

These four different neural progenitors (day-7 hPSC-derived dorsal forebrain, ventral forebrain, dorsal hindbrain and ventral hindbrain) could further differentiate into mutually-exclusive types of neuron (**Fig. 5a**). To this end, we provided neurotrophic factors (BDNF, GDNF, Forskolin and Vitamin C) while blocking NOTCH (to drive neural progenitors out of the cell cycle and into postmitotic neurons) for 7 additional days^139^, generating *MAPT*+ *SNAP25*+ neurons alongside some remaining undifferentiated *SOX2*+ neural progenitors (**Fig. 5a**, **Fig. S9a**). Dorsal forebrain progenitors exposed to these neuron-inducing signals differentiated into *TBR1*+ cortical glutamatergic Cajal-Retzius neurons (**Fig. 5b,c**, **Fig. S9a,b**). Conversely, ventral forebrain progenitors instead formed *DLX1*+ cortical interneurons, in addition to hypothalamic-like *OTP*+ and *ISL1*+ neurons that respectively expressed the *ADCYAP1*^140,141^ and *POMC*^142^ neuropeptide genes (**Fig. 5b,c**, **Fig. S9a,c,d**). This could enable *in vitro* modeling of hunger-suppressing *POMC*+ hypothalamic neurons; their importance *in vivo* is underscored by the severe obesity of *POMC*-deficient humans and mice^142^.

**Figure 5:**
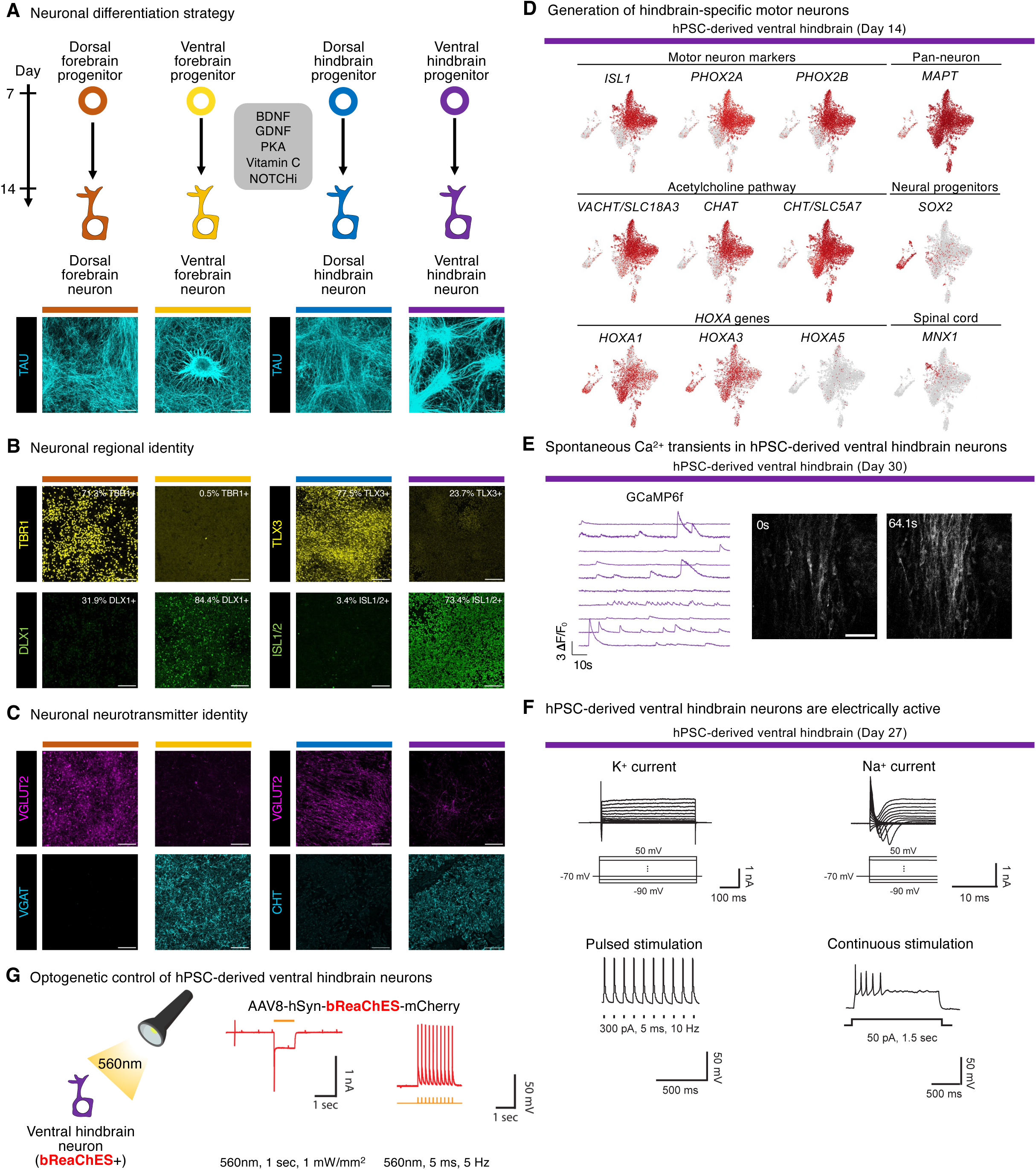
Generation of hindbrain motor neurons from hPSCs *in vitro*. A) Immunostaining of H1 hPSC-derived day-7 dorsal forebrain, ventral forebrain, dorsal hindbrain, or ventral hindbrain progenitors that were exposed to the same neuron-inducing signals for 7 additional days *in vitro*. Scale: 120 μm. B) Immunostaining of H1 hPSC-derived day-7 dorsal forebrain, ventral forebrain, dorsal hindbrain, or ventral hindbrain progenitors that were exposed to the same neuron-inducing signals for 7 additional days *in vitro*. Scale: 120 μm. C) Immunostaining of H1 hPSC-derived day-7 dorsal forebrain, ventral forebrain, dorsal hindbrain, or ventral hindbrain progenitors that were exposed to the same neuron-inducing signals for 7 additional days *in vitro*. Scale: 120 μm. D) scRNAseq of H7 hPSC-derived day-14 ventral hindbrain population. E) Live Ca^2+^ imaging of WTC11 *AAVS1-CAG-GCaMP6f* hPSCs differentiated into ventral hindbrain neurons for 30 days, which exhibited spontaneous Ca^2+^ transients as assessed within individual cells (*left*) and across the culture (*right*). Scale: 50 μm. F) Electrophysiological activity of H1 hPSC-derived ventral hindbrain neurons on day 27 of differentiation, showing voltage-dependent K^+^ or Na^+^ channel currents elicited by depolarization of the holding potential from -90 mV to 50 mV in voltage-clamp mode (*top*) and action potentials elicited by injection of pulsed (300 pA, 5 ms, 10 Hz) or prolonged (50 pA, 1.5 seconds) currents in current-clamp mode (*bottom*). G) Day 28 H1 hPSC-derived ventral hindbrain neurons transduced with AAV8-*hSyn-bReaChES-mCherry* were stimulated with either prolonged (1 second), or pulses of (5 ms, 5 Hz), 560nm light, and electrophysiological recording was performed to detect action potentials. For *in vitro* staining, a representative image from two biological replicates is shown. *In vitro* scRNAseq data is from a single experiment. Ca^2+^ imaging and electrophysiology were each performed on a single experiment.

In parallel, we differentiated hPSCs into hindbrain neurons. Hindbrain motor neurons project through the cranial nerves and secrete acetylcholine to control face and neck muscles^4-6^. While past work has differentiated hPSCs into certain types of hindbrain neuron^43,46,58-69^, it has remained challenging to generate human rhombomere 6-specific motor neurons, which project through cranial nerve IX to control muscles crucial for swallowing^5^. To meet this challenge, we differentiated hPSC-derived ventral hindbrain progenitors into hindbrain motor neurons that expressed hallmark transcription factor *ISL1*^143^ and acetylcholine pathway genes (*VACHT, CHAT* and *CHT*) (**Fig. 5b,c**, **Fig. S9a,e**). Cholinergic motor neurons reside in both the hindbrain and spinal cord, but our hPSC-derived hindbrain motor neurons expressed hallmarks of hindbrain rhombomere 5/6 identity (*HOXA1*, *HOXA3*, *PHOX2A*, and *PHOX2B*)^4-6,144^, while minimally expressing spinal cord markers (*HOXA5* and *MNX1/HB9*; **Fig. 5d**). These hindbrain motor neurons arose alongside smaller subsets of other ventral hindbrain neuron subtypes (**Fig. S9f**). In contrast, dorsal hindbrain progenitors exposed to the same neuron-inducing signals instead generated *TLX3*+ *LBX1*+ somatosensory-like neurons, which are crucial for sensory information reception and processing *in vivo*^132,145^ (**Fig. 5b**, **Fig. S9g**). In short, we generated hPSC-derived ventral and dorsal hindbrain neurons that continued to express rhombomere 5/6-specific markers.

hPSC-derived ventral hindbrain neurons were electrophysiologically active. They displayed spontaneous Ca^2+^ transients, as measured by GCaMP6f^146^ (**Fig. 5e**, **Fig. S10a-c**) and fired action potentials when injected with depolarizing currents (**Fig. 5f**) or optogenetically stimulated with the red-shifted excitatory opsin bReaChES^147^ (**Fig. 5g**, **Fig. S10d**). Similar results were observed with hPSC-derived dorsal hindbrain neurons (**Fig. S10e-g**). Posterior neural ectoderm thus provides a path to generate electrophysiologically-active hindbrain motor neurons *in vitro*.

### Evolutionary conservation

Finally, given that anterior and posterior neural ectoderm arise within the gastrulating mouse embryo, do similar progenitors exist in other species (**Fig. 6a**)? Building on past studies^28,85,148-150^, we found separate *Otx*+ *Sox2*+ anterior and *Gbx*+ *Sox2*+ posterior neural ectoderm populations in the gastrulating embryos of multiple vertebrate species, including macaque (**Fig. 6b**), chicken (**Fig. 6c**), and zebrafish (**Fig. 6d**). Multiplexed *in situ* staining of chicken and zebrafish embryos confirmed mutually-exclusive *Otx2*+ and *Gbx2*+ subsets of *Sox2*+ neural ectoderm cells (**Fig. 6c,d**). We additionally found that mutually-exclusive *otx*+ vs. *gbx*+ ectoderm arose during blastulation and gastrulation in a hemichordate lineage, the acorn worm *Saccoglossus kowalevskii*^151^ (**Fig. 6e**, **Fig. S10h**). Hemichordates are closely related to chordates, and shared a common ancestor ∼550-600 million years ago with the other deuterostome species examined here^152^. Our discovery of posterior ectoderm in gastrulating acorn worms is notable, as our prior work revealed a hindbrain-like molecular program in this non-chordate species^153^. Taken together, these results intimate that two parallel ectodermal progenitors might be deeply evolutionarily conserved across deuterostomes.

**Figure 6:**
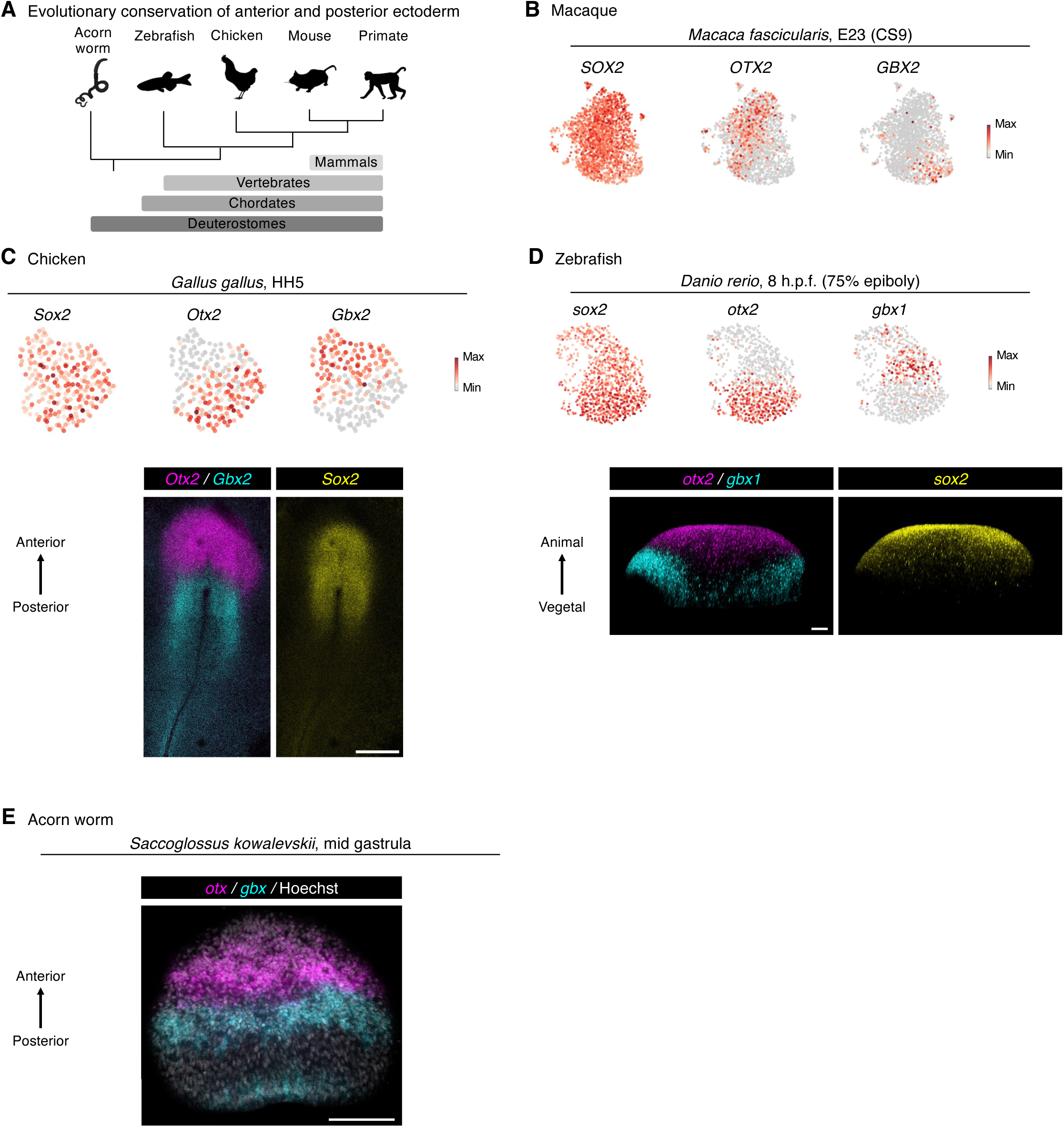
Evolutionary conservation of anterior and posterior ectoderm. A) Summary of evolutionary relationships. B) Single-cell RNA-sequencing of gastrulating macaque embryo (Carnegie Stage 9 [CS9]; embryonic day 23 [E23]); analysis performed on a previously-published resource ^170^. Neural ectoderm cells are shown. C) Single-cell RNA-sequencing of gastrulating chicken embryo (Hamburger-Hamilton stage 5 [HH5]); analysis performed on a previously-published resource ^171^. Neural ectoderm cells are shown (*top*). *In situ* staining of HH5 stage chicken embryo. Scale: 500 μm (*bottom*). D) Single-cell RNA-sequencing of gastrulating zebrafish embryo (8 hours post-fertilization [h.p.f.], 75% epiboly); analysis performed on a previously-published resource ^172^. Neural ectoderm cells are shown (*top*). *In situ* staining of 8 h.p.f. zebrafish embryo, scale bar: 50 μm (*bottom*). E) *In situ* staining of gastrulating acorn worm embryo. A maximum intensity projection of optical sections focusing on the ectoderm is shown. Scale: 100 μm. Staining was performed on a single embryo of each species.

## Discussion

At the crux of this work is the question of when forebrain/midbrain vs. hindbrain identities become separated during embryonic development. In one model, neural ectoderm represents a common progenitor for the entire brain^20-25^. By contrast, classical fate maps suggested that even as early as gastrulation, distinct neural ectoderm cells are already fated to form different brain regions^26-30^. However, these pioneering fate mapping studies did not test whether neural ectoderm cells are already committed to form specific brain regions (which requires the ability to isolate neural ectoderm cells and challenge them in various conditions) nor the genomic mechanisms underlying lineage restriction. Our data suggest that anterior and posterior neural ectoderm are already committed to form different brain regions *in vitro*. When challenged with hindbrain-inducing signals *in vitro*, hPSC-derived anterior neural ectoderm cannot “transdifferentiate” into hindbrain. Reciprocally, hPSC-derived posterior neural ectoderm cannot “transdifferentiate” into forebrain/midbrain. Mechanistically, hPSC-derived anterior and posterior neural ectoderm harbor divergent accessible chromatin landscapes, which foreshadow their different lineage potentials and furthermore resist inappropriate lineage-inducing signals.

More broadly speaking, our work relates to the longstanding question in developmental biology of how and when different regions of the central nervous system diversify from one another. Important studies have challenged the assumption that there is a common central nervous system progenitor, and have instead discovered separate progenitors to the brain vs. spinal cord (the latter called “neuromesoderm”)^25,92-102^. There are thus separate anterior (brain-specific) and posterior (spinal cord-specific) precursors that form the central nervous system^25,92-102^. We suggest further complexity: although the brain is often construed as a single organ, the brain may itself derive from two parallel progenitors that emerge early in development and that respectively form the forebrain/midbrain vs. the hindbrain. Analogously, the heart derives from two parallel progenitors, the first and second heart fields, which converge to form a single organ^154^. Taken together, the central nervous system may arise from the coalescence of multiple region-specific progenitors that separately give rise to the forebrain/midbrain (anterior neural ectoderm), hindbrain (posterior neural ectoderm), and spinal cord (neuromesoderm^25,92-102^). However, our results cannot exclude the possibility that a “pan-brain” progenitor exists for a very brief duration *in vivo*, between the emergence of the ectoderm germ layer (∼E6.75-E7.0)^16^ and the bifurcation of anterior vs. posterior neural ectoderm (∼E7.5).

Classical embryological studies generally support our model that anterior and posterior neural ectoderm represent parallel progenitors. Early forebrain and midbrain—which are both anterior neural ectoderm-derived lineages—can interconvert *in vivo* if confronted with various experimental conditions; however, in these studies, forebrain and midbrain could not apparently adopt hindbrain identity^155-158^. Therefore, anterior and posterior neural ectoderm cannot readily interconvert under normal conditions. However, earlier perturbations that may affect the initial emergence of anterior vs. posterior neural ectoderm during gastrulation—such as manipulating transcription factors that specify anterior vs. posterior neural ectoderm identity (e.g., *Otx2*)^159,160^ or deleting *Fgf8* (which is required for posterior neural ectoderm formation)^30,84^—alters the subsequent acquisition of midbrain vs. hindbrain identities *in vivo*. Perturbing early anterior and posterior neural ectoderm specification thus impacts the development of their downstream progeny. However, it remains to be determined whether anterior and posterior neural ectoderm are lineage committed *in vivo*; this is technically challenging, as it would entail transplanting purified, labeled populations of these progenitor cells into ectopic locations in the gastrulating mammalian embryo.

We find that the fundamental bifurcation between anterior vs. posterior neural ectoderm occurs surprisingly early, within the first 2 days of hPSC differentiation, long preceding the emergence of classically-defined neural progenitors or neurons. This parallels how these anterior vs. posterior neural ectoderm rapidly arise from pluripotent cells *in vivo* within 2 days, between E5.5 to E7.5. Influential studies previously demonstrated that hPSCs can be differentiated into two different types of anterior vs. posterior neural progenitor within 2 weeks *in vitro*, which express the classical neural progenitor markers PAX6 and SOX1^161,162^. However, the bifurcation of anterior vs. posterior neural ectoderm occurs earlier during development, taking place during gastrulation and preceding the expression of neural progenitor markers *Pax6* or *Sox1*^72,163^ at later developmental stages.

Defining and manipulating the early fundamental lineage decision between anterior vs. posterior neural ectoderm is paramount to precisely direct stem cells toward a hindbrain developmental path, in preference to a forebrain/midbrain lineage. If this early time window is missed, and differentiating stem cells proceed down the “wrong” developmental track, they cannot seem to readily crossover to adopt a different brain regional identity. This emphasizes the importance of the early bifurcation of anterior vs. posterior neural ectoderm, and reveals hitherto-cryptic diversity among the earliest human neural ectoderm cells *in vitro*.

Posterior neural ectoderm provides a platform to subsequently generate certain types of hindbrain neuron *in vitro*. We find that prevailing signals used to differentiate hPSCs towards neural fates (BMP, TGFβ, and WNT inhibition)^34-37,75,164-168^ generate anterior neural ectoderm, explaining the past success that the field has enjoyed in creating forebrain and midbrain neurons^1-3^. Conversely, we found that simultaneous FGF and RA activation alongside neural induction signals (BMP, TGFβ, and WNT inhibition) specified posterior neural ectoderm. We subsequently differentiated these precursors into hindbrain progenitors and hindbrain motor neurons corresponding to rhombomeres 5/6, which were electrophysiologically active and expressed defining hindbrain-specific transcription factors (*PHOX2A*, *PHOX2B* and anterior *HOX* genes)^4-6,144^ and acetylcholine pathway genes. This thus complements past work that successfully differentiated hPSCs into other types of hindbrain neuron^43,46,58-69^. In particular, the ability to create human rhombomere 5/6-specific motor neurons—which are crucial for swallowing *in vivo*—could provide a powerful platform to study SMA, ALS, and other neurodegenerative diseases wherein impaired swallowing can lead to choking and death^9-11^. More broadly, the hindbrain is essential for breathing, eating, sleep, wakefulness, and other life-critical functions^4-6^, and the ability to create human hindbrain neurons holds promise for modeling other deadly diseases that affect the hindbrain^7,8^.

Our results clearly show that recapitulating the initial bifurcation of anterior vs. posterior neural ectoderm is critical to differentiate hPSCs into neurons in a manner that reflects anterior-posterior regional identities (e.g., forebrain, midbrain, or hindbrain). Separate anterior and posterior ectoderm populations arise during gastrulation across deuterostome species as diverse as acorn worm, zebrafish, chicken, mouse, and primate, implying that this distinction between two different types of ectoderm predates the origins of chordates and arose ∼550-600 million years ago^152^. We conclude that the emerging notion of two parallel brain progenitors has ramifications for development, differentiation, and evolution.

## Acknowledgements

We thank James Li, Bruce Conklin, Teni Anbarchian, Roel Nusse, Fabian Suchy, Hiromitsu Nakauchi, Andrew Elefanty, Edouard Stanley, and Elizabeth Ng for sharing reagents. We also thank Siva Vijayakumar, Benjamin Woodruff, Chiara Anselmi, Ayelet Voskoboynik, and Andrew Petkovic for contributing to experiments, and William Talbot, Aaron Zorn, Kenneth Campbell, Robb Krumlauf, Philip Beachy, Carla Shatz, John Day, Alexander Pollen, Tomasz Nowakowski, Lay Teng Ang, and Nobuko Uchida for advice. Infrastructure support was provided by Valerie Park, Liying Ou, Kitty Lee, Catherine Carswell-Crumpton, Patricia Lovelace, Laura Dunkin-Hubby, and the Stanford Institute for Stem Cell Biology & Regenerative Medicine, Stem Cell FACS Core Facility, Veterinary Service Center, Cell Sciences Imaging Facility, and Diabetes Research Center (NIH P30DK116074). This work was supported by the NIH (DP5OD024558 [K.M.L.], DP2GM146258 [D.E.W.], R00GM121852 [D.E.W.], R01DK115728 [K.C.G.], R01DE027538 [M.E.B.], T32GM119995 [C.E.D.], T32GM007365 [R.T.J.], T32GM007790 [R.T.J.], and F31DE031154 [H.A.U.]), NSF (IOS1656628 [C.J.L.]), CIRM (TB1-01195 [R.E.A.S.], EDUC2-12677 [A.S.P.]), Spinal Muscular Atrophy Foundation (K.M.L.), Stanford MCHRI, Beckman and Ludwig Centers (K.M.L.), Siebel Stem Cell Institute (K.M.L.), Stinehart-Reed Seed Grant (K.M.L.), Anonymous, Fickel and Gilbert families (K.M.L.), Gatsby Charitable Foundation (M.M.), Howard Hughes Medical Institute (M.M. and K.C.G.), Ford Foundation, Stanford Graduate, and Stanford DARE Fellowships (C.E.D.), Stanford Medical Scholars Research Program (R.T.J.), Yale Saybrook College and Stacey Leondis Fellowships (C.X.), and Stanford MCHRI and Dean’s Postdoctoral Fellowships (Y.Q.). K.M.L. was supported as a Packard Foundation Fellow, Pew Scholar, Baxter Foundation Faculty Scholar, Human Frontier Science Program Young Investigator (RGY0069/2019) and Anthony DiGenova Endowed Faculty Scholar.

## Author contributions

C.E.D. performed lineage tracing. C.E.D., R.T.J., R.E.A.S., H.A., L.J.S., C.X., S.D., R.S.K., A.W., Y.Q., A.S.P., and K.M.L. differentiated and characterized hPSCs. C.E.D., A.R., R.T.J., and R.S.K. performed genomics analyses. Y.S.K. performed electrophysiological studies, supervised by M.M. Y.M. and K.C.G. provided synthetic WNT agonists. H.A.U. and M.E.B. stained chicken embryos. H.G. and D.E.W. stained zebrafish embryos. B.D.M., J.M.A.L. and C.J.L. stained acorn worm embryos. M.E.B., D.E.W. and C.J.L. contributed to evolutionary discussions. C.E.D., R.T.J., and K.M.L. designed the study. K.M.L. supervised the study.

## Supplementary Figure Legends

**Supplementary Figure 1:**
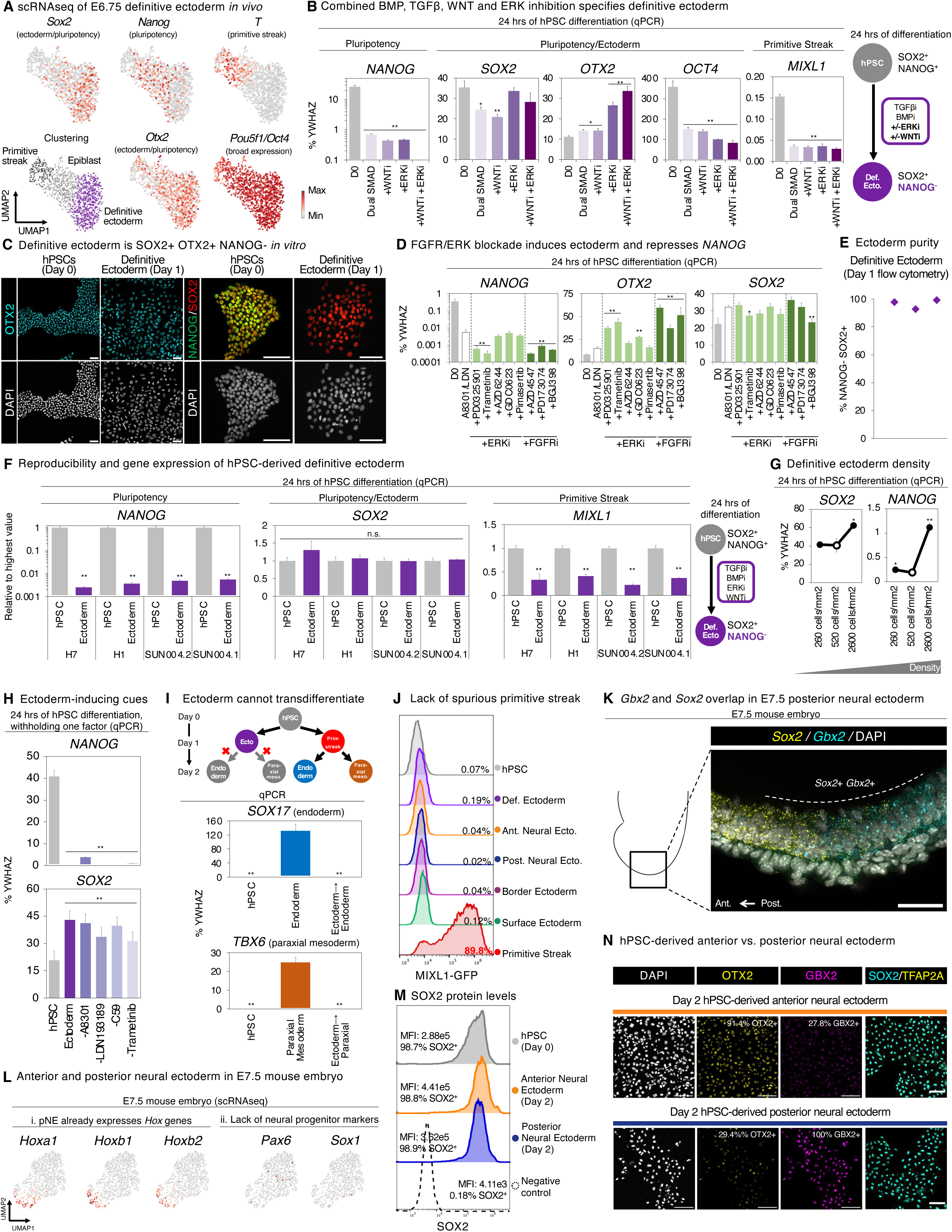
Differentiation of hPSCs into definitive ectoderm within 24 hours, and *in vivo* analysis of anterior and posterior neural ectoderm. A) Single-cell RNA-sequencing of E6.75 mouse embryos; analysis performed on a previously-published resource ^169^. Embryonic cells are shown, with extraembryonic cells excluded (*Elf5* and *Rhox5* log_2_ normalized expression = 0). *Otx2* is expressed in definitive ectoderm and pluripotent cells, consistent with previous studies ^16,79^. *Oct4* (*Pou5f1*) is broadly expressed across all cell-types, consistent with past studies ^70,71^. B) qPCR of H7 hPSCs differentiated for 24 hours with “dual SMAD inhibitors” (TGFβ inhibitor A-83-01 [1 μM] and BMP inhibitor LDN-193189 [250 nM]), in the presence or absence of WNT inhibitor (C59, 1 μM) and ERK inhibitor (Trametinib, 250 nM). Gene expression is shown relative to reference gene *YWHAZ*; 100% indicates that a gene is expressed at the same level as *YWHAZ*. “D0” = day-0 undifferentiated hPSCs. i: inhibitor. C) Immunostaining of hPSCs prior to, or after, differentiation into definitive ectoderm for 24 hours. NANOG and SOX2 immunostaining was performed with H7 hPSCs. OTX2 immunostaining was performed with H1 hPSCs. Scale bar = 100 μm. D) qPCR of H7 hPSCs differentiated for 24 hours with “dual SMAD inhibitors” (TGFβ inhibitor A-83-01 [1 μM] and BMP inhibitor LDN-193189 [250 nM]), in the presence or absence of individual ERK inhibitors (PD0325901, Trametinib, AZD6244, GDC0623 or Pimasertib) or FGFR inhibitors (AZD4547, PD173074 or BGJ398). Gene expression is shown relative to reference gene *YWHAZ*; 100% indicates that a gene is expressed at the same level as *YWHAZ*. D0: day-0 undifferentiated hPSCs. i: inhibitor. E) Flow cytometry of H1 hPSCs differentiated into definitive ectoderm for 24 hours. The percentage of SOX2+ NANOG-definitive ectoderm cells is shown. Individual datapoints depict the outcomes of independent experiments. F) qPCR of H1, H7, SUN004.1 and SUN004.2 hPSCs prior to, or after, differentiation into definitive ectoderm with BMP, ERK, TGFβ and WNT inhibitors (LDN-193189, Trametinib, A-83-01 and C59, respectively) for 24 hours. Gene expression is generally shown relative to the sample with the highest expression in each cell line. G) qPCR of H1 hPSCs seeded at different densities, and then differentiated into definitive ectoderm with BMP, ERK, TGFβ and WNT inhibitors (LDN-193189, Trametinib, A-83-01 and C59, respectively) for 24 hours. qPCR data normalized to undifferentiated hPSCs. Open circle indicates a seeding density of 520 cells/mm^2^, which was the baseline used throughout this study. Gene expression is shown relative to reference gene *YWHAZ*; 100% indicates that a gene is expressed at the same level as *YWHAZ*. H) qPCR of H1 hPSCs differentiated for 24 hours into definitive ectoderm with BMP, ERK, TGFβ and WNT inhibitors (LDN-193189, Trametinib, A-83-01 and C59, respectively), or with each ectoderm-inducing signal individually withheld. Gene expression is shown relative to reference gene *YWHAZ*; 100% indicates that a gene is expressed at the same level as *YWHAZ*. I) qPCR of H1 hPSCs that were either differentiated into anterior primitive streak ^78^ or definitive ectoderm for 24 hours, followed by treatment with either endoderm-inducing media ^77^ (Activin A [100 ng/mL] + LDN193189 [250 nM]) for 24 hours, or paraxial mesoderm-inducing media^78^ (A-83-01 [1 μM] + LDN193189 [250 nM] + CHIR99021 [3μM] + FGF2 [20 ng/mL]) for 24 hours. This revealed that hPSC-derived ectoderm largely failed to express either endoderm or paraxial mesoderm markers. By contrast, hPSC-derived primitive streak exposed to the same signals turned on either endoderm or paraxial mesoderm markers. Gene expression is shown relative to reference gene *YWHAZ*; 100% indicates that a gene is expressed at the same level as *YWHAZ*. J) Flow cytometry of HES3 *MIXL1-GFP* hPSCs differentiated into definitive ectoderm (day 1), anterior neural ectoderm (day 2), posterior neural ectoderm (day 2), border ectoderm (day 2), surface ectoderm (day 1), or mid primitive streak ^78^ (day 1, as a positive control to induce MIXL1 expression). Negligible numbers of MIXL1+ primitive streak cells emerged after ectoderm differentiation, reaffirming the suppression of spurious primitive streak differentiation. K) *In situ* staining of E7.5 mouse embryo, showing overlap of *Sox2* and *Gbx2* expression in pNE. Scale bar = 25 μm. L) Single-cell RNA-sequencing of E7.5 mouse embryos; analysis performed on a previously-published resource^169^. Neural ectoderm cells (identified by log_2_ normalized expression of *Sox2* > 0.5, and *Nanog, Tfap2a, Rhox5, Foxj1,* and *Elf5* = 0) are shown. M) Intracellular flow cytometry for SOX2 protein expression in H1 hPSCs, prior to or after differentiation into anterior neural ectoderm or posterior neural ectoderm for 2 days. The percentage of SOX2+ cells, and median fluorescence intensity (MFI) of SOX2 expression, are both shown. N) Immunostaining of H1 hPSCs differentiated into definitive ectoderm for 24 hours, followed by differentiation into aNE or pNE for 24 hours. Scale bar = 100 μm. qPCR data depicts the mean of two biological replicates, with s.e.m. shown. For flow cytometry and *in vitro* staining, a representative image from two biological replicates is shown.

**Supplementary Figure 2:**
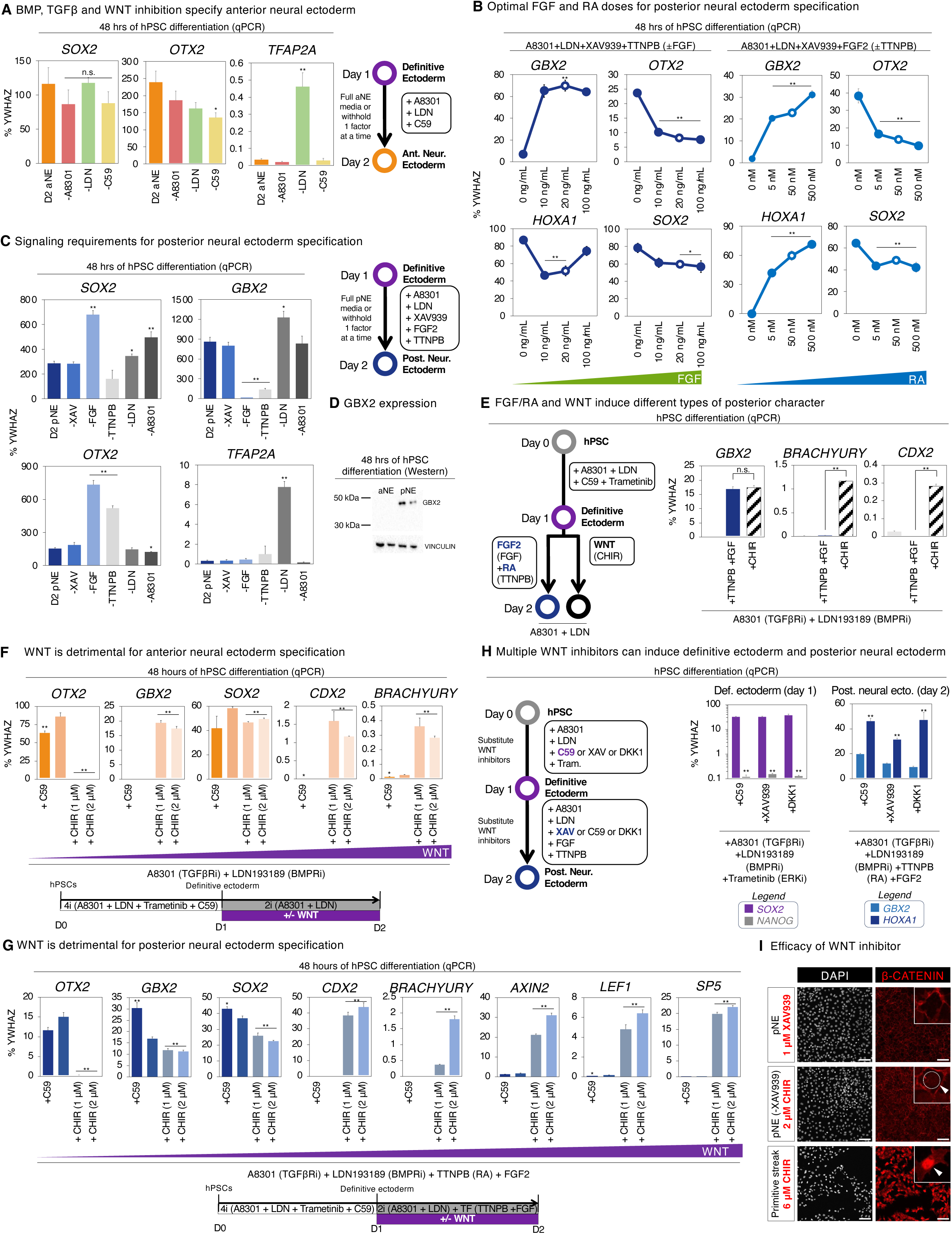
Differentiation of hPSCs into anterior or posterior neural ectoderm within 48 hours, and *in vivo* analysis of signaling fields. A) qPCR of H1 hPSCs differentiated into definitive ectoderm for 24 hours, followed by BMP, TGFβ and WNT inhibitors (LDN-193189, A-83-01 and C59, respectively) for 24 hours to generate anterior neural ectoderm. To assess the importance of each anterior neural ectoderm-inducing signal, each factor was individually withheld. Gene expression is shown relative to reference gene *YWHAZ*; 100% indicates that a gene is expressed at the same level as *YWHAZ*. D2 aNE: day-2 anterior neural ectoderm. B) qPCR of H1 hPSCs were differentiated into definitive ectoderm for 24 hours, followed by differentiation into posterior neural ectoderm for 24 hours. During posterior neural ectoderm induction, different doses of FGF agonist (FGF2) or RA agonist (TTNPB) were added to assess dose-dependent effects. Open circles indicate the optimal doses of FGF2 (20 ng/mL) and TTNPB (50 nM) for posterior neural ectoderm induction used as a baseline throughout this study. Gene expression is shown relative to reference gene *YWHAZ*; 100% indicates that a gene is expressed at the same level as *YWHAZ*. C) qPCR of H1 hPSCs differentiated into definitive ectoderm for 24 hours, followed by a combination of FGF and RA agonists (FGF2 and TTNPB, respectively) together with BMP, TGFβ and WNT inhibitors (LDN-193189, A-83-01 and C59, respectively) for 24 hours to generate posterior neural ectoderm. To assess the importance of each posterior neural ectoderm-inducing signal, each factor was individually withheld. Gene expression is shown relative to reference gene *YWHAZ*; 100% indicates that a gene is expressed at the same level as *YWHAZ*. “D2 pNE”: day-2 posterior neural ectoderm. D) Western Blot to detect the expression of GBX2 protein in H1 hPSC-derived day-2 aNE or day-2 pNE. VINCULIN protein serves as a loading control. E) qPCR of H1 hPSCs differentiated into definitive ectoderm for 24 hours, followed by exposure to (1) neural-inducing signals (A-83-01 + LDN-193189), (2) neural-inducing signals (A-83-01 + LDN-193189) + FGF2 (20 ng/mL) + TTNPB (50 nM) or (3) neural-inducing signals (A-83-01 + LDN-193189) + CHIR99021 (2 μM) for 24 hours. Gene expression is shown relative to reference gene *YWHAZ*; 100% indicates that a gene is expressed at the same level as *YWHAZ*. F) qPCR of H1 hPSCs were differentiated into definitive ectoderm for 24 hours, followed by differentiation into anterior neural ectoderm for 24 hours (with A-83-01, LDN-193189, and C59). Alternatively, to test the effect of WNT signaling on anterior neural ectoderm induction, WNT inhibitor (C59) was withheld, and WNT agonist (CHIR99021) was added instead. Gene expression is shown relative to reference gene *YWHAZ*; 100% indicates that a gene is expressed at the same level as *YWHAZ*. G) qPCR of H1 hPSCs were differentiated into definitive ectoderm for 24 hours, followed by differentiation into posterior neural ectoderm for 24 hours (with A-83-01, LDN-193189, C59, FGF2 and TTNPB). Alternatively, to test the effect of WNT signaling on posterior neural ectoderm induction, WNT inhibitor (C59) was withheld, and WNT agonist (CHIR99021) was added instead. Gene expression is shown relative to reference gene *YWHAZ*; 100% indicates that a gene is expressed at the same level as *YWHAZ*. H) First, qPCR was performed of H1 hPSCs that were differentiated into definitive ectoderm for 24 hours with A8301, LDN193189, Trametinib, and WNT inhibitor (either C59 [1 μM], XAV939 [1 μM], or DKK1 [300 ng/mL]), revealing that all three WNT inhibitors comparably induced definitive ectoderm (*left*). Second, qPCR was performed of H1 hPSCs that were differentiated into definitive ectoderm for 24 hours with A8301, LDN193189, Trametinib, and XAV939, followed by differentiation into posterior neural ectoderm for 24 hours with A8301, LDN193189, TTNPB, FGF2, and WNT inhibitor (either C59 [1 μM], XAV939 [1 μM], or DKK1 [300 ng/mL]), revealing that all three WNT inhibitors comparably induced posterior neural ectoderm (*right*). Given these comparable results, throughout the rest of this study we applied C59 for definitive ectoderm induction and XAV939 for posterior neural ectoderm induction, as indicated in bold in the accompanying cartoon. Gene expression is shown relative to reference gene *YWHAZ*; 100% indicates that a gene is expressed at the same level as *YWHAZ*. I) β-CATENIN immunostaining of H1 hPSCs that were differentiated into various cell-types. First, hPSCs were differentiated into definitive ectoderm for 24 hours, and then treated with posterior neural ectoderm-inducing media for 24 hours (A8301 [1 μM] + LDN193189 [100 nM] + TTNPB [50 nM] + FGF2 [20 ng/mL] + XAV939 [1 μM] + Thiazovivin [1 μM]), or as a positive control to induce nuclear β-CATENIN, modified posterior neural ectoderm media lacking WNT inhibitor and including a WNT agonist for 24 hours (A8301 [1 μM] + LDN193189 [100 nM] + TTNPB [50 nM] + FGF2 [20 ng/mL] + CHIR99021 [2 μM] + Thiazovivin [1 μM]). Alternatively, as a positive control to induce nuclear β-CATENIN, hPSCs were differentiated into mid primitive streak, with WNT agonist-containing media (Activin A [30 ng/mL] + BMP4 [40 ng/mL] + CHIR99021 [6 μM] + FGF2 [20 ng/mL]) for 24 hours. Nuclear β-CATENIN was absent from hPSC-derived posterior neural ectoderm, and either weakly or strongly positive in the two WNT agonist-treated cell populations. Scale bar = 100 μm. qPCR data depicts the mean of two biological replicates, with s.e.m. shown. For *in vitro* staining, a representative image from two biological replicates is shown.

**Supplementary Figure 3:**
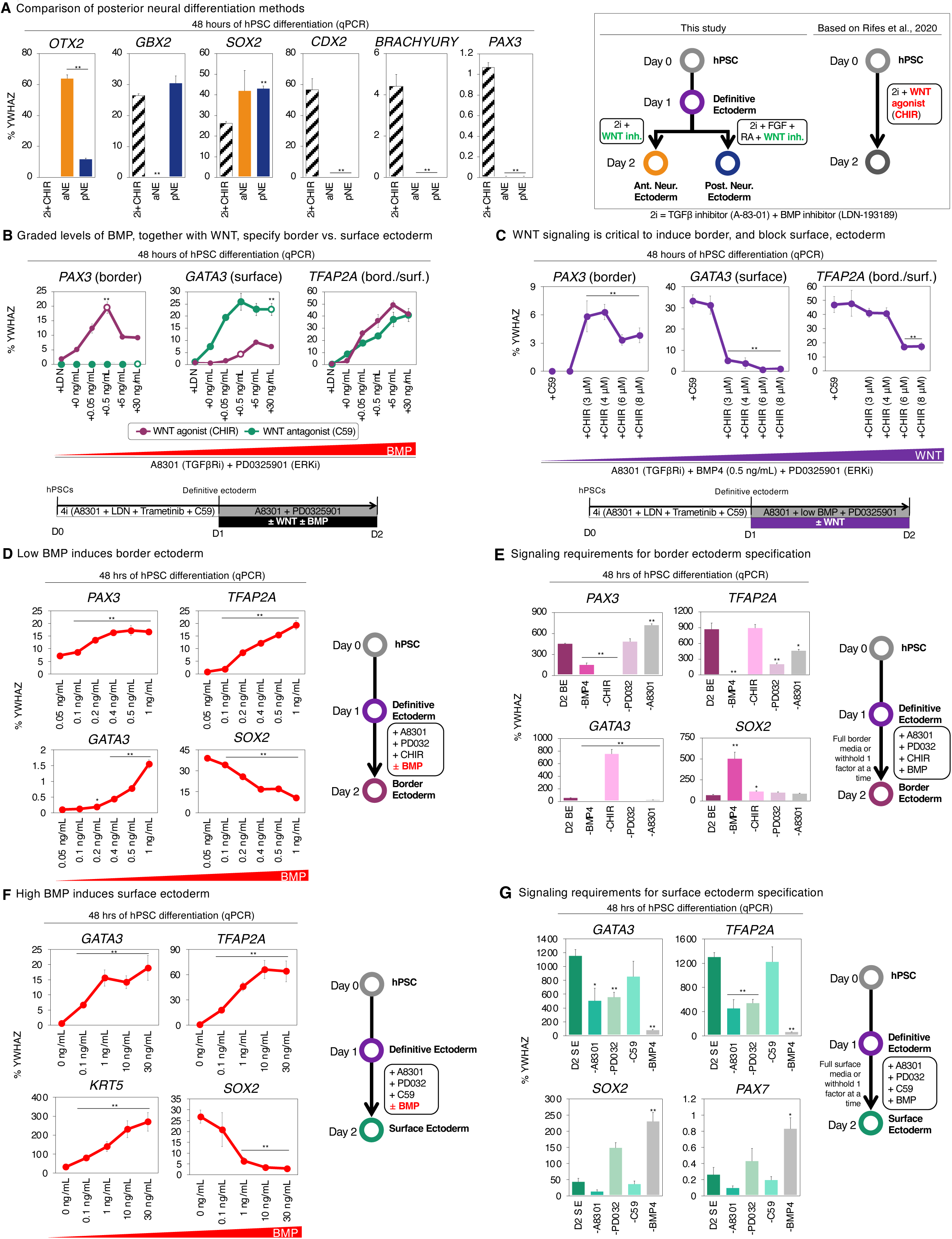
Differentiation of hPSCs into border or surface ectoderm within 48 hours, and comparison of posterior neural differentiation methods. A) qPCR of H1 hPSCs differentiated for 48 hours, using two separate strategies. First, hPSCs were differentiated into definitive ectoderm for 24 hours, followed by either anterior or posterior neural ectoderm induction for 24 hours (as described in this study). Alternatively, hPSCs were differentiated with BMP and TGFβ inhibitors (LDN-193189 and A-83-01, respectively; “2i”) together with WNT agonist (CHIR99021, 2 μM) for 48 hours, similarly to a previous report ^43^. Gene expression is shown relative to reference gene *YWHAZ*; 100% indicates that a gene is expressed at the same level as *YWHAZ*. B) qPCR of H1 hPSCs differentiated into definitive ectoderm for 24 hours, followed by treatment with non-neural signals (A-83-01 and PD0325901), in combination with different levels of BMP signaling (BMP agonist BMP4 [0.05-30 ng/mL] or alternatively, BMP inhibitor LDN-193189), in the presence or absence of either WNT agonist (CHIR99021, 3 μM) or WNT inhibitor (C59, 1 μM), for 24 hours. White circles indicate optimal BMP doses for border ectoderm induction (0.5 ng/mL BMP4) or surface ectoderm induction (30 ng/mL BMP4). Gene expression is shown relative to reference gene *YWHAZ*; 100% indicates that a gene is expressed at the same level as *YWHAZ*. C) qPCR of H1 hPSCs differentiated into definitive ectoderm for 24 hours, followed by border ectoderm-inducing signals (A-83-01, BMP4 [0.5 ng/mL], and PD0325901), in combination with different levels of WNT signaling (WNT agonist CHIR99021 [3-8 μM] or WNT inhibitor C59), for 24 hours. D) qPCR of H1 hPSCs differentiated into definitive ectoderm for 24 hours, followed by border ectoderm-inducing signals (A-83-01, CHIR99021 [3 μM], and PD0325901), in combination with different levels of BMP signaling (BMP agonist BMP4 [0.05-30 ng/mL] or alternatively, BMP inhibitor LDN-193189), for 24 hours. Gene expression is shown relative to reference gene *YWHAZ*; 100% indicates that a gene is expressed at the same level as *YWHAZ*. E) qPCR of H1 hPSCs differentiated into definitive ectoderm for 24 hours, followed by border ectoderm-inducing signals (A-83-01, BMP4, CHIR99021, and PD0325901) for 24 hours to generate border ectoderm. To assess the importance of each border ectoderm-inducing signal, each factor was individually withheld. Gene expression is shown relative to reference gene *YWHAZ*; 100% indicates that a gene is expressed at the same level as *YWHAZ*. D2 BE: day-2 border ectoderm. F) qPCR of H1 hPSCs differentiated into definitive ectoderm for 24 hours, followed by surface ectoderm-inducing signals (A-83-01, C59, and PD0325901), in the presence or absence of BMP4 (0-30 ng/mL), for 24 hours. Gene expression is shown relative to reference gene *YWHAZ*; 100% indicates that a gene is expressed at the same level as *YWHAZ*. G) qPCR of H1 hPSCs differentiated into definitive ectoderm for 24 hours, followed by surface ectoderm-inducing signals (A-83-01, BMP4, C59, and PD0325901) for 24 hours. To assess the importance of each border ectoderm-inducing signal, each factor was individually withheld. Gene expression is shown relative to reference gene *YWHAZ*; 100% indicates that a gene is expressed at the same level as *YWHAZ*. D2 SE: day-2 surface ectoderm. qPCR data depicts the mean of two biological replicates, with s.e.m. shown.

**Supplementary Figure 4:**
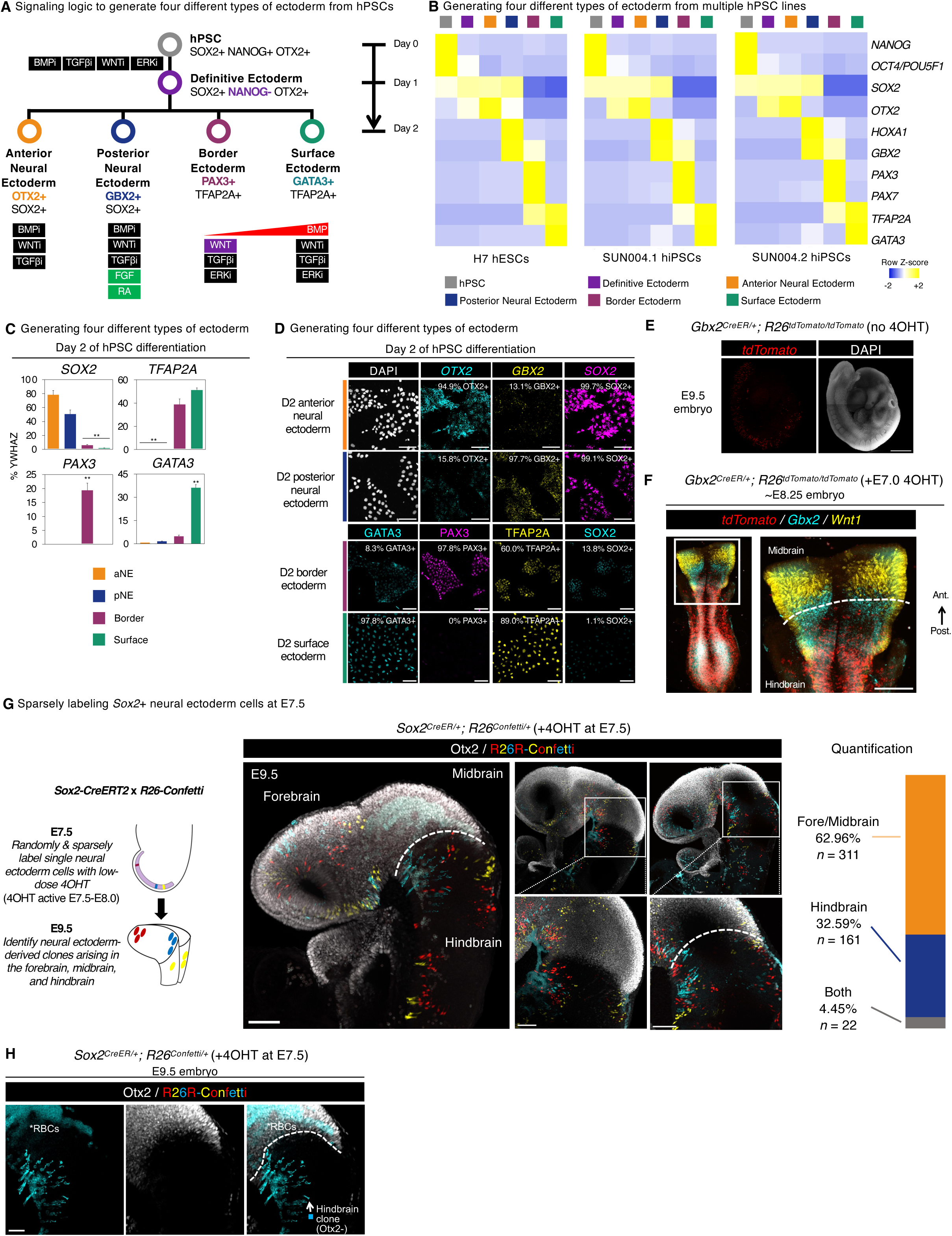
Differentiation of hPSCs into anterior neural, posterior neural, border or surface ectoderm within 48 hours, and lineage tracing of neural ectoderm cells within mouse embryos. A) Summary of the hPSC differentiation strategy described in this study. B) qPCR of H7 human embryonic stem cells (hESCs), SUN004.1 human induced pluripotent stem cells (hiPSCs), and SUN004.2 hiPSCs differentiated into definitive ectoderm for 24 hours, followed by either anterior neural ectoderm, posterior neural ectoderm, border ectoderm or surface ectoderm, for 24 hours. The heatmap depicts the Z-score, indicating the variation of gene expression across cell-types for a given gene; two biological replicates were used per cell-type. C) qPCR of H1 hPSCs differentiated into definitive ectoderm for 24 hours, followed by differentiation into either aNE, pNE, border ectoderm or surface ectoderm for 24 hours. Gene expression is shown relative to reference gene *YWHAZ*. D) Top: *In situ* hybridization of H1 hPSCs differentiated into definitive ectoderm for 24 hours, followed by differentiation into aNE or pNE for 24 hours. Scale: 100 μm. Bottom: Immunostaining of H1 hPSCs differentiated into definitive ectoderm for 24 hours, followed by differentiation into border or surface ectoderm for 24 hours. Scale: 100 μm. E) *In situ* staining of E9.5 *Gbx2-CreER*; *ROSA26-tdTomato* mouse embryo that was not exposed to 4OHT. This negative control revealed minimal *Gbx2-CreER*-driven recombination in the absence of 4OHT. Scale bar = 500 μm. F) E7.0 *Gbx2-CreER*; *ROSA26-LoxP-STOP-LoxP-tdTomato* mouse embryos were exposed to 1 mg 4OHT (intraperitoneally delivered into pregnant females) to label *Gbx2*^+^ pNE with tdTomato. Subsequently, *in situ* staining was performed on E8.25 embryos to visualize pNE-derived progeny in relation to the expression of *Wnt1* (which marks the midbrain/hindbrain boundary) and *Gbx2* (which marks the hindbrain). Scale bar = 100 μm. G) E7.5 *Sox2-CreER*; *Confetti* mouse embryos were exposed to 0.5 mg 4OHT (intraperitoneally delivered into pregnant females) to label single E7.5 *Sox2*^+^ neural ectoderm progenitors with one of three detectable fluorescent colors (red, cyan, or yellow). Subsequently, E9.5 embryos were immunostained to visualize neural ectoderm-derived cell clusters and where they resided in relation to the forebrain, midbrain, and hindbrain; photos of three independent embryos are shown (*left*). The forebrain/midbrain domain was molecularly determined by Otx2 immunostaining (*left*). Quantification of the locations of all neural ectoderm-derived cell clusters analyzed in this study, totaling 494 cell clusters from 16 embryos, obtained from 7 litters (*right*). Scale bar = 100 μm. H) Detailed analysis of the spatial position of a E7.5 *Sox2*^+^ neural ectoderm-derived cell cluster relative to the midbrain/hindbrain boundary. E7.5 *Sox2-CreER*; *Confetti* mouse embryos were exposed to 0.5 mg 4OHT (intraperitoneally delivered into pregnant females) to label single E7.5 *Sox2*^+^ neural ectoderm progenitors with one of three detectable fluorescent colors (red, cyan, or yellow). Subsequently, E9.5 embryos were immunostained to visualize neural ectoderm-derived cell clusters and where they resided in relation to the forebrain, midbrain, and hindbrain. The forebrain/midbrain domain was molecularly determined by Otx2 immunostaining. This is a higher magnification image of an embryo shown in **Fig. S4g**. Asterisks indicate autofluorescence from red blood cells (RBCs). Scale bar = 100 μm.

**Supplementary Figure 5:**
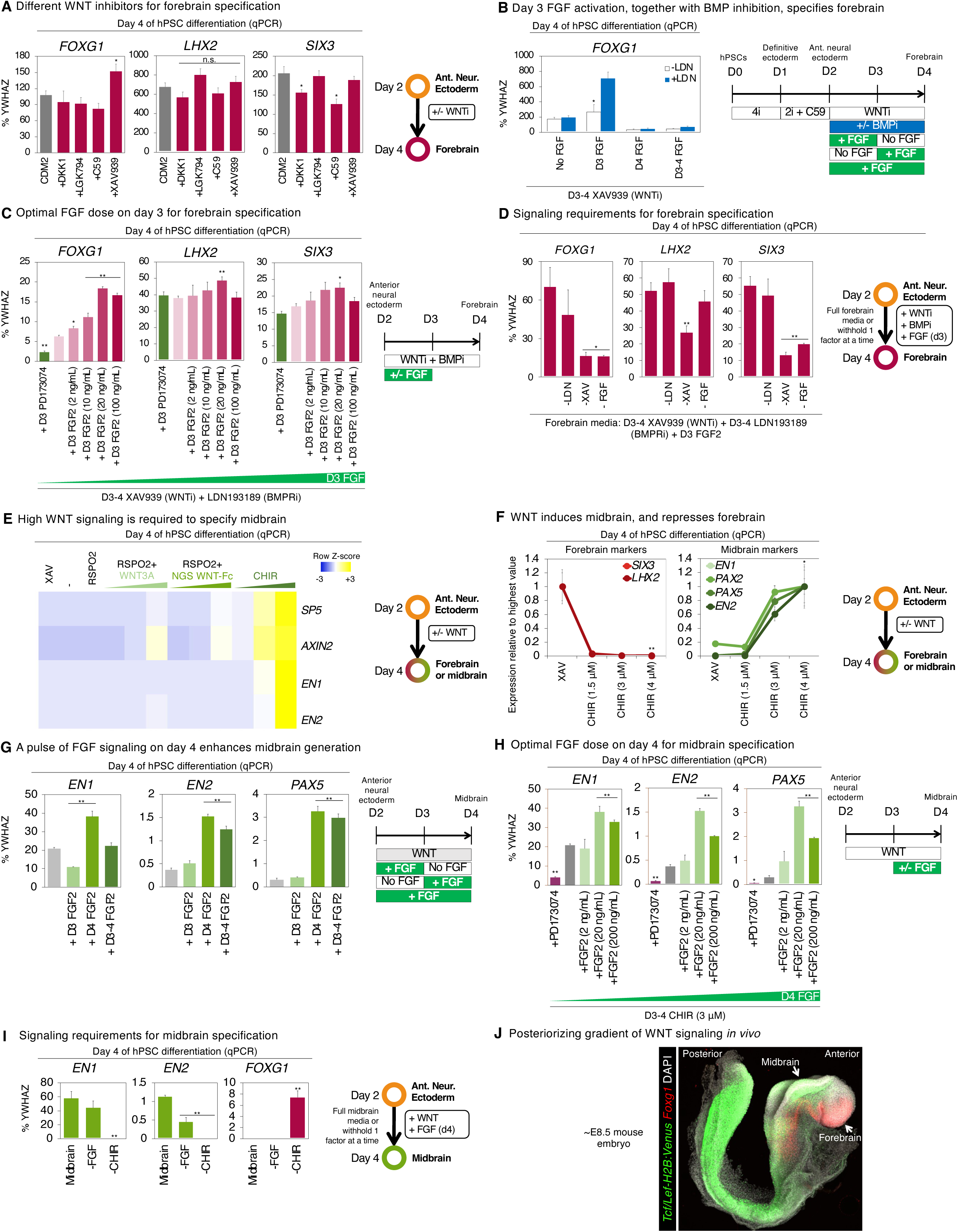
Differentiation of hPSCs into forebrain or midbrain progenitors within 4 days. A) qPCR of H1 hPSCs differentiated into definitive ectoderm for 24 hours, followed by anterior neural ectoderm for 24 hours, and then subsequently treated with either basal medium (CDM2) for 48 hours, or supplemented with WNT inhibitor DKK1 (LRP5/6 inhibitor, 300 ng/mL), LGK974 (Porcupine inhibitor, 100 nM), C59 (Porcupine inhibitor, 1 μM), or XAV939 (Tankyrase inhibitor, 1 μM). This revealed that XAV939 was most effective at upregulating forebrain marker *FOXG1*. Gene expression is shown relative to reference gene *YWHAZ*; 100% indicates that a gene is expressed at the same level as *YWHAZ*. B) qPCR of H1 hPSCs differentiated into definitive ectoderm for 24 hours, followed by anterior neural ectoderm for 24 hours, and then subsequently treated with XAV939 for 48 hours to generate forebrain progenitors. During the 48 hour period of forebrain induction, FGF2 (20 ng/mL) was added for the first 24 hour interval (“day 3”), the second 24 hour interval (“day 4”), or the entirety of the 48 hour interval (“days 3-4”), in the presence or absence of BMP receptor inhibitor LDN193189 (100 nM). This revealed that day 3 FGF2 treatment, together with BMP inhibition on days 3-4, was most effective at upregulating forebrain markers *FOXG1* and *SIX3*. Gene expression is shown relative to reference gene *YWHAZ*; 100% indicates that a gene is expressed at the same level as *YWHAZ*. C) qPCR of H7 hPSCs differentiated into definitive ectoderm for 24 hours, followed by anterior neural ectoderm for 24 hours, and then subsequently treated with XAV939 + LDN193189 + FGF2 (2-100 ng/mL) for 24 hours, and finally XAV939 + LDN193189 for 24 hours to generate forebrain progenitors. On the third day of differentiation (“D3”), different doses of FGF2 (2-100 ng/mL) were tested, or alternatively, exogenous FGF2 was withheld (no notation), in the presence or absence of FGFR inhibitor PD173074 (100 ng/mL). This revealed that 20 ng/mL of FGF2 on day 3 of differentiation was most effective at upregulating forebrain markers *SIX3* and *FOXG1*, consistent with how FGF induces *Foxg1* in the mouse embryonic forebrain^203^. Gene expression is shown relative to reference gene *YWHAZ*; 100% indicates that a gene is expressed at the same level as *YWHAZ*. D) qPCR of H1 hPSCs differentiated into definitive ectoderm for 24 hours, followed by anterior neural ectoderm for 24 hours, and then subsequently treated with XAV939 + LDN193189 + FGF2 for 24 hours, and finally XAV939 + LDN193189 for 24 hours to generate forebrain progenitors. To assess the importance of each forebrain-inducing signal (XAV939, LDN193189 or FGF2), each individual signal was respectively withheld during the 48 hour interval of forebrain induction. Gene expression is shown relative to reference gene *YWHAZ*; 100% indicates that a gene is expressed at the same level as *YWHAZ*. E) qPCR of H1 hPSCs differentiated into definitive ectoderm for 24 hours, followed by anterior neural ectoderm for 24 hours, and then subsequently treated with the following WNT pathway modulators for 48 hours: WNT inhibitor (XAV939, 1 μM), RSPO2 (25 nM), RSPO2 + WNT3A (increasing concentrations of 10, 100 or 1000 ng/mL), RSPO2 + NGS WNT-Fc^201^ (increasing concentrations of 0.03, 0.3 or 3 nM), or CHIR99021 (increasing concentrations of 0.75, 1.5 or 3 μM). For WNT3A or NGS WNT-Fc treatment conditions, RSPO2 was added to increase cell-surface levels of FRIZZLED receptors, thereby enhancing cellular responsiveness to these WNT pathway agonists^201^. NGS-WNT-Fc refers to an Fc-tagged synthetic WNT pathway agonist that acts by heterodimerizing FZD and LRP6 receptors^201^. F) qPCR of H1 hPSCs differentiated into definitive ectoderm for 24 hours, followed by anterior neural ectoderm for an additional 24 hours, and then subsequently treated with either WNT inhibitor (XAV939, 1 μM) or WNT agonist (CHIR99021, 1.5-4 μM) for an additional 48 hours. This showed that WNT induced midbrain progenitors, whereas WNT inhibitor specified forebrain progenitors. Gene expression is shown relative to the sample with the highest expression in this experiment. G) qPCR of H1 hPSCs differentiated into definitive ectoderm for 24 hours, followed by anterior neural ectoderm for 24 hours, and then subsequently differentiated towards midbrain with WNT agonist (CHIR99021, 3 μM) for 48 hours, in the presence or absence of FGF2 (20 ng/mL) on the first 24 hour interval, the second 24 hour interval, or the entire 48 interval of midbrain induction. Gene expression is shown relative to reference gene *YWHAZ*; 100% indicates that a gene is expressed at the same level as *YWHAZ*. H) qPCR of H1 hPSCs differentiated into definitive ectoderm for 24 hours, followed by anterior neural ectoderm for 24 hours, and then subsequently differentiated towards midbrain with WNT agonist (CHIR99021, 3 μM) for 48 hours, in the presence or absence of FGF2 (2-200 ng/mL) on last 24 hours of midbrain induction. Gene expression is shown relative to reference gene *YWHAZ*; 100% indicates that a gene is expressed at the same level as *YWHAZ*. I) qPCR of H1 hPSCs differentiated into definitive ectoderm for 24 hours, followed by anterior neural ectoderm for 24 hours, and then subsequently differentiated towards midbrain for 48 hours (CHIR99021 on days 3-4, and FGF2 on day 4). To assess the importance of each midbrain-inducing signal (CHIR99021 or FGF2), each individual signal was respectively withheld during the 48 hour interval of midbrain induction. J) *In situ* staining of an E8.5 *Tcf/Lef:H2B:Venus* reporter mouse embryo^173^ for *Foxg1* (forebrain marker) and *Venus* (indicating WNT transcriptional response). This disclosed a posteriorizing gradient of WNT signaling, with minimal WNT signaling in *Foxg1*^+^ forebrain, but high WNT signaling in the midbrain. qPCR data depicts the mean of two biological replicates, with s.e.m. shown.

**Supplementary Figure 6:**
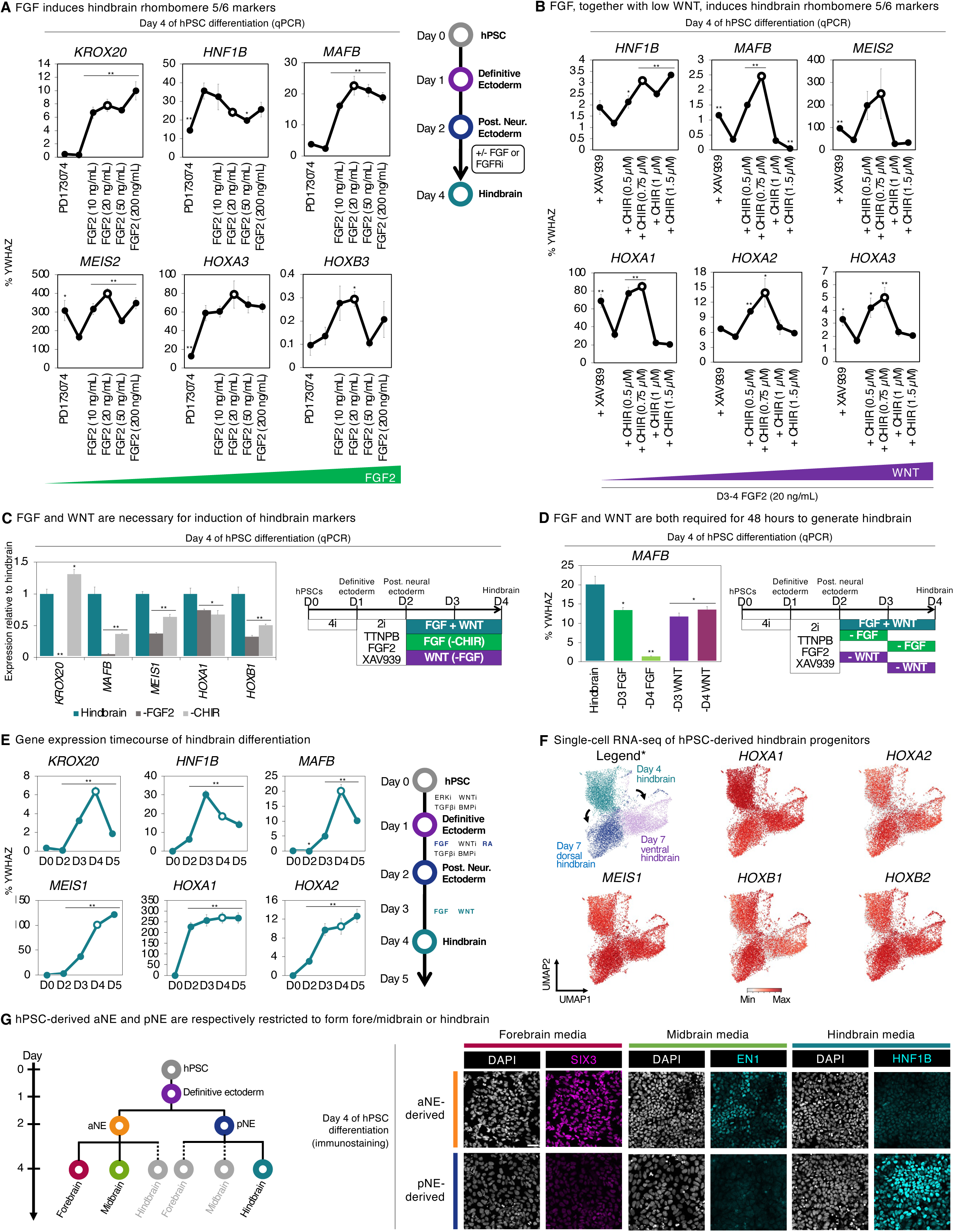
Differentiation of hPSCs into hindbrain progenitors within 4 days. A) qPCR of H1 hPSCs differentiated into definitive ectoderm for 24 hours, followed by posterior neural ectoderm for 24 hours, and then further differentiated for 48 hours in the presence or absence of FGF2 (10-200 ng/mL) or FGFR inhibitor (PD173074, 100 nM). qPCR data normalized to undifferentiated hPSCs. Gene expression is shown relative to reference gene *YWHAZ*; 100% indicates that a gene is expressed at the same level as *YWHAZ*. FGFR: FGF receptor. B) qPCR of H1 hPSCs differentiated into definitive ectoderm for 24 hours, followed by posterior neural ectoderm for 24 hours, and then further differentiated for 48 hours in FGF2 (20 ng/mL), in the presence or absence of WNT agonist (CHIR99021, 0.5-1.5 μM) or WNT inhibitor (XAV939, 1 μM). Gene expression is shown relative to reference gene *YWHAZ*; 100% indicates that a gene is expressed at the same level as *YWHAZ*. C) qPCR of H1 hPSCs differentiated into definitive ectoderm for 24 hours, followed by posterior neural ectoderm for 24 hours, and then further differentiated into hindbrain for 48 hours by FGF2 + CHIR99021. Alternatively, during the 48 hour interval of hindbrain induction, either FGF2 or CHIR99021 were individually withheld. This revealed that both FGF and WNT signaling are required for maximal *MAFB* expression. qPCR data normalized to levels found in hindbrain progenitors (e.g., cells treated with FGF2 + CHIR99021). D) qPCR of H1 hPSCs differentiated into definitive ectoderm for 24 hours, followed by posterior neural ectoderm for 24 hours, and then further differentiated into hindbrain for 48 hours by FGF2 + CHIR99021. Alternatively, during the 48 hour interval of hindbrain induction, either FGF2 or CHIR99021 were individually withheld for 24 intervals (“day 3” = the first 24 hour interval, whereas “day 4” = the second 24 hour interval). This revealed that both FGF and WNT signaling are continuously required for 48 hours to achieve maximal *MAFB* expression. Gene expression is shown relative to reference gene *YWHAZ*; 100% indicates that a gene is expressed at the same level as *YWHAZ*. E) qPCR of H1 hPSCs differentiated into definitive ectoderm for 24 hours, followed by posterior neural ectoderm for 24 hours, and then further differentiated into hindbrain by FGF2 + CHIR99021 for 24, 48 or 72 hours (*left*). This revealed that FGF and WNT treatment for 48 hours was optimal for hindbrain specification (as denoted by a white circle), which was used as a baseline for the remainder of this study. qPCR data normalized to undifferentiated hPSCs. Cartoon of hPSC differentiation into hindbrain progenitors, with various signaling pathways activated or inhibited in a temporally dynamic way at each stage (*right*). F) scRNAseq of H7 hPSC-derived day-4 hindbrain progenitors, and day-7 dorsal hindbrain or ventral hindbrain progenitors. Colors denote differentiation conditions. An asterisk (*) indicates that the legend is identical to that shown in Fig. 4c, and is reproduced here to aid the interpretation of other images shown in this subpanel. G) H1 hPSCs were differentiated into definitive ectoderm for 24 hours, and then subsequently differentiated into either anterior neural ectoderm (aNE) or posterior neural ectoderm (pNE) within an additional 24 hours. hPSC-derived aNE and pNE cells were then dissociated and then replated in either forebrain-, midbrain-, or hindbrain-inducing signals for 48 hours, and then immunostaining was performed. Scale bar = 50 μm qPCR data depicts the mean of two biological replicates, with s.e.m. shown. For *in vitro* staining, a representative image from two biological replicates is shown.

**Supplementary Figure 7:**
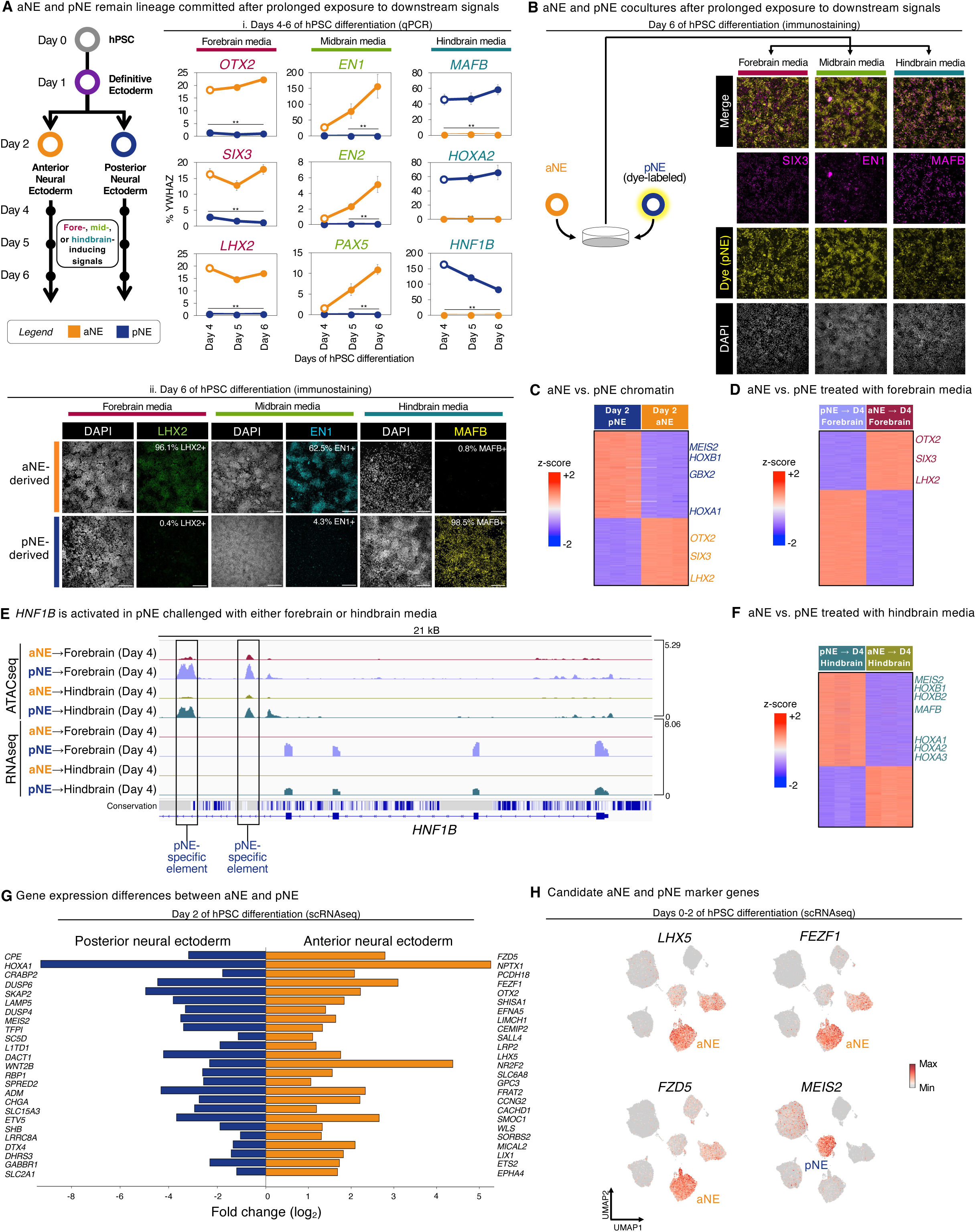
Accessible chromatin landscapes and lineage commitment of hPSC-derived anterior vs. posterior neural ectoderm. A) H1 hPSCs were differentiated into either anterior or posterior neural ectoderm for 2 days. They were then respectively challenged with either forebrain-, midbrain-, or hindbrain-inducing signals for 2, 3 or 4 days. i) qPCR was performed on days 4, 5 and 6 of hPSC differentiation. Gene expression is shown relative to reference gene *YWHAZ*; 100% indicates that a gene is expressed at the same level as *YWHAZ*. ii) Immunostaining was performed on day 6 of differentiation. Scale = 100 μm. B) H1 hPSCs were differentiated into either aNE or pNE within 2 days; pNE was fluorescently labeled, and then aNE (uncolored) and pNE (dye-labeled) were mixed. Cocultures were treated with either forebrain-, midbrain-, or hindbrain-inducing signals for 4 days, prior to immunostaining (i.e., immunostaining was performed on day 6 of hPSC differentiation). aNE: day-2 anterior neural ectoderm. pNE: day-2 posterior neural ectoderm. C) OmniATACseq was performed on day 2 H7 hPSC-derived aNE vs. pNE. A Z-score heatmap is shown, depicting all differentially accessible regions between these two cell-types (false discovery rate [FDR]<0.05). Each row indicates an individual genomic element. D) OmniATACseq was performed on day 2 H7 hPSC-derived aNE vs. pNE that were challenged with forebrain-inducing signals for 2 additional days (1 day of XAV939 [1 μM] + LDN193189 [100 nM] + FGF2 [20 ng/mL], followed by 1 day of XAV939 (1 μM) and LDN193189 [100 nM]). A Z-score heatmap is shown, depicting all differentially accessible regions between these two cell-types (FDR<0.05). Each row indicates an individual genomic element. E) OmniATACseq and RNAseq of hPSC-derived aNE and pNE that were treated with either forebrain- or hindbrain-inducing signals for 48 hours. F) OmniATACseq was performed on day 2 H7 hPSC-derived aNE vs. pNE that were challenged with hindbrain-inducing signals (CHIR99021 [750 nM] + FGF2 [20 ng/mL]) for 2 additional days. A Z-score heatmap is shown, depicting all differentially accessible regions between these two cell-types (FDR<0.05). Each row indicates an individual genomic element. G) Top 25 differentially expressed genes between H7 hPSC-derived day-2 anterior neural ectoderm vs. posterior neural ectoderm, as determined by scRNAseq. The top 25 differentially expressed genes were determined by P value, and all these genes showed statistically significant expression differences (P<0.05). H) Single-cell RNA-sequencing of H7 hPSCs differentiated into day-1 definitive ectoderm, day-2 anterior neural ectoderm, day-2 posterior neural ectoderm, day-2 border ectoderm, or day-2 surface ectoderm. Colors denote differentiation conditions. *Fzd5*, *Lhx5*, and *Fezf1* have previously been associated with anterior neural identity *in vivo* at later developmental stages^204-206^, whereas *Meis2* has been previously associated with posterior neural identity *in vivo*^207^. qPCR data depicts the mean of two biological replicates, with s.e.m. shown. For *in vitro* staining, a representative image from two biological replicates is shown. OmniATACseq and bulk-population RNAseq were each performed on two biological replicates. *In vitro* scRNAseq data is from a single experiment.

**Supplementary Figure 8:**
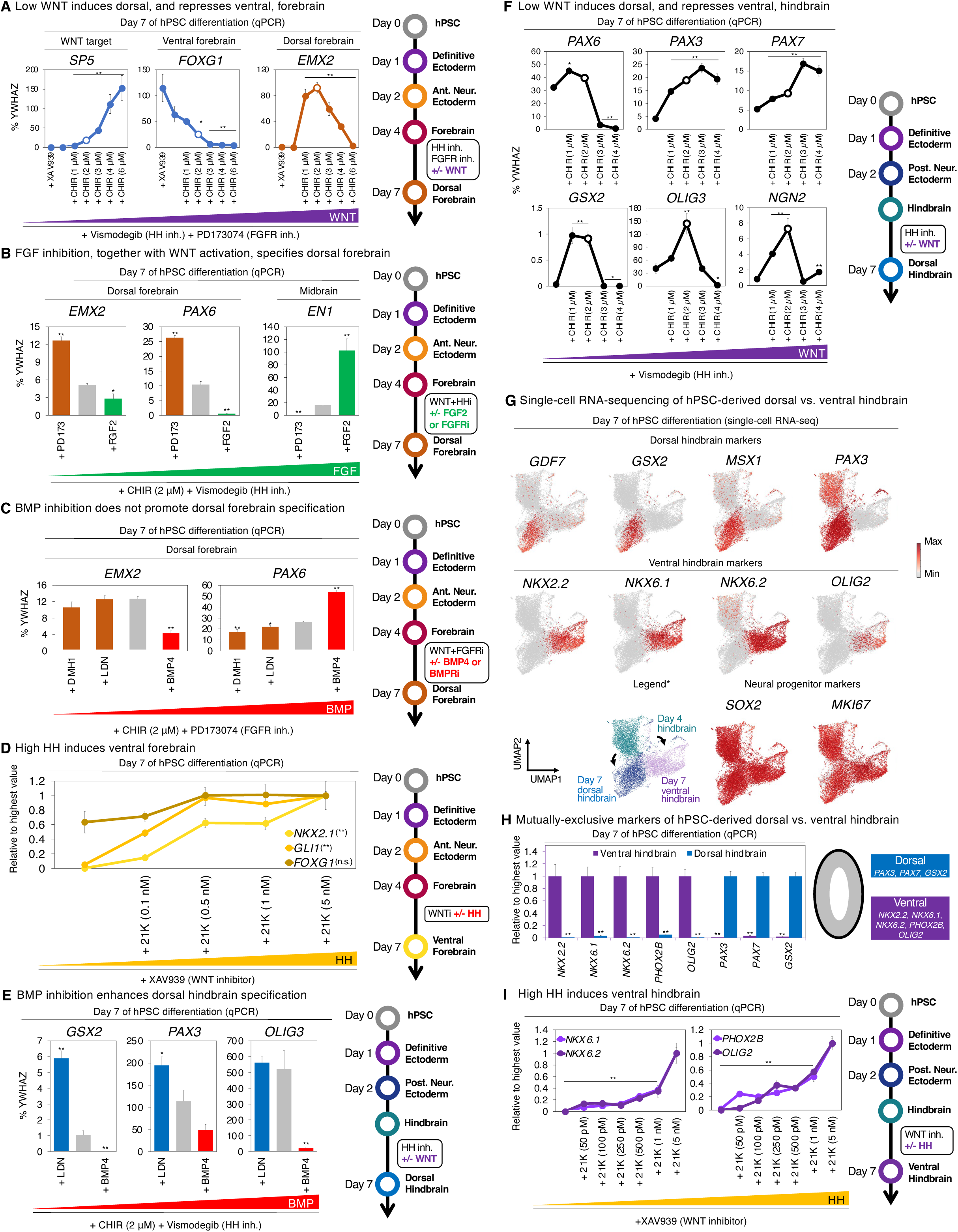
Differentiation of hPSCs into dorsal forebrain and ventral forebrain progenitors within 7 days. A) H1 hPSCs were differentiated into definitive ectoderm within 24 hours, and then anterior neural ectoderm within 24 hours, followed by forebrain progenitors within 48 hours. Forebrain progenitors were then treated with Vismodegib + PD173074 for 72 hours, in the presence or absence of WNT agonist (CHIR99021, 1-6 μM) or WNT inhibitor (XAV939, 1 μM). This revealed that 2 μM of CHIR99021 (indicated by white circle) was optimal for inducing dorsal forebrain marker *EMX2*. Gene expression is shown relative to reference gene *YWHAZ*; 100% indicates that a gene is expressed at the same level as *YWHAZ*. HH inh.: HEDGEHOG inhibitor. FGFR inh.: FGF receptor inhibitor. B) H1 hPSCs were differentiated into definitive ectoderm within 24 hours, and then anterior neural ectoderm within 24 hours, followed by forebrain progenitors within 48 hours. Forebrain progenitors were then treated with CHIR99021 (2 μM) + PD173074 (100 nM) for 72 hours, in the presence or absence of FGF2 (20 ng/mL) or FGFR inhibitor (PD173074, 100 nM). This revealed that FGF inhibition enhanced dorsal forebrain markers *EMX2* and *PAX6*, while preventing erroneous WNT-induced expression of midbrain marker *EN1*. Gene expression is shown relative to reference gene *YWHAZ*; 100% indicates that a gene is expressed at the same level as *YWHAZ*. FGFRi: FGF receptor inhibitor. C) H1 hPSCs were differentiated into definitive ectoderm within 24 hours, and then anterior neural ectoderm within 24 hours, followed by forebrain progenitors within 48 hours. Forebrain progenitors were then treated with CHIR99021 (2 μM) + PD173074 (100 nM) for 72 hours, in the presence or absence of BMP4 (10 ng/mL) or BMPR inhibitors (DMH1 [250 nM] or LDN193189 [100 nM]). Gene expression is shown relative to reference gene *YWHAZ*; 100% indicates that a gene is expressed at the same level as *YWHAZ*. BMPRi: BMP receptor inhibitor. D) H1 hPSCs were differentiated into definitive ectoderm within 24 hours, and then anterior neural ectoderm within 24 hours, followed by forebrain progenitors within 48 hours. Forebrain progenitors were then treated with XAV939 (1 μM) in the presence or absence of increasing doses of HEDGEHOG pathway agonist 21K (0.1-5 nM) for 72 hours to generate ventral forebrain. This showed that high HEDGEHOG pathway activation, together with WNT inhibition, led to maximum expression of ventral forebrain markers. Gene expression is shown relative to the sample with the highest expression in this experiment. HH: Hedgehog pathway agonist. WNTi: WNT inhibitor. E) H1 hPSCs were differentiated into definitive ectoderm within 24 hours, and then posterior neural ectoderm within 24 hours, followed by hindbrain progenitors within 48 hours. Hindbrain progenitors were then treated with HEDGEHOG inhibitor (Vismodegib, 150 nM) and WNT agonist (CHIR99021, 2 μM) for 72 hours to generate dorsal forebrain, in the presence or absence of BMP agonist (BMP4, 10 ng/mL) or BMPR inhibitor (LDN193189, 100 nM). This showed that BMP inhibition promoted expression of dorsal forebrain markers *GSX2* and *PAX3*. Gene expression is shown relative to reference gene *YWHAZ*; 100% indicates that a gene is expressed at the same level as *YWHAZ*. BMPRi: BMP receptor inhibitor. F) H1 hPSCs were differentiated into definitive ectoderm within 24 hours, and then posterior neural ectoderm within 24 hours, followed by hindbrain progenitors within 48 hours. Hindbrain progenitors were then treated with HEDGEHOG pathway inhibitor Vismodegib (150 nM) in the presence or absence of increasing doses of WNT pathway agonist CHIR99021 (1-4 μM) for 72 hours to generate dorsal forebrain. This showed that moderate WNT pathway activation, together with HH inhibition, led to maximum expression of dorsal hindbrain markers *GSX2* and *OLIG3*. Gene expression is shown relative to reference gene *YWHAZ*; 100% indicates that a gene is expressed at the same level as *YWHAZ*. BMPRi: BMP receptor inhibitor. G) scRNAseq of H7 hPSC-derived day-4 hindbrain progenitors, and day-7 dorsal hindbrain or ventral hindbrain progenitors. Colors denote differentiation conditions. An asterisk (*) indicates that the legend is identical to that shown in Fig. 4c, and is reproduced here to aid the interpretation of other images shown in this subpanel. H) H1 hPSCs were differentiated into definitive ectoderm within 24 hours, and then posterior neural ectoderm within 24 hours, followed by hindbrain progenitors within 48 hours. Hindbrain progenitors were then treated with either Vismodegib (150 nM) + CHIR99021 (2 μM) + DMH1 (250 nM) for 72 hours to generate dorsal forebrain, or alternatively, 21K (5 nM) + XAV939 (1 μM) for 72 hours to generate ventral forebrain. Gene expression is shown relative to the sample with the highest expression in this experiment. I) H1 hPSCs were differentiated into definitive ectoderm within 24 hours, and then posterior neural ectoderm within 24 hours, followed by hindbrain progenitors within 48 hours. Hindbrain progenitors were then treated with WNT pathway inhibitor XAV939 (1 μM) in the presence or absence of increasing doses of HEDGEHOG pathway agonist 21K (50 pM-5 nM) for 72 hours to generate ventral hindbrain. This showed that high doses of HEDGEHOG agonist, together with WNT inhibitor, were required to maximally induce ventral hindbrain markers. Gene expression is shown relative to the sample with the highest expression in this experiment. WNT inh.: WNT inhibitor. qPCR data depicts the mean of two biological replicates, with s.e.m. shown. *In vitro* scRNAseq data is from a single experiment.

**Supplementary Figure 9:**
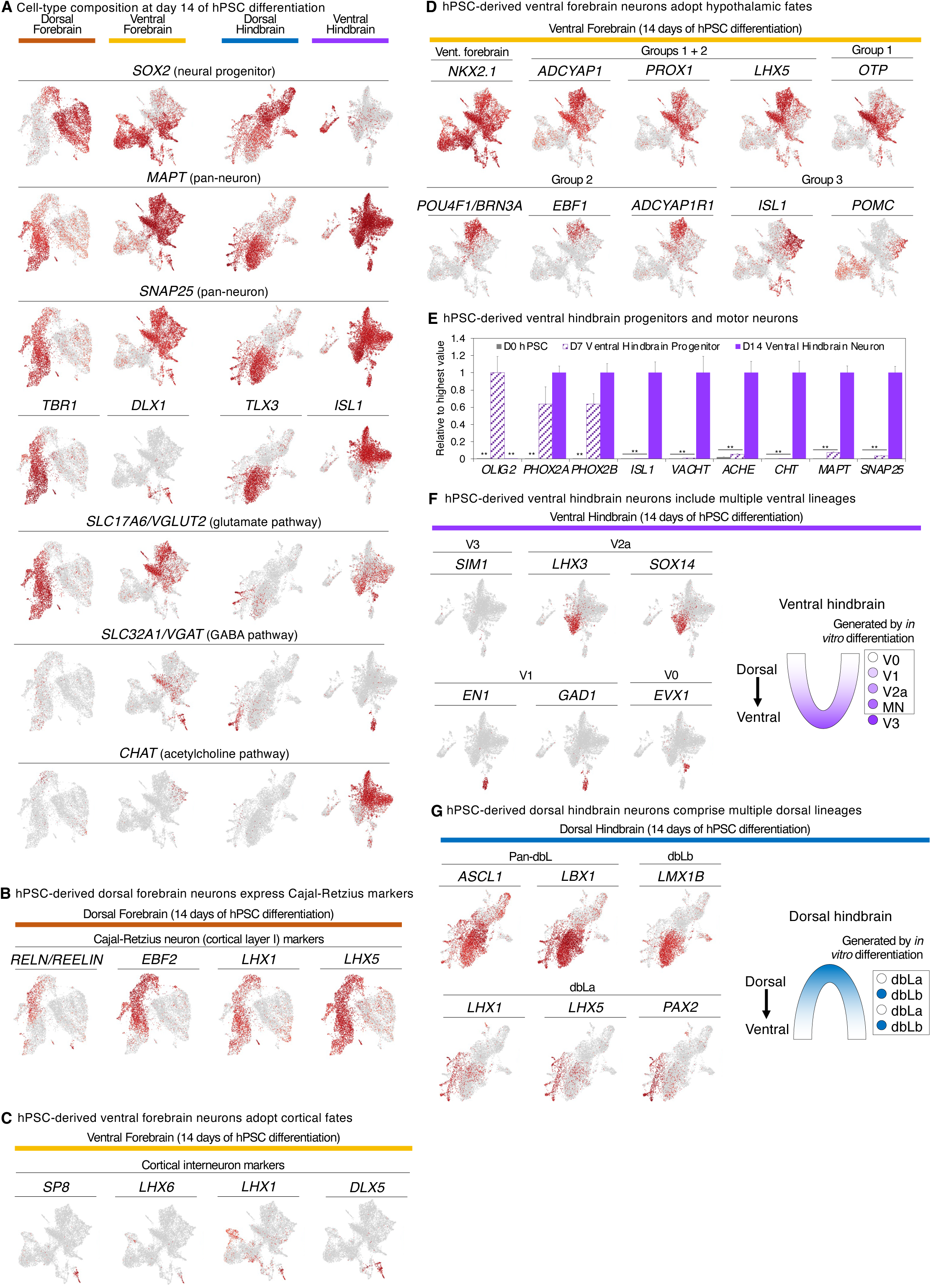
Differentiation of hPSCs into dorsal and ventral forebrain and dorsal and ventral hindbrain neurons within 14 days. A) scRNAseq of H7 hPSC-derived dorsal forebrain, ventral forebrain, dorsal hindbrain and ventral hindbrain populations on day 14 of differentiation. An asterisk (*) indicates images identical to those shown in Fig. 5d, which were reproduced here to aid the interpretation of other images shown in this subpanel. B) scRNAseq of H7 hPSC-derived dorsal forebrain populations on day 14 of differentiation reveals expression of Cajal-Retzius neuron markers ^208^. C) scRNAseq of H7 hPSC-derived ventral forebrain populations on day 14 of differentiation reveals expression of cortical interneuron markers. D) scRNAseq of H7 hPSC-derived ventral forebrain populations on day 14 of differentiation reveals expression of hypothalamus-associated markers. hPSC-derived groups I and II express *PROX1* ^209^, *LHX5* ^210^, and *ADCYAP1* ^140,141^, which are expressed *in vivo* by various regions of the hypothalamus, including the preoptic area, paraventricular nucleus, and supraoptic nucleus. hPSC-derived group I is additionally defined by *OTP*, the archetypic transcription factor required for the development of paraventricular and supraoptic nuclei ^211^. Instead, hPSC-derived group II is additionally defined by *POU4F1* (*BRN3A*) and *EBF1*, and may correspond to an early *Pou4f1*+ *Ebf1*+ *Pomc*-hypothalamic lineage *in vivo* ^212^. hPSC-derived group II also expresses *ADCYAP1R1* (the ADYCAP1 receptor), which is expressed by multiple brain regions, including the preoptic area, paraventricular nucleus, and supraoptic nucleus ^213^. Finally, hPSC-derived group III is demarcated by *ISL1* and *POMC*, which are expressed within the arcuate nucleus of the hypothalamus ^142^. E) qPCR of H1 hPSC-derived day-7 ventral hindbrain progenitors differentiated towards neurons for 7 additional days. Gene expression is shown relative to the sample with the highest expression in this experiment. F) scRNAseq of H7 hPSC-derived ventral hindbrain populations on day 14 of differentiation reveals expression of multiple ventral hindbrain neuron subtype markers (*left*), whose identities were extrapolated from ventral spinal cord neuron subtype markers ^214^. Cartoon of progenitor domains within the developing ventral hindbrain, based on embryological studies (*right*) ^214^. The box reflects the neuron subtypes putatively produced in our *in vitro* differentiation system. G) scRNAseq of H7 hPSC-derived dorsal hindbrain populations on day 14 of differentiation reveals expression of multiple dorsal hindbrain neuron subtype markers (*left*) ^132,145^. Cartoon of progenitor domains within the developing ventral hindbrain, based on embryological studies (*right*) ^132,145^. The box reflects the neuron subtypes putatively produced in our *in vitro* differentiation system. *In vitro* scRNAseq data is from a single experiment. qPCR data depicts the mean of two biological replicates, with s.e.m. shown.

**Supplementary Figure 10:**
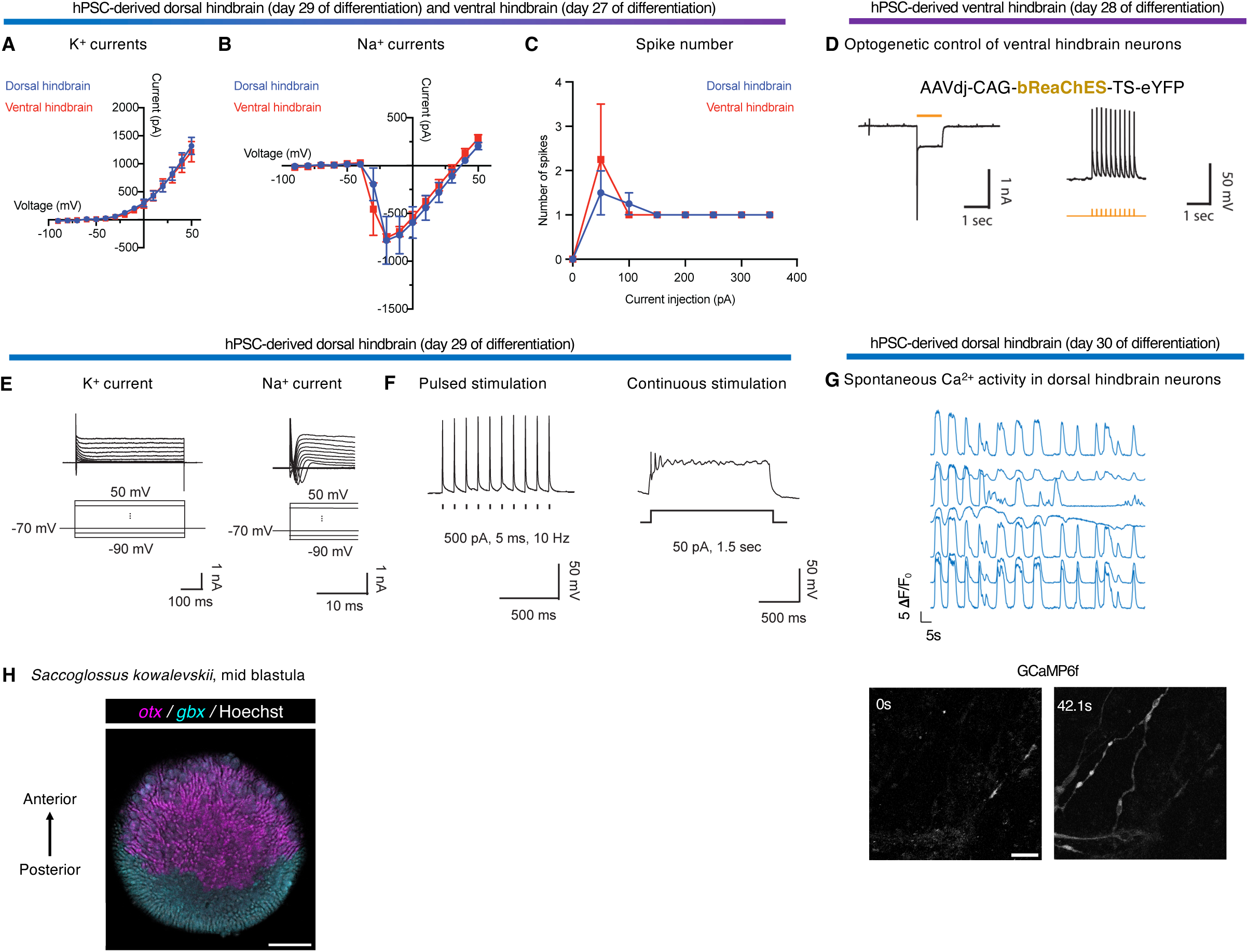
Functional assessment of hPSC-derived dorsal and ventral hindbrain neurons, and evolutionary conservation of anterior and posterior ectoderm. A) Summary of the current-voltage relationship of voltage-dependent K^+^ currents from dorsal and ventral hindbrain neurons. Currents were measured from -90 mV to +50 mV holding potentials. n (number of cells) = 4. Data are mean ± S.E.M. B) Summary of the current-voltage relationship of voltage-dependent Na^+^ currents from dorsal and ventral hindbrain neurons. Currents were measured from -90 mV to +50 mV holding potentials. n (number of cells) = 4. Data are mean ± S.E.M. C) Summary of the number of spikes elicited by prolonged current injection. D) Day 28 H1 hPSC-derived ventral hindbrain neurons transduced with AAVdj-*CAG-bReaChES-eYFP* were stimulated with either prolonged (1 second), or pulses of (5 ms, 5 Hz), 560nm light, and electrophysiological recording was performed to detect action potentials. E) Electrophysiological activity of H1 hPSC-derived dorsal hindbrain neurons on day 30 of differentiation, showing voltage-dependent K^+^ (left) or Na^+^ (right) channel currents elicited by depolarization of the holding potential from -90 mV to 50 mV in voltage-clamp mode. F) Current-clamp characterization of H1 hPSC-derived dorsal hindbrain neurons on day 30 of differentiation, showing action potentials elicited by injection of pulsed (500 pA, 5 ms, 10 Hz, left) or prolonged (50 pA, 1.5 seconds, right) currents in current-clamp mode. G) Live Ca^2+^ imaging of WTC11 *AAVS1-CAG-GCaMP6f* hPSCs differentiated into dorsal hindbrain neurons for 30 days, which exhibited spontaneous Ca^2+^ transients as assessed within individual cells (left) and across the culture (right). Scale bar = 50 μm. H) *In situ* staining of acorn worm embryo at the mid blastula stage. A maximum intensity projection of optical sections focusing on the ectoderm is shown. Scale bar = 100 μm. Ca^2+^ imaging and electrophysiology were each performed on a single experiment. *In vivo* staining was performed on a single embryo.

## Materials and Methods

### Mouse models

*Gbx2^CreER/+^* mice^174^ were originally developed by, and generously provided by, James Li’s laboratory (JAX #022135) and maintained on a CD1 background, heterozygous for the targeted allele. Mice carrying the *Gbx2^CreER^*allele were identified by PCR as described by Jackson Laboratory.

*Sox2^CreER/+^* mice^116^ were originally developed by Konrad Hochedlinger’s laboratory, were obtained from Jackson Laboratory (JAX #017593), and maintained on a C57BL/6 background, heterozygous for the targeted allele. Mice carrying the *Sox2^CreER^* allele were identified by PCR as described by Jackson Laboratory.

*ROSA26-CAG-LoxP-STOP-LoxP-tdTomato* (“Ai14”) mice^111^ were originally developed by the Allen Brain Institute, were obtained from Jackson Laboratory (JAX #007914), and were maintained homozygous for the targeted allele.

*ROSA26-Confetti* mice^175^ were originally developed by Hans Clevers’s laboratory, were obtained from Jackson Laboratory (JAX #017492), and were respectively maintained as homozygous for the targeted allele.

*Tcf/Lef:H2B-GFP* mice^173^ were originally developed by Anna-Katerina Hadjantonakis’s laboratory, were obtained from the Jackson Laboratory (#013752), and maintained on a CD1 background, heterozygous for the targeted allele. These mice were shared by Teni Anbarchian and Roel Nusse.

### Mouse husbandry and lineage tracing

Mice of the desired genetic backgrounds were mated, leading to timed pregnancies. Noon on the day a vaginal plug was detected was designated as E0.5. For Gbx2^CreER^-mediated lineage tracing, 1 mg of (*Z*)-4-Hydroxytamoxifen (henceforth referred to as 4OHT; Sigma, H7904-25MG) was administered. For Sox2^CreER^-mediated lineage tracing, 0.5 mg of 4OHT was administered.

4OHT was resuspended and administered as described previously ^115^. In brief, a 20 mg/mL stock of 4OHT was made by dissolving 25 mg of 4OHT powder in 1250 μL of ethanol. This solution was heated for 10 minutes at 60°C and sonicated briefly for 10-20 seconds. This 20 mg/mL stock of 4OHT was aliquoted to generate 50 μL aliquots (i.e., 1 mg per aliquot), which were stored at -20°C.

When ready to inject, a 50 μL aliquot of 4OHT was thawed at 65°C for 10 minutes. During the thawing process, the solution was vortexed and quickly centrifuged in a microcentrifuge every 2-3 minutes. Simultaneously, 250 μL of corn oil (Sigma, C8267-500mL) was warmed to 65°C. After the 4OHT solution was thawed for 10 minutes, 250 μL of warm corn oil was then added to 4OHT solution, thereby generating a final concentration of 3.33 mg/mL of 4OHT. This mixture was heated at 65°C for 45 minutes, with vortexing every 15 minutes. This 3.33 mg/mL of 4OHT solution was delivered via intraperitoneal injection, using a U-100 Insulin Syringe (Comfort Point, 26028), to pregnant female mice at the relevant labeling timepoint.

Mouse embryos of specific ages were obtained from pregnant mice. The biological sex of mouse embryos was not determined.

### *In situ* hybridization of whole mount mouse embryos

Hybridization chain reaction v3.0 (HCR3) ^176^ was used to perform whole-mount fluorescent *in situ* hybridization (FISH) of mouse embryos. Embryos were dissected in ice-cold 4% paraformaldehyde and subsequently fixed either for 1 hour (for E8.0 and earlier embryonic stages), or alternatively overnight (for E8.25 and later embryonic stages), prior to methanol dehydration. Hybridization mRNA probes, amplifiers, and buffers were obtained from Molecular Instruments, and the protocol for whole-mount staining of mouse embryos were followed as per the manufacturer’s instructions with the following modifications: Proteinase K digestions varied with embryonic stage; E6.5: 5 min; E7.5: 7.5 min; E8.5: 10 min; E9.5: 15 min. Embryos were incubated in SSCT + DAPI prior to mounting.

### Immunostaining of whole mount mouse embryos

Embryos were dissected in ice-cold PBS and fixed overnight on a rocker at 4C in 1% paraformaldehyde. They were washed 3 x 10 min in PBS at room temperature before being blocked overnight in embryo blocking buffer (3% BSA, 0.2% Tween-20 in PBS). Embryos were then incubated with primary antibodies (diluted in embryo blocking buffer) for 48 hours at 4C with gentle rocking. Subsequently, the embryos were washed for 4 hours in PBST (0.2% Tween-20 in PBS) before being transferred to incubation with secondary antibodies for 48 hours at 4C with gentle rocking. Embryos were cleared overnight in Rapiclear 1.49 (Sunjin labs) before being mounted, also in Rapiclear 1.49.

### Imaging of mouse embryos

Images were collected at room temperature on Olympus FV3000, Leica SP8, or Zeiss LSM980 confocal microscopes using 405, 488, 514, 561, and 640 nm wavelengths and 4X/0.16, 10X/0.4, and 20X/0.75 dry objectives, and 40X/1.3 oil objective. Data was displayed with FIJI ImageJ software^177^.

### Computational analysis of mouse embryo images in which neural ectoderm cells were sparsely labeled

*Sox2-CreER;Confetti* mouse embryo images were imported into Imaris 10 for quantification. Sox2+ neural ectoderm-derived cell clusters were identified in CFP, YFP, and RFP channels separately using the Spots function. Cell clusters were identified as spots within 50 μm of each other in the same channel using the Imaris SplitSpots plugin function. A surface feature of the forebrain and midbrain were generated from the Otx2 immunofluorescence signal in the far-red channel. Cell clusters were defined relative to the forebrain/midbrain surface element. Sox2+ neural ectoderm-derived cell clusters with less than 2 cells across the midbrain-hindbrain boundary were assigned to the region in which the majority of cells resided.

### Computational analysis of differentiated hPSC immunofluorescence images

Immunofluorescence images of differentiated hPSCs were quantified using either Fiji ^177^ or Imaris 9 to determine the percentage of marker-positive cells. The DAPI channel was used to define the nucleus of each individual cell, which was then defined as marker-positive or marker-negative using a threshold.

Maximum intensity projections of immunostaining images were imported into Fiji^177^ for cell quantification. A threshold of marker intensity was manually set to define marker-positive and marker-negative cells based on negative controls (i.e., cells that did not express the protein of interest). Subsequently, to define the border of each individual nucleus, we used the DAPI channel and employed the Analyze Particles function in Fiji, using specific size and circularity values that we kept consistent within each image. After the border of each individual nucleus was defined, the Analyze Particles function was used to determine whether the nucleus was marker-positive or marker-negative, in relation to the aforementioned marker intensity threshold.

Separately, Z-stacks of immunostaining images were imported into Imaris 9 to generate 3-dimensional models. For all fluorescence channels, we performed background subtraction via auto-thresholding to reduce noise. We then analyzed the DAPI channel, filtered the DAPI channel via the Intensity Mean function, and the Spots function was used to define the border of each individual nucleus, with a 7 μm radius for each nucleus in 3-dimensional space. To define whether each nucleus was marker-positive or marker-negative, we applied the Intensity Mean function on the fluorescence of each given marker. A minimum value for the Intensity Mean function was manually determined. The same threshold was used for a given marker across all cell-types analyzed.

### Cell culture

All cells in this study were cultured in standard incubator conditions (20% O_2_, 5% CO_2_ and 37°C).

### Basement membrane matrix for cell culture experiments

hPSCs were maintained and differentiated on cell culture plates that been pre-coated with Geltrex basement membrane matrix, as described previously ^178^. Geltrex (Thermo Fisher) was diluted 1:100 in DMEM/F12 (Thermo Fisher) and was kept at 4 °C. To coat cell culture plates, Geltrex solution was added to cell culture plastics for at least 1 hour at 37 °C. The volume of Geltrex solution added was roughly equivalent to half the working volume of the well or dish (e.g., 1 mL or 0.5 mL of Geltrex solution was added per well of a 6-well or 12-well plate, respectively). After coating with Geltrex, the basement membrane solution was aspirated, leaving behind a thin film; subsequently, cells were plated on the Geltrex-coated cell culture plastics.

A detailed protocol to coat cell culture plastics with Geltrex is available ^179^.

### Human pluripotent stem cell culture

Multiple hPSC lines, including both human embryonic stem cells (hESCs) and human induced pluripotent stem cells (hiPSCs) were used in this study: wild-type H1, H7, and H9 hESCs ^180^ (WiCell), wild-type SUN004.1 and SUN004.2 hiPSCs ^178^ (provided by Hiromitsu Nakauchi’s laboratory), HES3 *MIXL1-GFP* hPSCs ^181^ (provided by Andrew Elefanty’s, Edouard Stanley’s, and Elizabeth Ng’s laboratories), and WTC11 *AAVS1-CAG-GCaMP6f* hiPSCs ^182^ (provided by Bruce Conklin’s laboratory). Of note, SUN004.1 and SUN004.2 represent two different, clonally-derived hiPSC lines reprogrammed from peripheral blood mononuclear cells obtained from the same healthy adult volunteer (SUN004)^178^.

Undifferentiated hPSCs were maintained as described previously ^178^ in either mTeSR1 or mTeSR Plus media (StemCell Technologies) supplemented with 1% penicillin/streptomycin (Thermo Fisher), on Geltrex-coated cell culture plates. When the undifferentiated hPSC cultures became partially confluent, undifferentiated hPSCs were passaged by treating them with EDTA (Versene, Thermo Fisher) for 7 minutes at room temperature. After removal of EDTA, mTeSR was re-added and the plate was manually scraped to generate clumps, which were reseeded onto new Geltrex-coated plates.

A detailed protocol to culture and passage undifferentiated hPSCs is available ^179^.

### hPSC differentiation into ectodermal derivatives

The day before beginning differentiation (“day -1” [D-1]), hPSCs were enzymatically dissociated using Accutase (Thermo Fisher) and plated as single cells from 30,000-100,000 cells/cm^2^ on Geltrex-coated cell culture plates in mTeSR or mTeSR Plus + 1% penicillin/streptomycin + ROCK inhibitor thiazovivin (1 μM, Tocris). Inclusion of ROCK inhibitor was critical to ensure survival of single cells during seeding.

After 24 hours, cells were ready for subsequent differentiation. The first 7 days of hPSC differentiation were conducted in Chemically Defined Medium 2 (CDM2), whose composition has been described previously ^73,74^. CDM2 comprises 50% IMDM + GlutaMAX (Thermo Fisher, 31980-097), 50% F12 + GlutaMAX (Thermo Fisher), polyvinyl alcohol (1 mg/mL, Sigma) + chemically defined lipid concentrate (1% v/v, Thermo Fisher), 1-thioglycerol (450 μM, Sigma), recombinant human insulin (0.7 μg/mL, Sigma), human transferrin (15 μg/mL, Sigma), penicillin/streptomycin (1% v/v, Thermo Fisher). To create CDM2, polyvinyl alcohol was brought into suspension by gentle warming and magnetic stirring, and the media was filtered through a 0.22 μm filter prior to use.

A detailed protocol to passage hPSCs for differentiation, and to prepare CDM2 basal medium, is available ^179^.

#### Definitive ectoderm (D0-1, 24 hours)

D0 hPSCs were briefly washed before differentiation with DMEM/F12 to remove any traces of mTeSR Plus. They were then differentiated toward definitive ectoderm in CDM2 media supplemented with A8301 (1 μM, Tocris), Trametinib (250 nM, Cellagen Technology), C59 (1 μM, Tocris), and LDN193189 (100 nM, Stemgent) for 24 hours. Subsequently, D1 definitive ectoderm was used for either anterior neural ectoderm, posterior neural ectoderm, border ectoderm, or surface ectoderm induction.

#### Anterior neural ectoderm (D1-2, 24 hours)

D1 definitive ectoderm cells were resplit using Accutase (Thermo Fisher) and plated as single cells on Geltrex-coated cell culture plates in CDM2 supplemented with A8301 (1 μM, Tocris), LDN193189 (100 nM, Stemgent), C59 (1 μM, Tocris), and Thiazovivin (1 μM, Tocris) for 24 hours. Thiazovivin was added to promote cell survival after enzymatic passaging. For experiments ending after two days of differentiation, cells were plated at 37,500 cells/cm^2^, and for experiments ending after more than 2 days of differentiation (i.e., towards a forebrain fate), 75,000 cells/cm^2^ were plated. After 24 hours of anterior neural ectoderm media, cells were used for either forebrain or midbrain induction.

#### Posterior neural ectoderm (D1-2, 24 hours)

D1 definitive ectoderm cells were resplit using Accutase (Thermo Fisher) and plated as single cells on Geltrex-coated cell culture plates in CDM2 supplemented with A8301 (1 μM, Tocris), LDN193189 (100 nM, Stemgent), TTNPB (50 nM, Tocris), FGF2 (20 ng/mL, R&D Systems), XAV939 (1 μM, Tocris), and Thiazovivin (1 μM, Tocris) for 24 hours. Thiazovivin was added to promote cell survival after enzymatic passaging. For experiments ending after two days of differentiation, cells were plated at 37,500 cells/cm^2^, and for experiments ending after more than 2 days of differentiation (i.e., toward a hindbrain fate), 150,000 cells/cm^2^ were plated. After 24 hours of posterior neural ectoderm media, cells were used for hindbrain induction.

#### Border ectoderm (D1-2, 24 hours)

D1 definitive ectoderm cells were resplit using Accutase (Thermo Fisher) and plated as single cells at 37,500 cells/cm^2^ on Geltrex-coated cell culture plates in CDM2 supplemented with A8301 (1 μM, Tocris), PD0325901 (100 nM, Tocris), CHIR99012 (3 μM, Tocris), BMP4 (0.5 ng/mL, R&D Systems), and Thiazovivin (1 μM, Tocris) for 24 hours. Thiazovivin was added to promote cell survival after enzymatic passaging.

#### Surface ectoderm (D1-2, 24 hours)

D1 definitive ectoderm cells were resplit using Accutase (Thermo Fisher) and plated as single cells at 37,500 cells/cm^2^ on Geltrex-coated cell culture plates in CDM2 supplemented with A8301 (1 μM, Tocris), PD0325901 (100 nM, Tocris), BMP4 (30 ng/mL, R&D Systems), C59 (1 μM, Tocris), and Thiazovivin (1 μM, Tocris) for 24 hours. Thiazovivin was added to promote cell survival after enzymatic passaging.

#### Forebrain (D2-4, 48 hours)

D2 anterior neural ectoderm cells were briefly washed with DMEM/F12 prior to addition of forebrain media. D3 forebrain media consisted of CDM2 supplemented with XAV939 (1 μM, Tocris), LDN193189 (100 nM, Stemgent), and FGF2 (20 ng/mL, R&D Systems), and was changed after 24 hours into D4 forebrain media, which consisted of CDM2 supplemented with XAV939 (1 μM) and LDN193189 (100 nM). Subsequently, forebrain cells could be further differentiated toward either dorsal forebrain or ventral forebrain fates.

#### Midbrain (D2-4, 48 hours)

D2 anterior neural ectoderm cells were briefly washed with DMEM/F12 prior to addition of midbrain media. D3 midbrain media consisted of CDM2 supplemented with CHIR99021 (3 μM, Tocris) and was changed after 24 hours into D4 midbrain media, which consisted of CDM2 supplemented with CHIR99021 (3 μM) and FGF2 (20 ng/mL, R&D Systems).

#### Hindbrain (D2-4, 48 hours)

D2 posterior neural ectoderm cells were briefly washed with DMEM/F12 prior to addition of hindbrain media. D3 and D4 hindbrain media consisted of CDM2 supplemented with CHIR99021 (0.75 μM, Tocris) and FGF2 (20 ng/mL, R&D Systems).

Hindbrain media was replaced after 24 hours. Subsequently, hindbrain cells could be further differentiated toward either dorsal hindbrain or ventral hindbrain fates.

#### Dorsal Forebrain (D4-7, 72 hours)

D4 forebrain progenitors were briefly washed with DMEM/F12 prior to addition of dorsal forebrain media. D5, D6, and D7 dorsal forebrain media consisted of CDM2 supplemented with CHIR99021 (2 μM, Tocris), Vismodegib (150 nM, Selleck Chemicals), and PD173074 (100 nM, Tocris). Dorsal forebrain differentiation was conducted for 72 hours, and dorsal forebrain media was replaced every 24 hours.

#### Ventral Forebrain (D4-7, 72 hours)

D4 forebrain progenitors were briefly washed with DMEM/F12 prior to addition of ventral forebrain media. D5, D6, and D7 dorsal forebrain media consisted of CDM2 supplemented with XAV939 (1 μM, Tocris) and SAG 21K (5 nM, Tocris). Ventral forebrain differentiation was conducted for 72 hours, and ventral forebrain media was replaced every 24 hours.

#### Dorsal Hindbrain (D4-7, 72 hours)

D4 hindbrain progenitors were briefly washed with DMEM/F12 prior to addition of dorsal hindbrain media. D5, D6, and D7 dorsal hindbrain media consisted of CDM2 supplemented with CHIR99021 (2 μM, Tocris), Vismodegib (150 nM, Selleck Chemicals), and LDN193189 (100 nM, Tocris). Dorsal hindbrain differentiation was conducted for 72 hours, and dorsal hindbrain media was replaced every 24 hours.

#### Ventral Hindbrain (D4-7, 72 hours)

D4 hindbrain progenitors were briefly washed with DMEM/F12 prior to addition of ventral hindbrain media. D5, D6, and D7 ventral hindbrain media consisted of CDM2 supplemented with XAV939 (1 μM, Tocris) and SAG 21K (5 nM, Tocris). Ventral hindbrain differentiation was conducted for 72 hours, and ventral hindbrain media was replaced every 24 hours.

#### Neuronal differentiation (D7-14, 7 days)

hPSC-derived day 7 dorsal forebrain, day 7 ventral forebrain, day 7 dorsal hindbrain, and day 7 ventral forebrain progenitors were subsequently differentiated into neurons for 7 days. Neuronal differentiation media was modified from a previous report ^139^. From day 8-14, cells were differentiated in BDNF (20 ng/ml, PeproTech), GDNF (20 ng/ml, PeproTech), Forskolin (10 μM, Tocris), 2-Phospho-L-ascorbic acid (200 μg/ml, Sigma-Aldrich), and DAPT (10 μM, R&D Systems), with different basal media used on each day of differentiation. The general approach was to gradually transition differentiating cells from CDM2 to Neurobasal differentiation media. Media composition was as follows: 1) D7-8 media (i.e., first 24 hours of neuronal differentiation) contained: 25% Neurobasal media (see below) + 75% CDM2 media was added; 2) D8-9 media (i.e., second 24 hours of neuronal differentiation): 50% Neurobasal media + 50% CDM2 media was added; 3) D9-10 (i.e., third 24 hours of neuronal differentiation), 75% Neurobasal media + 25% CDM2 media was added; 4) D10-14 (i.e., remaining 96 hours of neuronal differentiation), 100% Neurobasal media. In other words, throughout the entire neuronal differentiation phase, cells were exposed to the same concentration of neuronal differentiation signals (BDNF, GDNF, Forskolin, 2-Phospho-L-ascorbic acid, and DAPT) while a gradual change in basal media occurred. “Neurobasal media” is comprised of Neurobasal media (Thermo Fisher) supplemented with 1% v/v N2 (Thermo Fisher), 2% v/v B27 w/o Vitamin A (Thermo Fisher), and 1% v/v penicillin/streptomycin (Thermo Fisher). The media was passed through a sterilizing 0.22 μM filter before use.

### hPSC differentiation into non-ectodermal cell-types

As a positive control to achieve MIXL1-GFP expression, hPSCs were differentiated into mid primitive streak as described previously ^78^, by culturing them in Activin A (30 ng/mL) + BMP4 (40 ng/mL) + CHIR99021 (6 μM) + FGF2 (20 ng/mL) in CDM2 basal media for 24 hours.

To generate definitive endoderm and paraxial mesoderm, hPSCs were first differentiated into anterior primitive streak as described previously ^78^, by culturing them in Activin A (30 ng/mL) + CHIR99021 (4 μM) + FGF2 (20 ng/mL) in CDM2 basal media for 24 hours. hPSC-derived anterior primitive streak was then briefly washed with DMEM/F12 and then treated with either definitive endoderm-inducing media ^77^ (Activin A [100 ng/mL] + LDN193189 [250 nM] in CDM2 basal media) for 24 hours, or paraxial mesoderm-inducing media ^78^ (A-83-01 [1 μM] + LDN193189 [250 nM] + CHIR99021 [3 μM] + FGF2 [20 ng/mL]) for 24 hours.

### Fluorescent labeling of hPSC-derived cells for coculture experiments

Day 2 anterior and posterior neural ectoderm (aNE and pNE, respectively) were generated as described above, and then associated using Accutase (Thermo Fisher). aNE cells were kept on ice, while pNE cells were resuspended at 10^6^ cells/mL in prewarmed PBS containing 1-5 μM ViaFluor 488 Cell Proliferation dye (Biotium) for 10 minutes at 37 °C. Dye was prepared on the day of the labeling. Cells were washed with DMEM/F12 (Thermo Fisher) prior to pelleting. For mixed cultures, aNE and dye-labeled pNE were mixed at a 1:1 ratio prior to plating and subsequent culture in either forebrain, midbrain, or hindbrain media, as described. After 2-4 days of culture in either forebrain, midbrain, or hindbrain media, cells were fixed and stained as described above, prior to imaging.

### Quantitative PCR

Cells were lysed with RNeasy Lysis Buffer (Qiagen) and RNA was subsequently extracted using the RNeasy Plus Kit (Qiagen) as per the manufacturer’s instructions. RNA was reverse transcribed into cDNA using the High-Capacity cDNA Reverse Transcription Kit (Applied Biosystems). qPCR was subsequently performed on the cDNA with the SensiFAST SYBR Lo-ROX Kit (Bioline) on a QuantStudio 5 qPCR machine (Thermo Fisher). qPCR primer sequences are provided in **Table S1**. The cycling parameters were as follows: initial dissociation (95 °C, 2 mins) followed by 40 cycles of amplification and SYBR signal detection (95 °C dissociation, 5 seconds; 60 °C annealing, 10 seconds; followed by 72 °C extension, 30 seconds), with a final series of steps to generate a dissociation curve at the end of each run. During qPCR data analysis, the fluorescence threshold to determine Ct values was set at the linear phase of amplification. Expression of all genes was normalized to the levels of the reference gene *YWHAZ*.

In histograms that depict qPCR data, the y-axis label of “% *YWHAZ*” indicates that the expression level of each gene is depicted relative to the expression of reference gene *YWHAZ*, such that “100%” indicates that the gene is expressed at the same level of *YWHAZ*. Alternatively, other histograms depicting qPCR data carry a y-axis label of “Expression relative to highest value”, which indicates that the expression of each gene is depicted relative to the biological sample with the highest value of that gene in a given experiment.

qPCR histograms depict the mean of two biological replicates, defined as two independent wells treated with the same differentiation media in a single experiment. Two technical qPCR replicate measurements were performed on each biological replicate. Error bars represent the standard error of the mean (s.e.m.).

qPCR primer sequences used to target human genes are as follows:

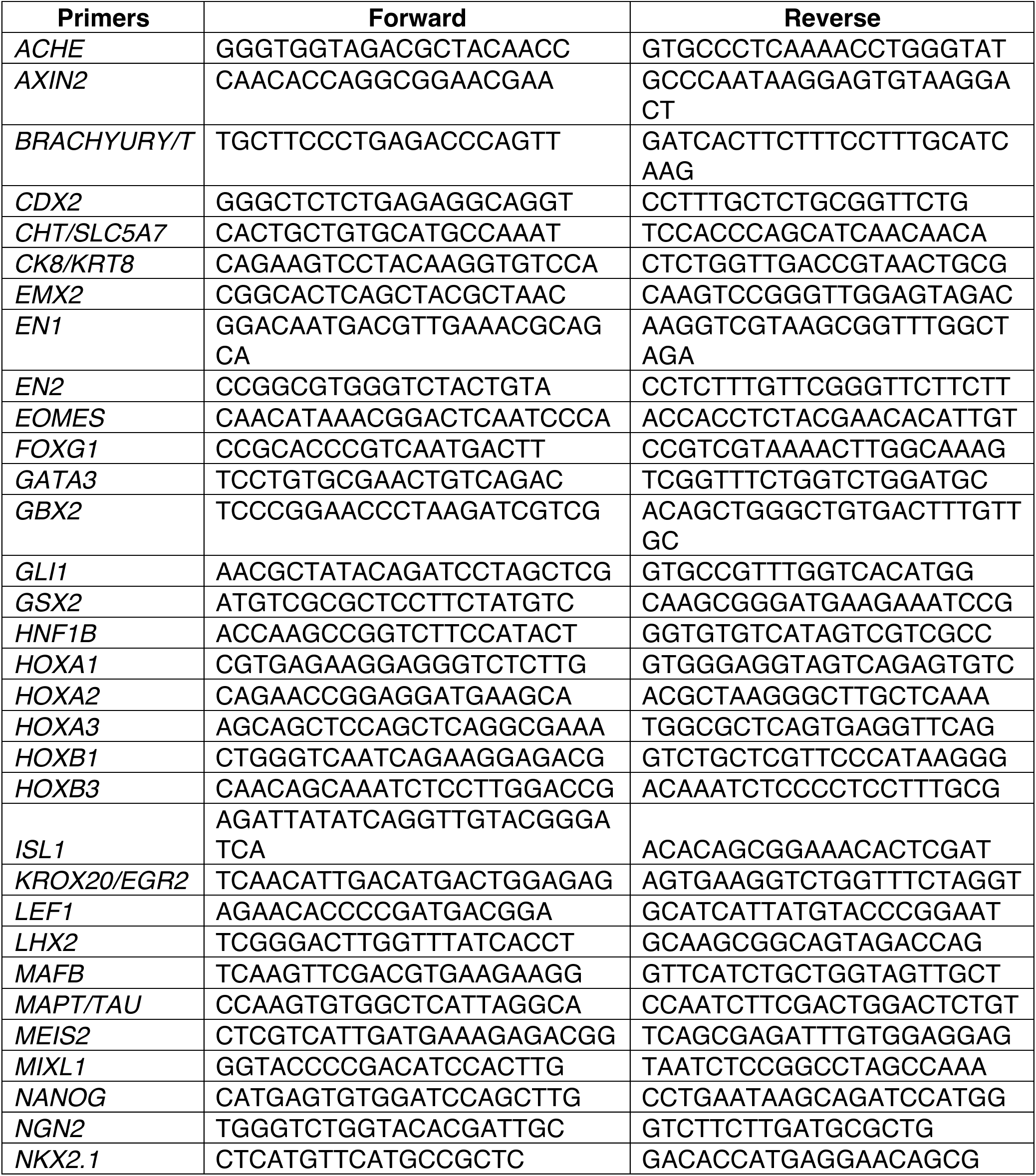
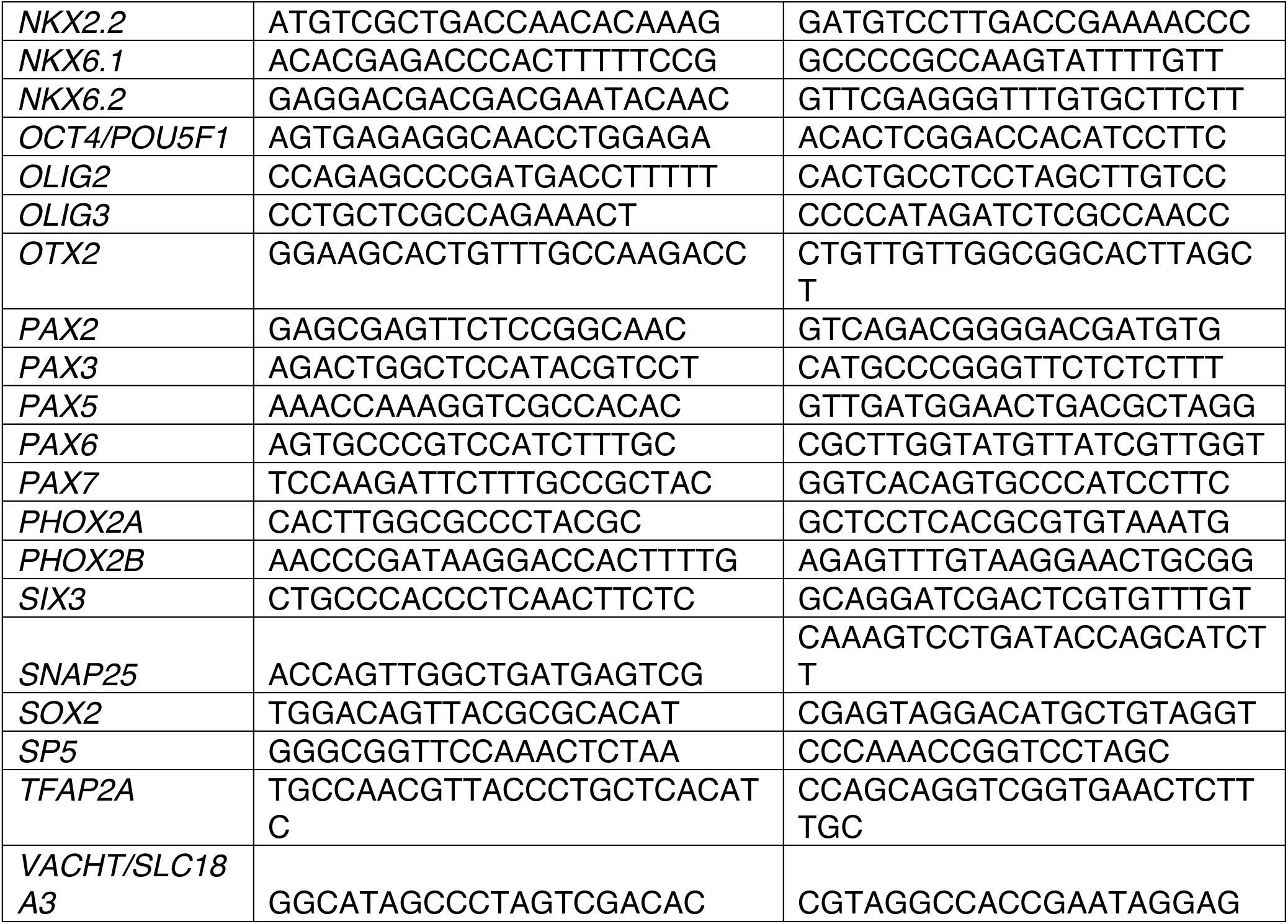

### Immunocytochemistry of hPSC-derived cell-types

Adherent cells were washed once with PBS, fixed for 10 minutes in 4% paraformaldehyde (Fisher Scientific) at room temperature, and then washed three more times with PBS. They were subsequently permeabilized with permeabilization/blocking solution (0.1% Triton-X100 + 5% donkey serum + PBS) for 1 hour at 4 °C. Subsequently, primary antibodies were added in permeabilization/blocking solution overnight at 4 °C. Cells were then washed three times with PBS, and secondary antibodies were added in permeabilization/blocking solution for 1 hour at 4 °C. Cells were then washed twice with PBS, and then nuclear counterstaining was performed with 0.2 μg/mL DAPI + PBS for 10 minutes at room temperature. Imaging was conducted on an Olympus FV3000 confocal microscope.

### *In situ* hybridization of hPSC-derived cell-types

*In situ* hybridization of hPSC-derived anterior and posterior neural ectoderm was performed using hybridization chain reaction v3.0 (HCR3)^176^, in accordance with the Molecular Instruments protocol for cultured mammalian cells. In brief, DNA probe sets were generated by Molecular Instruments, Inc., targeting the RNA sequences of the *Mus musculus* genes *Otx2, Gbx2*, and *Sox2*. In brief after generating hPSC-derived cell types, cells were fixed with 4% formaldehyde, watched twice with PBS, and permeabilized overnight at -20 degrees Celsius in 70% ethanol. Cells were then washed twice with 5x sodium chloride sodium citrate plus 0.1% Tween20 (SSCT), and prehybridized in probe hybridization buffer (from Molecular Instruments). The probe solution was prepared by adding 1.2 pmol of each probe set (*Otx2, Gbx2,* and *Sox2*) to 300μL of probe hybridization buffer at 30 degrees Celsius, and added to cells after removing the pre-hybridization buffer. Samples were then incubated overnight at 37 degrees Celsius. Cells were then washed four times with probe wash buffer (from Molecular Instruments) and twice with SSCT. Cells were pre-amplified with 300μL of amplification buffer (from Molecular Instruments). Separately, 18 pmol of hairpin h1 and 18 pmol of hairpin h2 (B1-647, B2-546, and B3-488) were prepared by snap cooling 6μL of 3μM stock. Pre-amplification buffer was then removed from samples and replaced with the hairpin solution. Samples were incubated overnight in the dark at room temperature. Samples were washed 5 times with SSCT and prepared for confocal imaging.

### Western Blot

hPSC-derived anterior and posterior neural ectoderm cells were dissociated with TrypLE Express, centrifuged at 1400 rpm for 5 minutes, washed in PBS, and then centrifuged again. Cells were then incubated on ice in RIPA buffer for 30-60 minutes to lyse them. RIPA buffer is composed of: 0.05 M Tris-HCL pH 7.5 (Thermo Fisher Scientific, 15567027), 0.15M NaCl (Thermo Fisher Scientific, J60434AE), 0.1% SDS (Thermo Fisher Scientific, 28364), 0.5% sodium deoxycholate (Millipore Sigma, D6750), 0.1% Triton X-100 (Millipore Sigma, T8787), 1:50 protease inhibitor (Millipore Sigma, 11697498001), and 1:10,000 benzonase nuclease (Millipore Sigma, E1014-25KU).

This whole cell lysate was then centrifuged at 12,000g for 10 minutes at 4 °C to remove debris from protein samples. Protein concentration was then quantified using the Pierce bicinchoninic acid (BCA) method (Pierce, 23227).

For each sample, 10 μg of total protein was combined with gel-loading dye (Boster Biological Technology, AR1112), boiled at 99 °C for 5 minutes, briefly centrifuged, and then loaded on a 4-12% Bis-Tris gel (Thermo Fisher Scientific, NW04125BOX). In another lane of the same gel, a protein ladder (Thermo Fisher Scientific, 26620) was added. Electrophoresis was conducted for 30 minutes at 200 volts in MOPS buffer (Thermo Fisher Scientific, B0001).

After electrophoresis, proteins were transferred to a nitrocellulose membrane using the Mini Blot Module (Thermo Fisher Scientific, B1000) for 1 hour at 15 volts. The nitrocellulose membrane was then blocked in blocking solution (PBS [Thermo Fisher, 10010023] + 5% milk [Bio-Rad, 1706404] + 0.1% Tween-20 [Sigma, 11332465001]). The blot was probed with primary antibodies for GBX2 (Thermo Fisher, PA5-66953, diluted 1:1000) and VINCULIN (Cell Signaling Technology, E1E9V, diluted 1:1000), diluted in blocking solution. Primary antibodies were incubated overnight at 4 °C on a shaker. Blots were then washed three times for 5 minutes each with PBS + 0.1% Tween-20 on a shaker. Subsequently, the blot was incubated with horseradish peroxidase (HRP)-conjugated secondary antibodies (Cell Signaling Technology, 7074, diluted 1:500), diluted in blocking solution. Secondary antibodies were incubated for 1 hour at room temperature on a shaker. Finally, blots were then washed three times for 5 minutes each with PBS + 0.1% Tween-20 on a shaker. Proteins were visualized using chemiluminescence substrates (Millipore Sigma, WBLUF0500) and imaged on a ChemiDoc imager (Bioradiations, 170-8280).

### Flow cytometry

hPSCs and hPSC-derived ectoderm cells were dissociated by incubating them in TrypLE Express (Thermo Fisher) for 5 minutes at 37 °C. The resulting cell suspension was diluted 1:10 in FACS Buffer, which comprised 0.5% BSA (Thermo Fisher) and 2 mM EDTA (Thermo Fisher) in PBS, and subsequently pelleted at 300 x *g*.

Intracellular protein staining for flow cytometry was conducted using the Inside Stain Kit (Miltenyi Biotec) as per manufacturer’s instructions.

Alternatively, live cells were analyzed, in which case no fixation or permeabilization was used.

Flow cytometry was performed on a Beckman Coulter CytoFlex analyzer (Stanford Stem Cell Institute FACS Core). For data analysis, cells were gated in FlowJo software based on forward and side scatter area parameters, followed by height and width parameters for doublet discrimination. An example of the flow cytometry gating strategy used for live cells is shown in **Supplementary Fig. 1**.

### Bulk RNA-sequencing of hPSC-derived cell-types

Cells were lysed with RNeasy lysis buffer (Qiagen) and RNA was subsequently extracted using the RNeasy Plus Kit (Qiagen) as per the manufacturer’s instructions. RNA-seq libraries were generated by Novogene and sequenced on an Illumina NovaSeq 6000.

Read quality control was performed using FastQC^183^. Subsequently, index adaptors and low-quality base-calls were removed using TrimGalore. RNA-seq reads were aligned to human reference genome hg38 using Kallisto^184^. Gene-level counts were derived from transcript-level counts using tximport^185^.

BAM files were converted to bigwig files, which were then averaged across samples for visualization, using deepTools^186^. RNA-seq tracks were visualized in IGV^187^.

### Single-cell RNA-sequencing of hPSC-derived cell-types

hPSCs and differentiated ectoderm cells were enzymatically dissociated in Accutase (Thermo Fisher) at 37 °C. Day 14 neuron-containing populations were dissociated in papain (Worthington). All samples were run through a 40 μM filter prior to fixation for 24 hours at 4C and storage at -80C. Single cell gene expression was profiled using the Chromium Single Cell Gene Expression Flex kit (10x Genomics) according to the manufacturer’s recommendations. A multiplexed experimental design was used in which 10,000 cells were targeted to be recovered per probe barcode, and 5000 cells per sample were targeted to be captured on Chip Q. Library preparation was confirmed using Agilent Bioanalyzer (High Sensitivity DNA kit), and libraries were sequenced on a NovaSeq 6000.

### Computational analysis of single-cell RNA-sequencing data from hPSC-derived cell-types

Cell Ranger software^188^ (10X Genomics) was used to perform read alignment to the hg38 human reference genome, barcode counting, and unique molecular identifier (UMI) counting. Downstream analysis was conducted using scanpy^189^. Two similar, but distinct, types of single-cell transcriptome quality control criteria were then applied.

First, for days 0-2 of differentiation: we filtered out (1) genes that were expressed in fewer than 3 cells and (2) cells with fewer than 3,000 UMIs, more than 5% mitochondrial reads, more than 100,000 UMIs, and those in which the reference gene *YWHAZ* was not detected.

Second, for all later days of differentiation: we filtered out (1) genes that were expressed in fewer than 3 cells and (2) cells with fewer than 2,000 UMIs, more than 15% mitochondrial reads, and more than 100,000 UMIs.

Subsequently, data were normalized to 10,000 reads per cell and log_e_ transformed. Expression of marker genes was denoted by different color shades and superimposed over the entire cell population as visualized on a Uniform Manifold Approximation and Projection (UMAP) projection^190^. All cells within the entire population that passed the aforementioned quality control criteria were shown on the UMAP projection, without pre-selecting any subset of cells.

The top 25 differentially expressed genes between day-2 anterior neural ectoderm vs. posterior neural ectoderm were determined, based on the fold change in expression between these two cell-types. A threshold of P value < 0.05 was applied to exclusively examine statistically significant differences in expression.

An interactive web browser to explore scRNAseq data of the hPSC-derived cell-types generated in this study is available: https://anglohlabs.shinyapps.io/dundes_jokhai_lohlab/.

### OmniATAC-seq of hPSC-derived cell-types

OmniATAC-seq was performed using a modified version of the original protocol^191^, using modifications described by Klaus Kaestner’s laboratory (University of Pennsylvania). In brief, 50,000 live, freshly isolated cells from each sample were washed with 50 μl cold 1X PBS and pelleted. 50 μl of cold lysis buffer (0.1% v/v NP-40, 0.1% v/v Tween-20, 0.01% v/v Digitonin in resuspension buffer, comprised of 10 mM Tris-HCL, 10 mM NaCl, and 3 mM MgCl_2_) was added to each sample and incubated for 3 minutes on ice. 1 mL of wash buffer (resuspension buffer with 0.1% v/v Tween-20) was added to each sample before centrifugation at 500xg for 10 minutes at 4C. The supernatant was discarded, and each nuclei pellet was resuspended in 50 μl of transposition buffer (25 μl 2X TD Buffer, 16.5 μl 1X PBS, 0.5 μl 10% Tween-20, 0.5 μl 1% Digitonin, 2.5 μl Tn5 Transposase, 5 μl nuclease-free water). The mixture was incubated for 30 minutes at 37°C on a thermomixer set to 1000 rpm.

DNA was isolated using the Qiagen MinElute Reaction Cleanup Kit, as per manufacturer’s instructions. Libraries were generated using Illumina Nextera DNA Unique Dual Indexes and NEBNext High-Fidelity 2X PCR Master Mix. Libraries were double-side purified using AMPure XP beads, and quality was assessed using Agilent Bioanalyzer (High Sensitivity DNA kit). Sequencing was conducted on a NovaSeq 6000.

### Computational analysis of OmniATAC-seq data

Read quality for ATAC-seq data was checked using FastQC and size-distribution and transcriptional start site (TSS) enrichment was confirmed using ATAQV. Reads were trimmed using NGMerge and aligned to human genome hg38 using bowtie2^192^ with the flags --very-sensitive -I 10 -X 2000. Duplicate reads were removed using samblaster, and samtools was used to convert file formats. Peaks were called on individual replicates using MACS2^193^. All peaks across all samples were merged and blacklisted regions were excluded using bedtools to create a set of peaks that define all possible accessible regions across the assayed cell-types.

TMM (Trimmed Mean of M-Values) normalization was performed on each ATAC-seq replicate to accurately account for differences in library composition and sequencing depth. First, featureCounts was used to count reads in accessible regions from each library. EdgeR was then used to calculate size factors for each library, and the final normalization factor was calculated as 1,000,000/(size factor * number of reads in peaks). Normalized bigwig files were generated for individual replicates by using deeptools, with scale factor equal to the final normalization factor that was calculated, and fragments larger than 120 bases were excluded. Replicates were then merged using wiggletools. Tracks were visualized in IGV^187^. Evolutionary conservation of genome sequence across 46 vertebrate species, as determined using Phastcons^194^, was also depicted as an additional track in IGV.

DESeq2^195^ was used to quantify differences in accessibility between cell-types using the count matrix generated from featureCounts. The statistical significance of differential accessibility was determined using the DESeq2 pipeline^195^. In brief, a Wald test was first applied to determine the P value. These P values were then adjusted for multiple hypothesis testing using the Benjamini-Hochberg procedure to controls for the false discovery rate, resulting in Q values.

Replicate similarity was confirmed by visual inspection of TMM normalized bigwigs and by PCA. Peaks were annotated using ChIPseeker, and volcano plots of differential regions between cell types were visualized using EnhancedVolcano.

Enriched motifs were determined using HOMER^196^. The statistical significance of motif enrichment was determined using the Fisher exact test, which is based on the hypergeometric distribution^196^.

### Ca^2+^ live imaging

WTC11 *AAVS1-CAG-GCaMP6f* hiPSCs^182^ (which express the fluorescent Ca^2+^ sensor GCaMP6f^146^) were differentiated into dorsal and ventral hindbrain neurons in 14 days as described above. From days 15-30, neurons were maintained in the same neuron differentiation media described above (Neurobasal media + N2 + B27 + BDNF + GDNF + Forskolin + 2-Phospho-L-ascorbic acid + DAPT + penicillin/streptomycin), and this media was changed every other day. On day 30, cells were imaged on an Olympus FV3000 confocal microscope at room temperature with an air 20X/0.75 objective using a 488 nm laser at 7.9 Hz. Time-lapse movies were imported into FIJI ImageJ, where cells were identified with user-defined ROIs on a maximum intensity projection. A background ROI was also user-defined. For each frame, the pixel value of the background ROI was subtracted from each cellular ROI prior to calculating ΔF/F_0_. A representative subset of ROI traces was selected for plotting.

### Electrophysiology

hPSC-derived neurons were placed in an extracellular Tyrode’s medium (150 mM NaCl, 4 mM KCl, 2 mM CaCl_2_, 2 mM MgCl_2_, 10 mM HEPES [pH 7.3 - 7.4], and 10 mM glucose [osmolarity 335-345]). A borosilicate patch pipette (Harvard Apparatus) with a resistance of 5-6 MΩ was filled with intracellular medium (140 mM potassium-gluconate, 10 mM EGTA, 2 mM MgCl_2_, and 10 mM HEPES [pH 7.2]). Signals were amplified and digitized using the Multiclamp 700B and DigiData1400 (Molecular Devices), and a Leica DM LFSA microscope was used to visualize the cells. Voltage-clamp experiments were conducted at a membrane potential of −70 mV.

For Na^+^ and K^+^ current measurement, cells were held at potentials from -80 mV to +50 mV in steps of 10 mV. Liquid junction potentials (LJPs) were corrected using the Clampex build-in LJP calculator as previously described^197^. Current clamp measurements in neurons were conducted with the current injected to hold the potential at -65 mV to -70 mV, followed by stepwise tonic current injection (from – 100 pA to 350 pA, 1.5 second) to elicit action potentials. Phasic currents (200-500 pA, 5 ms, 10 Hz) were injected to generate action potentials. Analyses of electrophysiological results were performed using ClampFit software (Axon Instruments).

### Optogenetics

At day 21 of hPSC differentiation, ventral hindbrain neurons were transduced with adeno-associated viruses (AAVs) encoding either AAVdj-CAG-bReaChES-TS-EYFP or AAV8-hSyn-bReaChES-mCherry for 7 days. bReaChES encodes a red-shifted excitatory opsin^147^. Both *CAG* and human *SYNAPSIN* (hSYN) promoters have been shown to drive expression in motor neurons^198^. We confirmed that viral transduction was successful, owing to expression of the virally-encoded fluorescent protein (either EYFP or mCherry), as visualized using fluorescent microscopy.

On day 28 of hPSC differentiation (i.e., after 7 days of AAV transduction), optogenetic manipulation was performed. The Spectra X Light engine (Lumencor) served as a light source, and a 560 nm filter was used for orange light stimulation, followed by coupling into a Leica DM LFSA microscope.

Patch pipettes (3–6 MΩ) were pulled using a P2000 micropipette puller (Sutter Instruments). For the voltage clamp characterization, cells were recorded at -70 mV holding potential, and 1.0 mW/mm^2^ light was delivered for 1 second. For the current clamp characterization, neurons were stimulated with 5-ms light pulses of 560 nm light at 1.0 mW/mm^2^ with 5 Hz frequency.

### Analysis of embryonic scRNAseq datasets

Single-cell RNA-sequencing data of gastrulating mouse embryos was described in a previous study (Pijuan-Sala et al., 2019)^169^, and was downloaded from ArrayExpress accession number E-MTAB-6967. E6.75 epiblast cells were identified by filtering for cells with *Elf5* and *Rhox5* log_2_ normalized expression = 0, thereby excluding extraembryonic cells. E7.5 neural ectoderm cells were identified with log_2_ normalized expression of *Sox2* > 0.5, and *Nanog, Tfap2a, Rhox5, Foxj1,* and *Elf5* = 0.

Single-cell RNA-sequencing data of gastrulating non-human primate embryos was described in a previous study (Zhai et al., 2022)^170^, and was downloaded from NCBI GEO accession number GSE193007. Neural ectoderm cells were identified with log_2_ normalized expression of *SOX2* > 1 and, *TFAP2A*, *HAND1*, and *APLNR* = 0.

Single-cell RNA-sequencing data of gastrulating chicken embryos was described in a previous study (Williams et al., 2022)^171^, and was downloaded from NCBI GEO accession number GSE181577. Neural ectoderm cells were identified from the annotated “ectoderm” cluster with log_2_ normalized expression of *Sox2* > 0.

Single-cell RNA-sequencing data of gastrulating zebrafish embryos was described in a previous study (Wagner et al., 2018)^172^, and was downloaded from NCBI GEO accession number GSE112294. Neural ectoderm cells were identified with log_2_ normalized expression of *sox2* > 0.5 and *tfap2a* < 0.5.

### *In situ* hybridization of whole mount hemichordate embryos

DNA probe sets were generated by Molecular Instruments, Inc., targeting the RNA sequences of the *Saccoglossus kowalevskii* genes *otx* (NM_001164888.1) and *gbx* (NM_001164886.1). Twenty probe sets were generated for each gene, and DNA oligo pools were resuspended to a final probe concentration of 1μmol/μl in 50 mM Tris buffer, pH 7.5. HCR amplifiers, B1-Alexa Fluor-546 for *otx* and B2-Alexa Fluor-647 for *gbx* were ordered from Molecular Instruments, Inc.

Hybridization chain reaction v3.0 (HCR3)^176^ was performed as described previously, in combination with the EZ Clear tissue clearing method^199^. Embryos were fixed for 30 minutes at room temperature in fixation buffer (3.7% formaldehyde, 0.1 M MOPS pH 7.5, 0.5 M NaCl, 2 mM EGTA pH 8.0, 1 mM MgCl2, 1X PBS), and subsequently stored in ethanol (EtOH) at - 20°C until use. Embryos were rehydrated from EtOH into RNAse-free water using an EtOH/water series (75%/25%, 50%/50%, 25%/75%, 100% water, 5 minutes each step) and then rinsed twice in water for 10 minutes. The fixed embryos were cleared overnight (12-16 hours) in 50% v/v tetrahydrofuran in water at 4°C, followed by washing three times for 30 minutes each in water, and then twice for 5 minutes each in PBST (1X PBS + 0.1% Triton X-100). Embryos were then incubated in 500 μl of Detergent Solution (1% SDS, 0.5% Tween, 50 mM Tris-HCl, pH 7.5, 1 mM EDTA, pH 8.0, 150 mM NaCl) for 30 minutes. Samples were rinsed twice with PBST and then washed four times for 5 minutes each with PBST, followed by 5 minutes in 5X SSCT (5X sodium chloride sodium citrate, 0.1% Tween 20). Embryos were pre-hybridized in 100 μl of probe hybridization buffer (30% formamide, 5X sodium chloride sodium citrate, 9 mM citric acid, pH 6.0, 0.1% Tween 20, 50 μg/mL heparin, 1X Denhardt’s solution, 10% dextran sulfate) for 1 hour at 37°C before adding 50 μl of probe solution (6 μl of each probe mixture to 50 μl final volume of probe hybridization buffer) and incubated overnight (12-16 hours) at 37°C. Samples were then washed 4 × 30 min with 1 mL of probe wash buffer (30% formamide, 5X sodium chloride sodium citrate, 9 mM citric acid, pH 6.0, 0.1% Tween 20, 50 μg/mL heparin) at 37°C. Samples were further washed twice for 5 minutes each with 1 mL of 5X SSCT at room temperature. Embryos were then pre-amplified with 100 mL of amplification buffer (5X sodium chloride sodium citrate, 0.1% Tween 20, 10% dextran sulfate) for 30 minutes at room temperature before adding the prepared hairpin solution. Hairpin solution was prepared by heating 4 μl of hairpin h1 and 4 μl of hairpin h2 in separate tubes at 95°C for 90 seconds, then cooled to room temperature in the dark for 30 minutes. The h1 and h2 hairpins were combined with 50 μl final volume amplification buffer, and then added to the pre-incubating samples. Samples were incubated in the hairpin solution overnight (12-16 hours) at room temperature in the dark. Hairpin solution was removed by washing with 1 mL of 5X SSCT at room temperature following a series (2 x 5 min, 2 x 30 min, 1 x 5 min). Finally, samples were incubated in NaCl-PBS Hoechst solution (1X PBS, 500 mM NaCl, 1:1000 Hoechst) in the dark at 4°C overnight (12-16 hours) before rinsing in 1X PBS. Embryos were then mounted on slides. The PBS was removed from the embryos while on the slide, and then 30 μl of EZ Clear mounting medium (80% Nycodenz, 7M urea, 0.05% sodium azide, 0.02 M sodium phosphate) was added to clear the embryos. Imaging was conducted on a Zeiss LSM 700 confocal microscope using the Zen software package (Carl Zeiss). For each sample, series of optical sections were acquired with a z-step interval of 6 μm. Optical sections of ectodermal tissues were compiled into a maximum intensity z-projection using FIJI v2.14.0.

### *In situ* hybridization of whole mount zebrafish embryos

Procedures involving zebrafish were approved by the University of California San Francisco IACUC. Adult zebrafish were kept at 28.5 °C in a 14-hour light and 10-hour dark cycle. All zebrafish husbandry was performed in accordance with institutional and national ethical and animal welfare guidelines. Sex/gender was not accounted for as zebrafish sex determination takes place later, during juvenile stages of development.

Zebrafish embryos at 8 hours post-fertilization (hpf) were fixed overnight in 4% paraformaldehyde at 4°C and gradually dehydrated into methanol. Whole mount *in situ* hybridization was performed using hybridization chain reaction v3.0 (HCR3)^176^. Hybridization mRNA probes, amplifiers, and buffers were purchased from Molecular Instruments, and the protocol provided by the manufacturer for whole mount zebrafish embryos was followed with no modifications. Embryos were mounted in ProLong Gold Antifade Mountant (Thermo Fisher, Cat# P10144) and imaged on an upright Stellaris 5 scanning confocal microscope with a Fluotar 25X water immersion lens. Images were visualized using Imaris to generate 3D renderings (Imaris 9.3.1, Oxford Instruments).

### *In situ* hybridization of whole mount chicken embryos

Hybridization chain reaction v3.0 (HCR3)^176^ was performed using the protocol suggested by Molecular Technologies, with minor modifications. Briefly, embryos were fixed in 4% paraformaldehyde (PFA) overnight at 4°C, washed in 0.1% PBS-Tween (PBSTw), dehydrated in a series of 25%, 50%, 75%, and 100% methanol washes, and stored in 100% methanol overnight at -20°C. The following day, embryos were rehydrated and incubated with 10 pmol of *Otx2* (B10 initiator), *Gbx2* (B2 initiator), and *Sox2* (B1 initiator) probes dissolved in hybridization buffer overnight at 37°C. The subsequent day, following several washes in hybridization probe wash buffer and 0.1% 5x-SSC-Tween, embryos were incubated in 30 pmol of amplifier hairpins H1 and H2 labeled with Alexa 488, Alexa 546, and Alexa 647 diluted in amplification buffer at room temperature overnight. The next day, embryos were washed in 0.1% 5x-SSC-Tween and imaged. All probes were designed and produced by Molecular Technologies.

### Statistical analysis

As aforementioned, qPCR histograms depict the mean of two biological replicates, and error bars represent the standard error of the mean (s.e.m.). Sample sizes were based on those typically used in past studies from our laboratory. No datapoints were excluded, apart from qPCR technical replicates that failed. Experiments were repeated adequately to ensure reproducibility, and the number of experiments is generally described either in the figure legends or the **Methods** section. No randomization was performed. Investigators were not blinded. A two-tailed, unpaired t-test was used to test for statistical significance for qPCR data shown in histograms. One asterisk (*) denotes P<0.05, and two asterisks (**) denotes P< .01. For each experiment, we compared all values relative to the following controls:

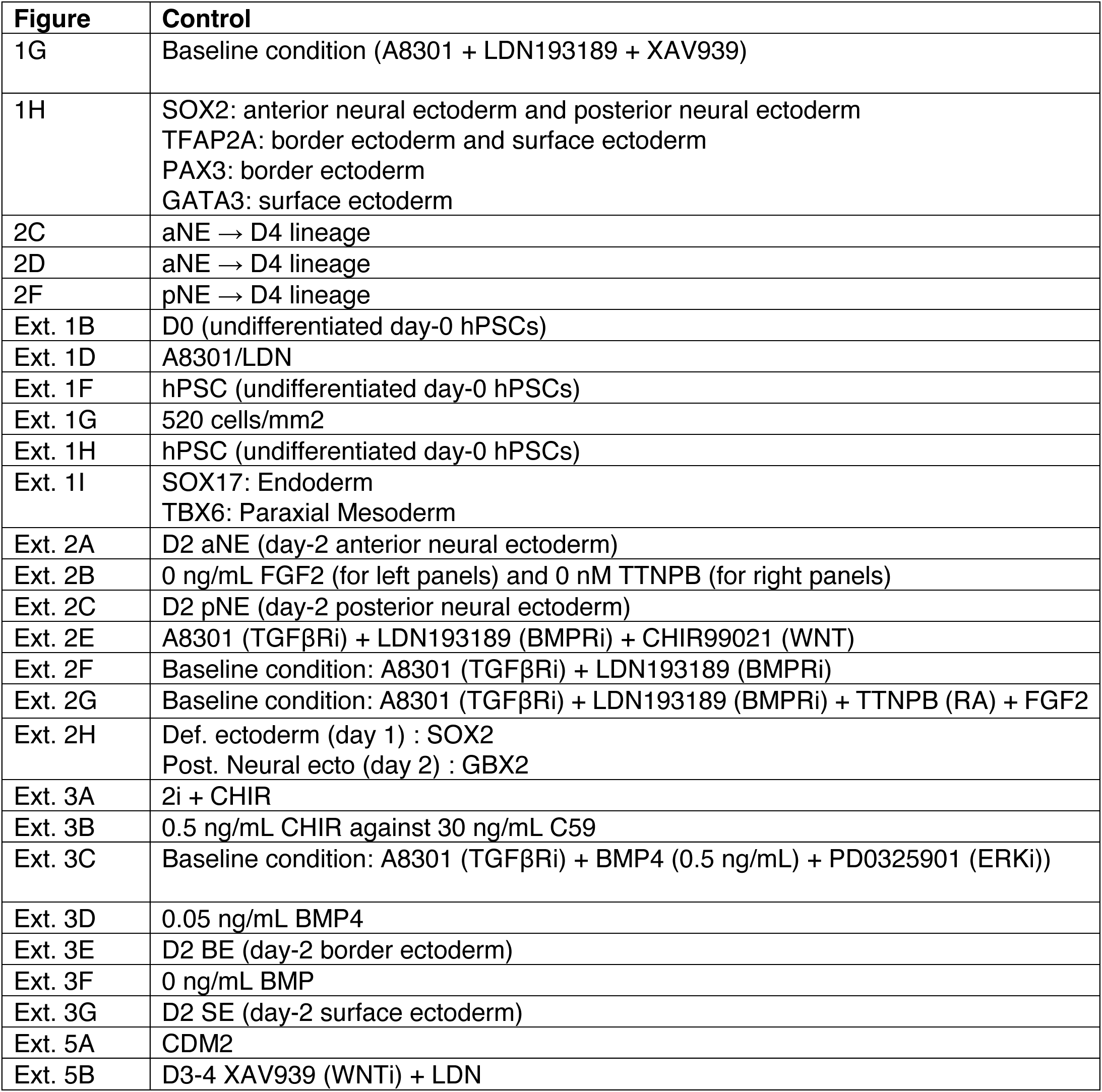
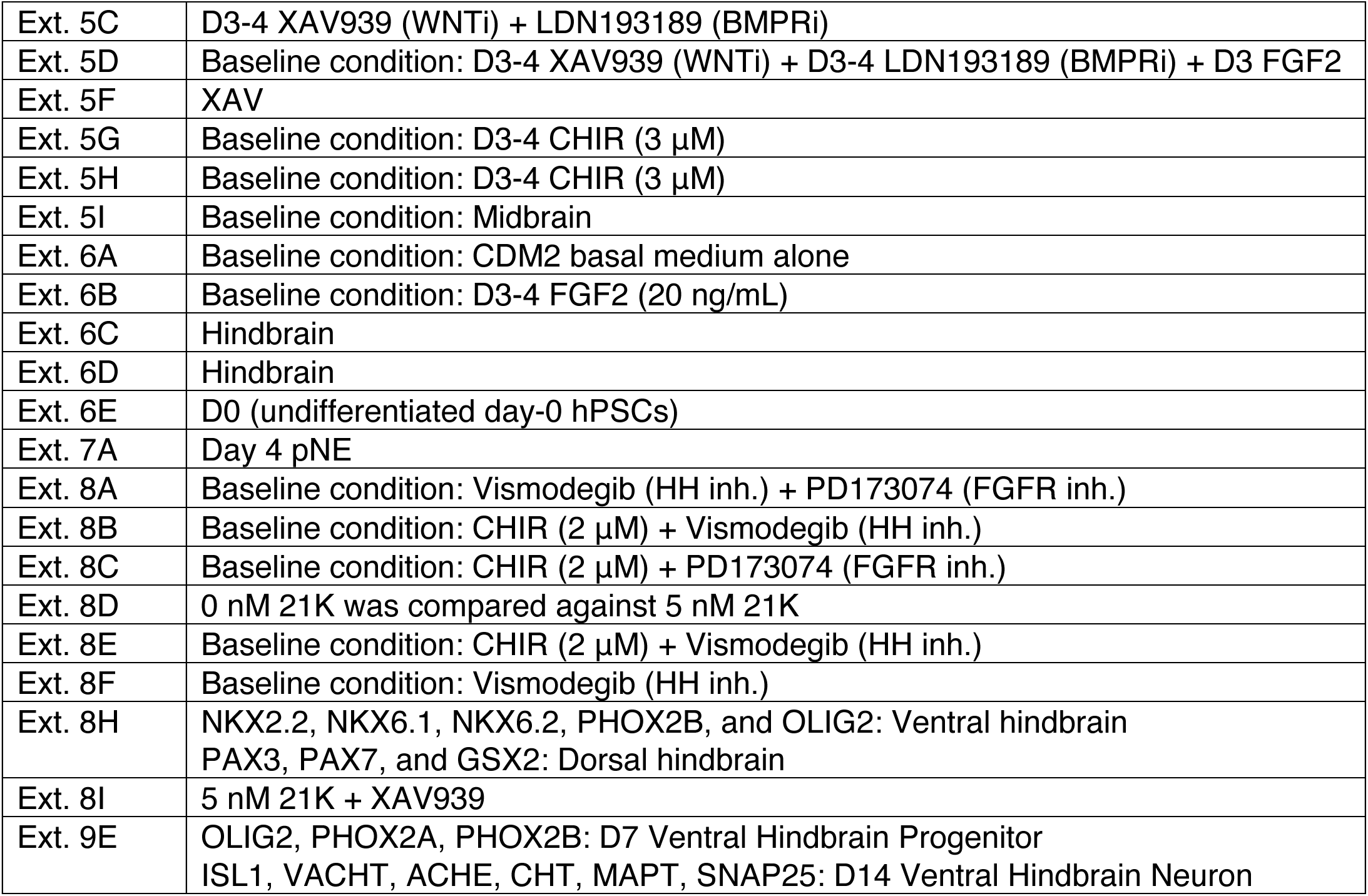

### Data collection

Confocal images were acquired on an Olympus FV3000 confocal microscope using Olympus Fluoview software. Flow cytometry data were acquired on a Beckman Coulter Cytoflex flow cytometer using BD Biosciences FACSDiva software. qPCR data were acquired on a Thermo Fisher QuantStudio 5 qPCR machine using QuantStudio Design and Analysis software. Deep sequencing was performed on a NovaSeq 6000 sequencer using NovaSeq Control Software. Electrophysiological signals were amplified and digitized using the Multiclamp 700B amplifier and DigiData 1440 digitizer (Molecular Devices) followed by data acquisition and recording with pCLAMP 10 software.

### Data analysis

Confocal image analysis was performed using Imaris and ImageJ software. Flow cytometry data analysis was performed using FlowJo software. qPCR data analysis and statistical tests were performed on Microsoft Excel and GraphPad Prism software. Deep sequencing datasets were analyzed in Python, and further details regarding specific computational analyses of sequencing datasets are provided in the respective sections of the **Methods** statement. Electrophysiological data were analyzed using ClampFit software (Axon Instruments).

### Data visualization

Confocal microscope images were plotted using Imaris and ImageJ software. Flow cytometry plots were plotted using FlowJo software. qPCR histograms were plotted using Microsoft Excel. Genomics plots were plotted using Python, with the exception of genomics tracks, which were visualized using IGV^187^. qPCR heatmaps were generated using the online Heatmapper platform^200^ (http://heatmapper.ca/). Animal silhouettes were modified from open-source images, including those from https://www.phylopic.org/. Figures were assembled with Microsoft PowerPoint.

### Ethics and regulatory approval

All hPSC experiments were performed under the regulatory oversight of the Stem Cell Research Oversight (SCRO) committee. All experiments on living mice were performed under the regulatory oversight of the Administrative Panel on Laboratory Animal Care (APLAC) committee.

### Data and materials availability

Bulk-population RNA-sequencing, single-cell RNA-sequencing, and OmniATAC-sequencing datasets generated as part of this study are available at the NCBI Gene Expression Omnibus (GEO): accession numbers GSE286146 (OmniATAC-sequencing), GSE286147 (bulk RNA-sequencing), and GSE286148 (single-cell RNA-sequencing). These datasets are grouped under SuperSet accession number GSE286214. An interactive web browser to explore single-cell RNA-sequencing data of the hPSC-derived cell-types generated in this study is available: https://anglohlabs.shinyapps.io/dundes_jokhai_lohlab/.

In this study, we also analyzed published single-cell RNA-sequencing datasets of gastrulating embryos; these datasets were generated from various species. Single-cell RNA-sequencing data of gastrulating mouse embryos were generated in a previous study (Pijuan-Sala et al., 2019) ^169^, and was downloaded from ArrayExpress accession number E-MTAB-6967. Single-cell RNA-sequencing data of gastrulating non-human primate embryos were generated in a previous study (Zhai et al., 2022) ^170^, and was downloaded from NCBI GEO accession number GSE193007. Single-cell RNA-sequencing data of gastrulating chicken embryos were generated in a previous study (Williams et al., 2022) ^171^, and was downloaded from NCBI GEO accession number GSE181577. Single-cell RNA-sequencing data of gastrulating zebrafish embryos were generated in a previous study (Wagner et al., 2018) ^172^, and was downloaded from NCBI GEO accession number GSE112294.

Computational scripts used in this study to analyze sequencing datasets are available on GitHub: https://github.com/lohlaboratory/ane-pne.

### Experimental reagents

| Reagent or Resource | Source | Identifier |
| --- | --- | --- |
| **Antibodies** |  |  |
| Rabbit anti-TFAP2A (for immunostaining) | Abcam | ab108311, concentration 1:250 |
| Mouse anti-GATA3 (for immunostaining) | Thermo Fisher | MA1-028, concentration 1:1000 |
| Mouse anti-PAX3 (for immunostaining) | R&D Systems | mAB2457, concentration 1:50 |
| Goat anti-SOX2 (for immunostaining) | R&D Systems | AF2018, concentration 1:100 |
| Rabbit anti-RFP (for immunostaining) | Rockland | 600-401-379, concentration 1:500 |
| Goat anti-5-HT (for immunostaining) | Immunostar | 20079, concentration 1:500 |
| Goat anti-OTX2 (for immunostaining) | R&D Systems | AF1979, concentration 1:200 |
| Mouse anti-LHX2 (for immunostaining) | Invitrogen | MA5-15834, concentration 1:2000 |
| Mouse anti-EN1 (for immunostaining) | DSHB | 4611, concentration 1:35 |
| Goat anti-HNF1B (for immunostaining) | R&D Systems | AF3330, concentration 1:250 |
| Rabbit anti-MAFB (for immunostaining) | Sigma | HPA005653, concentration 1:800 |
| Rabbit anti-HOXA3 (for immunostaining) | Novus | NBP1-83234, concentration 1:100 |
| Rabbit anti-SIX3 (for immunostaining) | Rockland | 600-401-A26S, concentration 1:1000 |
| Mouse anti-PAX6 (for immunostaining) | Santa Cruz Biotechnology | sc81649, concentration 1:50 |
| Rabbit anti-NKX2.1 (for immunostaining) | Abcam | ab76013, concentration 1:250 |
| Rabbit anti-OLIG3 (for immunostaining) | Sigma | HA018303, concentration 1:1000 |
| Rabbit anti-OLIG2 (for immunostaining) | Millipore | AB9610, concentration 1:500 |
| Mouse anti-TAU/MAPT (for immunostaining) | Millipore | MAB3420, concentration 1:1000 |
| Rabbit anti-TBR1 (for immunostaining) | Abcam | ab31940, concentration 1:2000 |
| Rabbit anti-DLX1 (for immunostaining) | Thermo Fisher | PA5-28899, concentration 1:1000 |
| Mouse anti-VGLUT2 (for immunostaining) | Thermo Fisher | MA5-31378, concentration 1:200 |
| Rabbit anti-VGAT (for immunostaining) | Synaptic | 131 003, concentration 1:1000 |
| Rabbit anti-CHT/SLC5A7 (for immunostaining) | Abcam | ab56074, concentration 1:100 |
| Mouse anti-ISL1/2 (for immunostaining) | DSHB | 39.4D5, concentration 1:500 |
| Rabbit anti-TLX3 (for immunostaining) | Thermo Fisher | PA5-83146, concentration 1:500 |
| Rabbit anti-NANOG (for immunostaining) | Abcam | AB21624, concentration 1:200 |
| Rabbit anti-GBX2 (for immunostaining) | Proteintech | 21639-1-AP, concentration 1:1000 |
| Mouse anti-β-catenin (for immunostaining) | Abcam | Ab305261, concentration 1:50 |
| Human anti-NANOG-APC (for intracellular flow cytometry) | Miltenyi | 130-105-079, concentration 1:11 |
| Mouse anti-SOX2-PE (for intracellular flow cytometry) | BD Biosciences | 245610, concentration 1:5 |
| Rabbit anti-Gbx2 (for Western Blot) | Thermo Fisher | PA5-66953, concentration 1:1000 |
| Rabbit anti-Vinculin (for Western Blot) | Cell Signaling Technology | E1E9V, concentration 1:1000 |
| Donkey anti-goat Alexa 647 | Thermo Fisher | A21447, concentration 1:500 |
| Donkey anti-guinea pig Alexa 488 | Jackson ImmunoResearch | 706-545-148, concentration 1:500 |
| Donkey anti-mouse Alexa 555 | Thermo Fisher | A32773, concentration 1:500 |
| Donkey anti-mouse Alexa 647 | Thermo Fisher | A31571, concentration 1:500 |
| Donkey anti-rabbit Alexa 555 | Thermo Fisher | A-31572, concentration 1:500 |
| Donkey anti-rabbit Alexa 647 | Jackson ImmunoResearch | 711-607-003, concentration 1:500 |
| Donkey anti-goat Alexa 488 | Thermo Fisher | A32814, concentration 1:500 |
| **Chemicals, peptides, and recombinant proteins** |  |  |
| mTesR Plus medium | STEMCELL Technologies | #100-0276 |
| Penicillin/streptomycin | Thermo Fisher Scientific | 15-140-122 |
| Geltrex LDEV-Free, hESC-Qualified, Reduced Growth Factor Basement Membrane Matrix | Thermo Fisher Scientific | A1413302 |
| Versene solution | Thermo Fisher Scientific | 15040066 |
| Accutase - Enzyme Cell Detachment Medium | Thermo Fisher Scientific | 00-4555-56 |
| TrypLE Express Enzyme (1X) | Thermo Fisher Scientific | 12604013 |
| Fetal Bovine Serum Premium Select | Atlanta Biologicals | S11550H |
| 2-mercaptoethanol | Gibco | 21985023 |
| DMEM/F12 + GlutaMAX | Gibco | 10565042 |
| IMDM + GlutaMAX | Thermo Fisher | 31980-097 |
| F12 + GlutaMAX | Thermo Fisher | 31765-092 |
| Polyvinyl alcohol | Sigma | P8136-250G |
| Chemically defined lipid concentrate | Thermo Fisher | 11905-031 |
| 1-thioglycerol | Sigma | M6145-100ML |
| Recombinant human insulin | Sigma | 11376497001 |
| Recombinant human VEGF | R&D Systems | 293-VE-0500 |
| Recombinant human FGF2 | R&D Systems | 233-FB-01M |
| Human transferrin | Sigma | 10652202001 |
| Recombinant human BMP4 | R&D Systems | 314-BP-050 |
| Penicillin/streptomycin | Thermo Fisher | 15070-063 |
| Recombinant Activin | R&D Systems | 338-AC-500/CF |
| GDC-0941 | Cellagen Technology | C4321-25 |
| Forskolin | Tocris | 1099 |
| Thiazovivin | Tocris | 3845 |
| XAV939 | Tocris | 3748 |
| Ascorbic acid-2-phosphate | Sigma | 49752-10G |
| DMH1 | Tocris | 4126 |
| CHIR99201 | Tocris | 4423 |
| PD0325901 | Tocris | 4192 |
| Bovine serum albumin (BSA) | Sigma | A2153 |
| Neurobasal medium | Thermo Fisher Scientific | 21103049 |
| B27 | Thermo Fisher Scientific | 17504044 |
| N2 | Thermo Fisher Scientific | 17502001 |
| NGS WNT-Fc | K. Christopher Garcia's laboratory | 201 |
| RSPO2/R-spondin 2 | K. Christopher Garcia's laboratory | 201 |
| Recombinant human WNT3A | R&D Systems | 5036-WN-010 |
| **Critical commercial assays** |  |  |
| RNeasy Plus Mini Kit | Qiagen | 10010049 |
| RNeasy Micro kit | Qiagen | 74004 |
| High-Capacity cDNA Reverse Transcription Kit | Applied Biosystems | 4368814 |
| SYBRGreen | Thomas Scientific | BIO-94050 |
| 10x Genomics Chromium single cell 3' GEM reagent kit (v3) | 10x Genomics | 1000075 |
| Triton X-100 | Sigma | X100-500ML |
| PBS, pH 7.4 | Thermo Fisher | 10010049 |
| Molecular Probes DAPI (4',6 Diamidino 2 Phenylindole, Dihydrochloride) | Thermo Fisher Scientific | D1306 |
| **Experimental models: cell lines** |  |  |
| H1 hESCs | WiCell | WiCell, WA01 |
| H7 hESCs | WiCell | WiCell, WA07 |
| H9 hESCs | WiCell | WiCell, WA09 |
| SUN004.1 hiPSCs | Hiromitsu Nakauchi's laboratory | 178 |
| SUN004.2 hiPSCs | Hiromitsu Nakauchi's laboratory | 178 |
| HES3 *MIXL1-GFP* hPSCs | Andrew Elefanty's, Edouard Stanley's, and Elizabeth Ng's laboratories | 181 |
| WTC *AAVS1-CAG-GCaMP6f* hiPSCs | Bruce Conklin's laboratory | 182 |
| **Software and algorithms** |  |  |
| ImageJ/Fiji | 177 | https://imagej.net/software/fiji/ |
| FastQC | 183 | http://www.bioinformatics.babraham.ac.uk/projects/fastqc/ |
| Trim Galore | 202 | http://www.bioinformatics.babraham.ac.uk/projects/trim_galore/ |
| Bowtie2 | 192 |  |
| MACS2 | 193 |  |
| DESeq2 | 195 |  |
| HOMER | 196 |  |

